## Supplementary Materials for "A Quantitative Genetic Model of Background Selection in Humans"

September 8, 2023

1.9 The Fitness-Effective Population Size Under Weak and Strong Negative Selection . . 16

**2 Statistical Methods** **18**

### 1 Quantitative Genetic Linked Selection Model Theory

#### 1.1 Stochastic Sources of Neutral Allele Frequency Change

Here, we step through a quick derivation of Santiago and Caballero (1995). Throughout, we assume random mating, hermaphroditic individuals, and a constant population size. The change in a neutral allele's frequency in one generation can be partitioned into the three sources of stochasticity: the random associations with fitness backgrounds, the non-heritable randomness in family size, and the Mendelian noise from heterozygotes segregating. If we let $x_{0,i} \in \{0, 1/2, 1\}$ be the frequency of neutral alleles individual $i$ in generation 0 carries, we can partition the random neutral allele frequency of the population ($x_t$ without the individual index) into the underlying stochastic causes,

$$x_1 = \frac{1}{2N} \sum_{i=1}^{N} \left( x_{0,i} k_{0,i} + \sum_{j=1}^{k_{0,i}} \delta_{0,i,j} \right) \tag{1}$$

where $k_{0,i}$ is the number of surviving gametes parent $i$ passes on, and $\delta_{0,i,j}$ is a random term that encapsulates the noise due to Mendelian segregation of heterozygotes. If parent $i$ is a homozygote ($x_{0,i} \in \{0, 1\}$), then $\delta_{0,i,j} = 0$, whereas if $x_{0,i} = 1/2$ then $\delta_{0,i,j} = \pm 1/2$ with equal probability. This is because a heterozygous parent will transmit half a neutral allele in expectation, but each round of Mendelian segregation must pass on either 0 or 1 alleles, which the random $\pm 1/2$ term imposes. The factor of $1/2$ is due to the fact that we're summing over $N$ diploids, but considering the number of gametes they transmit. Since each diploid parent must have two offspring to maintain a constant population size, $1/N \sum_i k_{0,i} = 2$.

The frequency in the initial generation is $x_0 = 1/N \sum_{i=1}^{N} x_{0,i}$, though to indicate that we treat this as fixed rather than random, we use $p_0 := x_0$. Then, the allele frequency change is,

$$\begin{aligned} \Delta x_1 = x_1 - p_0 &= \frac{1}{2N} \sum_{i=1}^{N} x_{0,i}(k_{0,i} - 1) + \frac{1}{2N} \sum_{i=1}^{N} \sum_{j=1}^{k_{0,i}} \delta_{0,i,j} \\ \Delta x_1 &= K_1 + H_1 \end{aligned} \tag{2}$$

where $K_1$ and $H_1$ are the random change in neutral allele frequency change due to offspring number (including heritable and non-heritable components), and Mendelian segregation in generation 1.

#### 1.2 The Variance in Neutral Allele Frequency Change

Now, let us look the variance of $\mathrm{V}(\Delta x_1)$ over evolutionary replicates. Since the Mendelian segregation and the offspring random process are independent,

$$\mathrm{V}(\Delta x_1) = \mathrm{V}(K_1) + \mathrm{V}(H_1). \tag{3}$$

Looking at each term,

$$\mathrm{V}(K_1) \approx \frac{1}{4N^2} \sum_{i=1}^{N} \mathrm{V}\left(x_{0,i}(k_{0,i} - 1)\right) \tag{4}$$

where we ignore the covariance terms due to the sum, since these are on order $1/N^3$. In the first generation from an arbitrary starting point, the neutral alleles assort independently into diploids with respect to their fitness, so we can simply the variance as

$$\begin{aligned} \mathrm{V}(K_1) &\approx \frac{1}{4N^2} \sum_{i=1}^{N} \mathrm{V}(x_{0,i})\mathrm{V}(k_{0,i} - 1) \\ &\approx \frac{1}{4N}\mathrm{V}(x_{0,i})\mathrm{V}(k_{0,i}). \end{aligned} \tag{5}$$

Assuming no correlation between parental gametes (e.g. no inbreeding), $\mathrm{V}(x_{0,i})$ is the binomial variance in individual allele frequency, or $p_0(1 - p_0)/2$, and $\mathrm{V}(k) := \mathrm{V}(k_{0,i})$ is the offspring variance of an individual given by the reproduction process. For example, if the reproduction process is a neutral multinomial Wright–Fisher, $\mathrm{V}(k) \approx 2$. Then, the variance in allele frequency change due to non-heritable offspring number variation is,

$$\mathrm{V}(K_1) \approx \frac{p_0(1 - p_0)}{2N} \frac{\mathrm{V}(k)}{4}. \tag{6}$$

The Mendelian noise variance term can be derived similarly. First, the sum over each parent's transmitted gametes can be simplified by noting that parents are exchangeable over evolutionary replicates with respect to their contribution to this term. In a constant population size, the double summation over parents and their offspring can be replaced by a summation over offspring, since both sum $N$ exchangeable terms. Then, note that $\mathrm{V}(\delta_{i,j}) = 1/4 p_0(1 - p_0)$, so

$$\mathrm{V}(H_1) = \frac{p_0(1 - p_0)}{2N} \frac{1}{2}. \tag{7}$$

Finally, we have

$$\begin{aligned} \mathrm{V}(\Delta x_1) &= \mathrm{V}(K_1) + \mathrm{V}(H_1) \qquad (8)\\ &\approx \frac{p_0(1-p_0)}{2N}\frac{\mathrm{V}(k)}{4} + \frac{p_0(1-p_0)}{2N}\frac{1}{2} \\ &\approx \frac{p_0(1-p_0)}{2N}\left(\frac{\mathrm{V}(k)}{4} + \frac{1}{2}\right) \qquad (9) \end{aligned}$$

(c.f. Santiago and Caballero 1995 equation 2 and Buffalo and Coop 2019 equation 30). Note that if the reproduction process is a neutral multinomial Wright–Fisher, $\mathrm{V}(k) \approx 2$, and this simplifies to the expected Wright–Fisher variance. If we were to define a variance *drift-effective* population size $N_e$ by setting this variance in neutral allele frequency change to the expected variance under a Wright–Fisher model ($V_{\mathrm{WF}}$), we'd have

$$\begin{aligned} V_{\mathrm{WF}} &:= \frac{p_0(1-p_0)}{2N_e} \\ V_{\mathrm{WF}} &= \mathrm{V}(\Delta p_1) \\ N_e &= \frac{4N}{\mathrm{V}(k)+2} \qquad (10) \end{aligned}$$

(c.f. Wright 1938).

Now, we look at the variance in allele frequency change across two generations for a neutral allele. The change in neutral allele frequency, regardless of the selective system, is directionless in expectation. This is because whichever allele we track is arbitrary by the definition of neutrality, which imposes a symmetry, $\mathbb{E}(\Delta x) = 0$. Then, the variance of change in the first generation is,

$$\mathrm{V}(\Delta x_1) = \mathbb{E}_1\left[(x_1 - x_0)^2\right] = \frac{x_0(1-x_0)}{2N}, \qquad (11)$$

and across two generations,

$$
\begin{aligned}
\mathrm{V}(x_2 - x_0) &= \mathbb{E}\left[(x_2 - x_0)^2\right] \\
&= \mathbb{E}_1\left[\mathbb{E}_2\left[((x_2 - x_1) + (x_1 - x_0))^2 | x_1\right]\right] \\
&= \mathbb{E}_1\left[\mathbb{E}_2\left[(\Delta x_2 + \Delta x_1)^2 | x_1\right]\right] \\
&= \mathbb{E}_1\left[\mathbb{E}_2\left[\Delta {x_2}^2 | x_1\right]\right] + \mathbb{E}_1\left[\Delta {x_1}^2\right] + 2\mathbb{E}_1\left[\mathbb{E}_2[\Delta x_2 \Delta x_1 | x_1]\right] \\
&= \mathbb{E}_1\left[\mathbb{E}_2\left[\Delta {x_2}^2 | x_1\right]\right] + \mathbb{E}_1\left[\Delta {x_1}^2\right] + 2\mathcal{C}_{1,2} \\
&= \frac{p_0(1-p_0)}{2N}\left(1 - \frac{1}{2N}\right) + \frac{p_0(1-p_0)}{2N} + 2\mathcal{C}_{1,2}
\end{aligned} \tag{12}
$$

Consequently, the variance of the neutral allele frequency changes each generation according to the probability of failing to coalescence each generation and the pairwise covariance terms $\mathcal{C}_{i,j}$ that build up due to associations with heritable fitness backgrounds. Note that the covariance term $\mathcal{C}_{1,2} = \mathbb{E}_1\left[\mathbb{E}_2[\Delta x_2 \Delta x_1 | x_1]\right] = 0$ when there is no heritable fitness variation, and this simplifies to the well-known equation for variance in a Wright–Fisher population.

#### 1.3 Heritable Fitness and the Accumulation of Autocovariance

Next, we turn our attention to the case where there is heritable fitness variation, which leads to non-zero covariance terms $\mathcal{C}_{i,j}$. These terms emerge when $\mathrm{V}(k)$ has a heritable component of fitness that can be transmitted along with the neutral allele, thereby affecting the neutral allele's trajectory in later generations. If we partition the offspring number into heritable and non-heritable components, $k_i = 2f_i + \varepsilon_i$, then $\mathrm{V}(k_i) = 4V_A + V_n$. When there is heritable fitness variation across individuals, $V_A > 0$. Because the population size is assumed to be constant, the mean heritable fitness across individuals is constrained to be $\mathbb{E}_i(f_i) = 1$; this implies that the individual fitness values $f_i$ are relative fitnesses as is standard in population genetics, in this case to the population mean. Thus $V_A$ is the squared coefficient of heritable fitness variation (this is often denoted as $C^2 = V_A/\bar{w}^2$, see Charlesworth 1987; Crow 1958; Houle 1992).

We can further decompose the $K_1$ term into $K_1 = S_1 + D_1$,

$$
\begin{aligned}
\mathrm{V}(\Delta x_1) &= \mathrm{V}(S_1) + \mathrm{V}(D_1) + \mathrm{V}(H_1) \\
&\approx \frac{p_0(1-p_0)}{2N} V_A + \frac{p_0(1-p_0)}{2N}\frac{V_n}{4} + \frac{p_0(1-p_0)}{2N}\frac{1}{2}
\end{aligned} \tag{13}
$$

(c.f. Santiago and Caballero 1995 equation 11).

To see how the covariance terms accumulate, consider $\mathrm{V}(x_3 - p_0)$. Note that $\mathbb{E}(S_t) = \mathbb{E}(D_t) = \mathbb{E}(H_3) = 0$, since which neutral allele we track is arbitrary, so by symmetry the expected change is zero. Then,

$$
\begin{aligned}
\mathrm{V}(x_3 - p_0) &= \mathbb{E}\left[(S_1 + D_1 + H_1 + S_2 + D_2 + H_2 + S_3 + D_3 + H_3)^2\right] \\
&= \mathbb{E}({S_1}^2) + \mathbb{E}({S_2}^2) + \mathbb{E}({S_3}^2) \\
&+ \mathbb{E}(S_1 S_2) + \mathbb{E}(S_1 S_3) + \mathbb{E}(S_2 S_3) \\
&+ \mathbb{E}({D_1}^2) + \mathbb{E}({H_1}^2) + \mathbb{E}({D_2}^2) + \mathbb{E}({H_2}^2) + \mathbb{E}({D_3}^2) + \mathbb{E}({H_3}^2). \qquad (14)
\end{aligned}
$$

The covariance terms $\mathbb{E}(S_i S_j)$ for $j > i$ represent the expected neutral allele frequency change from a neutral allele becoming associated with a fitness background in generation $i$ and that fitness association persisting until generation $j$. However, the covariances terms $\mathbb{E}(S_j S_i)$ for $j > i$ are zero since associations in the future cannot affect the past (see p. 1041 of Buffalo and Coop 2019). Let us consider $S_1$ and $S_2$. We have,

$$
\begin{aligned}
S_1 &= \frac{1}{2N} \sum_{i=1}^{N} x_{0,i}(f_{0,i} - 1) = \mathrm{cov}(x_{0,i}, f_{0,i}) \\
S_2 &= \frac{1}{2N} \sum_{i=1}^{N} x_{1,i}(f_{1,i} - 1) = \mathrm{cov}(x_{1,i}, f_{0,i}) \qquad (15)
\end{aligned}
$$

which are chance covariances (across the population, *not* evolutionary replicates) created by the random sorting of neutral alleles and fitness into individuals, *each generation*. These covariances are equivalent to the Robertson-Price equation (Price 1970; Robertson 1966), which predict the change in neutral frequency due to heritable fitness (and likewise with the change due to non-heritable factors).

Considering the underlying haplotypes that lead to $x_{0,i}$ and $f_{0,i}$ can give us an equation for the dynamics of the $\mathbb{E}(S_i S_j)$ (for $j > i$) terms over evolutionary replicates. Let us partition the allele frequency and fitness by into the average of the paternal contributions. The neutral allele frequency per gamete is either 0 or 1, and we assume the fitnesses across gametes are additive. Then, as long as there is random mating, we can simplify the associations in a diploid by looking at a single gamete,

$$\begin{aligned}
S_1 = \text{cov}(x_{0,i}, f_{0,i}) &= \text{cov}\left(\frac{x'_{0,i} + x''_{0,i}}{2}, f'_{0,i} + f''_{0,i}\right) \\
&= \frac{1}{2}\left(\text{cov}(x'_{0,i}, f'_{0,i} + f''_{0,i}) + \text{cov}(x''_{0,i}, f'_{0,i} + f''_{0,i})\right) \\
&= \text{cov}(x'_{0,i}, f'_{0,i} + f''_{0,i}) \\
&= \text{cov}(x'_{0,i}, f'_{0,i}) + \text{cov}(x'_{0,i}, f''_{0,i}) \\
&= S'_1 + S''_1
\end{aligned} \tag{16}$$

which follows from symmetry again, since it does not matter which gamete carrying the neutral allele we track (c.f. Santiago and Caballero 1998 p. 2107). The primes now indicate whether the selected locus is on the same gamete as the neutral allele (single prime), or the homologous gamete (double prime).

After meiosis, a fraction $1 - r$ of these associations between the tracked neutral neutral allele $x'_{0,1}$ and fitness background $f'_{0,i}$ persist, and a fraction $r$ disassociate its currently coupled background and create an association with the homologous fitness background, $f''_{0,i}$. When linkage is tight, this latter term can be ignored as we will do here, though it does impact background levels of neutral diversity (Santiago and Caballero 1995). Additionally, the fitness effects of the selected background may change in later generations, in complex ways (Barton 1986; Turelli and Barton 1990). Santiago and Caballero (1995) assume an equilibrium level of fitness variation $V_A$, with the fitness variance associated with a particular background decaying at a simple geometric decay at rate $1 - \kappa$ each generation, which is sufficient for background selection. An approximate rate of the decay can be worked out for the particular selective system. In general, this can be a much more complicated function; Buffalo and Coop (2019) show that even fluctuating selection can be accommodated.

Now, the associations created in generation 1 and that persist in future generations as (ignoring $S''_i$ terms) follow the pattern

$$\begin{aligned}
S'_2 &= S_1(1-\kappa)(1-r) \\
S'_3 &= S_1(1-\kappa)^2(1-r)^2 \\
S'_4 &= S_1(1-\kappa)^3(1-r)^3.
\end{aligned} \tag{17}$$

Then, the cumulative effect of the fitness associations created in generation 1 can be written as,

$$S_1 Q_t = S_1 + S_2' + S_3' + \ldots + S_t'$$
$$Q_t = 1 + \sum_{i=1}^{t} (1-r)^i (1-\kappa)^i \tag{18}$$

or for continuous $t$,

$$Q_t = 1 + \frac{(1-\kappa)(1-r)\left(1-(1-\kappa)^t(1-r)^t\right)}{\kappa + (1-\kappa)r} \tag{19}$$

which as $t \to \infty$, converges to,

$$Q_\infty = \frac{1}{\kappa + r(1-\kappa)}. \tag{20}$$

This shows the long reach of heritable fitness factors in altering allele frequency through the generations. In terms like $\mathbb{E}(S_1 S_2)$, a fraction $S_1'$ of the heritable allele frequency change $S_2$ were *previously* built associations between the neutral allele and a fitness background from $S_1$. As long as the direction of selection is the same, the expected change of this fraction of associations formed in the first generation to later generations 2 and 3 would be,

$$\mathbb{E}(S_1 S_2) = \mathbb{E}(S_1^2)(1-\kappa)(1-r) = \mathbb{E}(S_1^2) Q_2^2$$
$$\mathbb{E}(S_1 S_3) = \mathbb{E}(S_1^2)(1-\kappa)^2(1-r)^2 = \mathbb{E}(S_1^2) Q_3^2. \tag{21}$$

The $Q_t^2$ terms represent the inflation of variance due to autocovariance in frequency change across generations due to linked polygenic selection. Incorporating the impact of time-invariant autocorrelation created by linked selection as a rescaled rate of drift is analogous to using the Green–Kubo relation to find the transport coefficient in molecular dynamics with velocity autocorrelation (Green 1954; Kubo 1957).

With these covariance terms, we now have a full model for $\mathrm{V}(p_t - p_0)$. We will look at the case for $t = 2$. As shown in Supplementary Equation (13), the second moment of every term is a function of the variance in neutral allele frequency in that generation, $x_t(1-x_t)$. This variance does not stay its initial value of $x_0(1-x_0)$; it decays each generation due to coalescence from the drift and selection processes modeled by this approach. The rate at which this decay happens is the effective population size each generation implied by this model. We can write,

$$
\begin{aligned}
\mathrm{V}(x_2 - p_0) &= \mathbb{E}({S_1}^2) + \mathbb{E}(S_1 S_2) + \mathbb{E}({S_2}^2) \\
&\quad + \mathbb{E}({D_1}^2) + \mathbb{E}({H_1}^2) + \mathbb{E}({D_2}^2) + \mathbb{E}({H_2}^2) \\
\frac{\mathrm{V}(x_2 - p_0)}{p_0(1 - p_0)} &= \frac{V_A}{2N} + \frac{V_A Q_2^2}{2N}\left(1 - \frac{1}{2N}\right) + \frac{V_A}{2N}\left(1 - \frac{1}{2N}\right) \\
&\quad + \frac{V_n}{8N} + \frac{1}{4N} + \frac{V_n}{8N}\left(1 - \frac{1}{2N}\right) + \frac{1}{4N}\left(1 - \frac{1}{2N}\right). \qquad (22)
\end{aligned}
$$

In general, the variance in allele frequency change after $t$ generations in a system with heritable fitness due to a single locus $r$ recombination fraction away from the neutral site is given by,

$$
\frac{\mathrm{V}(x_t - p_0)}{p_0(1 - p_0)} = \sum_{i=1}^{t} \frac{1}{2N}\left(V_A Q_i^2 + \frac{V_n}{4} + \frac{1}{2}\right)\left(1 - \frac{1}{2N}\right)^{i-1}. \qquad (23)
$$

We will see later that with the appropriate form for $Q_t^2$, this equation holds for polygenic selection (see Supplementary Section 1.5). Additionally, we note that since the probability of coalescence (or, equivalently, identity by descent) is proportional to the variance in allele frequency change (Barton 2000), this equation for the variance encodes the pairwise coalescent rate of the population through time under weak polygenic linked selection. Others have found that the genealogies under purifying selection can be characterized by such a time-dependent effective population size (Nicolaisen and Desai 2013).

#### 1.4 The Fitness-Effective Population Size

Now, we consider how to turn this expression for the variance in neutral allele frequency at time $t$ under weak linked selection into an expression for the *fitness-effective* population size analogous to drift-effective population size given in Supplementary Equation (10). In an ideal Wright–Fisher population, the instantaneous variance is $\mathrm{V}(\Delta x_t) = p_{t-1}(1-p_{t-1})/2N_{d,t-1}$. Santiago and Caballero consider the allele frequency change at some $t$ to define their effective population size at time $t$ (which we here are calling fitness-effective population size, $N_{d,t}$),

$$
N_{d,t} := \frac{p_{t-1}(1 - p_{t-1})}{2\mathrm{V}(x_t) - \mathrm{V}(x_{t-1})}. \qquad (24)
$$

The time-dependency is due to $Q_i^2$, which reflects the fact that autocovariances build up, but at a decreasing rate. Note that levels of pairwise diversity depend on the full distribution of pairwise coalescence times, or the $N_{d,t}$ (e.g. see equation 4 of Santiago and Caballero 2016 and equation 16 of Santiago and Caballero 1998).

Using Supplementary Equation (23) we have,

$$N_{d,t} \approx \frac{4N}{4V_A Q_t^2 + V_n + 2} \tag{25}$$

(cf. equation 18, Santiago and Caballero 1995). Note if $V_A = 0$, this equation reduces to Wright's (1938) equation.

In many cases, we care about the asymptotic $N_f$,

$$N_f := \lim_{t\to\infty} N_{d,t} \approx \frac{4N}{4V_A Q_\infty^2 + V_n + 2}. \tag{26}$$

Now, we turn to different expressions for $Q_t^2$ and $V_A$ under different types of polygenic selection.

#### 1.5 The Fitness-Effective Population Size Under Polygenic Selection

The results of the previous section assume only a single locus a recombination fraction $r$ apart from the neutral locus is determining the coalescence rate. The model of Santiago and Caballero (1998) extends this to the case where fitness is polygenic. As they show in their original paper, this system can accommodate a variety of different equilibrium selection systems as long as expressions for $V_A$ and $1-\kappa$ can be worked out. Other authors have extended similar models to more elaborate breeding structures (Woolliams et al. 1993; Wray and Thompson 1990).

First, we extend Supplementary Equation (25) to a polygenic system. Throughout, we assume a multiplicative fitness model (i.e. independent effects, no epistasis), where the fitness of individual $i$ is the product of their fitness contributions from each locus $l$, $w_{i,l}$, giving $w_i = \prod_{l=1}^{n} w_{i,l}$. Then, assuming mean fitness is one and independence between sites, the total fitness variation is multiplicative across the locus-level fitnesses,

$$V_A = \prod_{l=1}^{n}(1 + V_{h,l}) - 1 \tag{27}$$

Similarly, the $Q_t^2 V_A$ term in the single-locus model is multiplicative under the polygenic model, as the change in site-specific fitness variances are assumed independent,

$$Q_t^2 V_A = \prod_{l=1}^{n}\left(1 + \frac{V_{h,l} Q_{t,l}^2}{2}\right) - 1 \tag{28}$$

where $Q_{t,l}$ is the cumulative impact of selection due to the selected site $l$ (c.f. equation 4 Santiago and Caballero 1998). The factor of $1/2$ is due to the fact that we are ignoring the chance that the fitness background on the homologous chromosome recombines onto the haplotype with the tracked neutral allele (i.e. due to the $Q_t''$ associations). In other words, only half the fitness variation in a diploid can stay associated with the neutral allele through the generations.

To simplify the derivation, assume $V_n = 2$ as under a Wright–Fisher model. Then, the effective population size is,

$$\begin{aligned} N_{d,t} &\approx \frac{N}{V_A Q_t^2 + 1} \\ &\approx \frac{N}{\prod_{l=1}^{n}\left(1 + \frac{V_{h,l} Q_{t,l}^2}{2}\right)} \\ &\approx N \exp\left(-\sum_{l=1}^{n} \frac{V_{h,l} Q_{t,l}^2}{2}\right) \end{aligned} \tag{29}$$

(c.f. Santiago and Caballero 1998 equation 4b). If we write each site $l$'s contribution to the variance as a deviation from the average contribution, $V_{h,l}^2 = 1/n V_A + \varepsilon_l$, then

$$\begin{aligned} N_{d,t} &\approx N \exp\left(-\frac{1}{2}\sum_{l=1}^{n}\left(\frac{1}{n}V_h + \varepsilon_l\right) Q_{t,l}^2\right) \\ &\approx N \exp\left(-\frac{V_h}{2n}\sum_{l=1}^{n} Q_{t,l}^2 + \sum_{l=1}^{n} \varepsilon_l Q_{t,l}^2\right) \end{aligned} \tag{30}$$

Santiago and Caballero assume that over evolutionary replicates, as selected sites are randomly distributed, there is not systematic covariance between the fitness variation at a site and the decay of its association with the neutral site, e.g. $\mathbb{E}(\varepsilon_l Q_{t,l}^2) = 0$, and thus ignore the second sum term, leading to

$$N_{d,t} \approx N \exp\left(-\frac{V_h}{2n}\sum_{l=1}^{n} Q_{t,l}^2\right). \tag{31}$$

##### 1.6 Continuous Approximation to the Segment Under Selection

Our model differs from that of Santiago and Caballero (1998) in that we do not integrate over the entire genome. In their 1998 model, Santiago and Caballero consider the impact of both sites under selection on the same chromosome, as well as sites on independently assorting chromosomes. We only consider the contribution of a single segment under purifying selection, and the reduction from additional segments accumulate multiplicatively. Like their model, we imagine the reduction experienced by a focal neutral site in the middle of the segment.

The $Q_{t,l}$ terms depend on $l$ only through the recombination fraction $r(l)$ between the neutral allele and the selected site. Assuming a constant per-basepair recombination rate of $r_{\mathrm{BP}}$, $r(l) = r_{\mathrm{BP}} l$ we can approximate the sum with an integral for the asymptotic $Q_\infty$. The reduction experienced in the middle of a chromosome is twice the reduction experienced from each half,

$$\begin{aligned} N_{d,\infty} &\approx N \exp\left(-\frac{V_h}{2L}\sum_{l=1}^{L} Q_{\infty,l}^2\right) \\ &\approx N \exp\left(-\frac{V_h}{M}\int_0^{M/2} Q_\infty(r)^2 dr\right) \end{aligned} \tag{32}$$

where $M = nr_{\mathrm{BP}}$ is the total segment length in Morgans. Then, the total asymptotic effect of associations from all sites as $Q_\infty^2$. Then,

$$\begin{aligned} Q_\infty^2 &= \frac{2}{M}\int_0^{M/2} Q_\infty(r)^2 dr \\ &= \frac{2}{M}\int_0^{M/2}\left(\frac{1}{1-(1-r)Z}\right)^2 dr. \\ &= \frac{2}{(1-Z)(2-(2-M)Z)} \end{aligned} \tag{33}$$

Santiago and Caballero further approximate this; we can derive their approximation by setting $Z = 1-\kappa$ and doing a first-order Taylor series expansion around $\kappa$ (since the loss in variance is presumed to be small). Then,

$$Q_\infty^2 = \frac{2}{M(1-Z)} + \frac{2(M-2)}{M^2} + \frac{2\left(M^2-4M+4\right)(1-Z)}{M^3} + \mathcal{O}(\kappa^2) \tag{34}$$

Santiago and Caballero's approximation only keeps this first term, e.g. their $Q_\infty^2 \approx 2/(M(1-Z))$, which is only accurate in the domain $M > 0.2$.

$$N_{d,\infty}^{(SC98)} \approx N \exp\left(-\frac{V_A}{(1-Z)M}\right) \tag{35}$$

(c.f. Santiago and Caballero 1998 equation 8). For small $M$, the small $\kappa$ approximation strongly deviates from the full equation; thus, throughout, we use the unapproximated version,

$$N_{d,\infty} \approx N \exp\left(-\frac{V_A}{(1-Z)(2-(2-M)Z)}\right). \tag{36}$$

For the reasons described earlier, a single asymptotic $N_{d,\infty}$ cannot fully characterize pairwise coalescent rates in all cases, and thus neutral diversity levels depend on the full $N_{d,t}$, for which we integrate equation (19) to get,

$$N_{d,t} \approx N \exp\left(-\frac{V_A}{M}\int_0^{M/2}\left(1+\frac{(1-k)(1-r)\left(1-(1-k)^t(1-r)^t\right)}{k+(1-k)r}\right)dr\right). \tag{37}$$

#### 1.7 The Variance Dynamics

The heritable variance in fitness $V_A$ associated with the tracked neutral allele does not remain constant through the generations, but decays due to selection and drift, and is maintained by the input of new mutations. For an arbitrary polygenic selective system, the dynamics are extraordinarily complicated due to the complexity of multilocus selection, and the dynamics of the trait response to selection depend on the variance, which in turn depends on higher moments of the fitness distribution (the "moment-closure problem"). We discuss the simplification of Santiago and Caballero's model here.

Each generation, the genetic variance changes due to changes in the allele frequencies at selected sites and linkage disequilibria between these sites. These are typically expressed as recursions, but certain models permit these changes to be expressed as a proportional reduction (e.g. truncation selection and stabilizing selection models; Keightley and Hill 1988 p. 36, and Walsh and Lynch 2018 p. 557). Santiago and Caballero's model follows this approach, assuming the variance decays due to selection and drift at a constant rate $Z := 1-\kappa$ through the generations, such that the variance in the next generation excluding mutation is $V_A' = (1-\kappa)V_A$ and $\Delta V_A = -\kappa V_A$.

With mutation, $\Delta V_A = V_m - \kappa V_A$. The model of Santiago and Caballero assumes that under the long-run equilibrium, $\Delta V_A = 0$, such that $\kappa V_A = V_m$, and thus the rate of loss is $\kappa = {}^{V_m}/_{V_A}$. Then the variance for the dynamics simplify to,

$$\begin{aligned} V_A' &= \left(1-\frac{V_m}{V_A}\right)V_A \\ Z &:= \frac{V_A'}{V_A} = 1-\kappa = 1-\frac{V_m}{V_A} \end{aligned} \tag{38}$$

(c.f. Santiago and Caballero 1998 equation 9).

#### 1.8 The Fitness-Effective Population Size Under Strong Background Selection

Under strong background selection, multiple segregating deleterious sites contribute to fitness variation. BGS models assume a multiplicative fitness model, such that the fitness of an individual is

$$w = (1 - s)^n \tag{39}$$

where $n$ is the number of deleterious mutations this individual carries.

Classic background selection theory assumes an infinite population size, such that mutation frequencies are at their mutation-selection equilibrium and the expected number of mutations across individuals is $\mathbb{E}(n) = {}^{\mu}/_{s} L$ where $L$ is the total number of basepairs that can be mutated (the mutational target size). Under a multiplicative fitness function in an infinite population, selection cannot generate linkage disequilibria (Turelli and Barton 1990). Thus the classic strong background selection model assumes additive genetic fitness is equal to the additive genic fitness (which excludes the contribution of covariance between selected sites),

$$\begin{aligned} V_{\text{BGS}} := \mathrm{V}(w) &= 2s^2 \sum_{l=1}^{L} p_l(1 - p_l) \\ &= 2Ls^2 \frac{\mu}{s}\left(1 - \frac{\mu}{s}\right) = Us + \mathcal{O}(\mu^2) \approx Us. \end{aligned} \tag{40}$$

since under mutation-selection balance $p_l = {}^{\mu}/_{s}$ and $U$ is defined $U := 2L\mu$, the deleterious mutation rate per diploid genome, per generation. They then set $V_A = V_{\text{BGS}}$, and note that $V_m \approx Us^2$, since mutations have frequency of ${}^{1}/_{2N}$ and each increase the variance by a factor of $s^2$. Note that because the classic BGS model assumes deterministic selection dynamics (i.e. an infinite population), the equilibrium $Z$ given by equation (38) excludes the reduction due to drift. The reduction factor under the strong BGS model is,

$$Z_{\text{BGS}} := 1 - \frac{Vm}{V_{\text{BGS}}} = 1 - \frac{Us^2}{Us} = 1 - s \tag{41}$$

(c.f. Santiago and Caballero 1998 equation 10). An alternate derivation can be found by working out the deterministic reduction in variance from the single-locus dynamics.

Now, we can use these values for $Z_{\text{BGS}}$, $V_m$, and $V$ in equation (36). Through this alternate quantitative genetics derivation, we arrive at the same equation as classic BGS theory,

$$N_{d,\infty}^{\mathrm{BGS}} = N \exp\left(-\frac{U}{2s + M - Ms}\right). \tag{42}$$

This matches equation 8 of Hudson and Kaplan (1995) except for the $Ms$ term and equation 10 of Nordborg et al. (1996) except for the $2s$ term.

##### 1.9 The Fitness-Effective Population Size Under Weak and Strong Negative Selection

With the possibility of weak selection, deleterious alleles can drift up in frequency and fix. Consequently, the fitness distribution is no longer stationary, as the set of haplotypes without any deleterious mutations can be lost due to drift. This changes the shape of the fitness distribution, and thus the variance parameter that our model relies upon (Gessler 1995; Good and Desai 2013; Haigh 1978; Higgs and Woodcock 1995; O'Fallon et al. 2010). So far, finding an explicit equation for the variance has been difficult due to the "moment-closure problem" previously mentioned. In their approximation, Santiago and Caballero (2016) consider a modified additive genic variance that is below $V_{\mathrm{BGS}}$ due to the loss of variation from each fixation of a weakly deleterious mutation. This fixation process is related to the Muller's Ratchet in assexual populations, the rate $R$ of fixation of deleterious mutations per generation per region, since each fixation directly reduces the additive genic variance of the fitness distribution. However, in general, the rate of this ratchet is unknown since it depends on the fitness variance (and thus higher moments of the fitness distribution).

In their 2016 paper, Santiago and Caballero derive use an approach based on Fisher's Fundamental Theorem of Natural Selection (equation 2, Santiago and Caballero 2016, equation 15A García-Dorado 2007). A more formal derivation follows from Higgs and Woodcock (1995) (though they assume a haploid asexual system, the end result is the same up to a factor of two). In their work, they derive equations for the change per generation for the moments and cross-moments of the fitness distribution under multiplicative selection in a haploid model. They find the haploid rate of the ratchet is $r = u - sv_n$ (equation 13.3, Higgs and Woodcock 1995), where we use lowercase to distinguish between haploid and diploid models. Here, $v_n$ is the variance in the *number* of deleterious mutations; as long as selection is not too strong, the haploid additive variance is $v_A \approx s^2 v_n$. Substituting this for $v_n$ and rearranging, the haploid fitness variance is related to the $v_A = us - rs$, which is identical to Santiago and Caballero's (2016) equation 2, noting that the waiting time between fixations (i.e. ratchet clicks) is $T = 1/r$. Intuitively, the fitness variance $V_{\mathrm{BGS}} = Us$ is reduced by the rate at which deleterious mutations fix and remove fitness variation from the population.

Now, note that under a diploid model, the fitness variance is twice that of the haploid model, so $V_A = 2v_A$. Then, the fitness variance is,

$$V_A = Us - 2Rs. \tag{43}$$

where $U = 2u$ and $R = r$ is the diploid rate of the ratchet, which depends on higher moments of the fitness distribution, but only differs from $r$ through a rescaling of the population size. We also find that Equation (5) and Supplementary Equation 43 above can also be found through Kimura's diffusion models with a flux of mutations into infinite, discrete sites (1969). Note that if selected sites are sufficiently deleterious that they cannot fix (or the population is infinite), $R = 0$ and the additive fitness variance is identical to that under strong background selection, $V_A = Us = V_{\text{BGS}}$ (equation 40). We note that while we use $V_A$ here following Santiago and Caballero (2016), this is only the *additive genic* component of additive fitness variation, as our simulations confirm in the main text.

Next, we turn to the fixation probability of a new deleterious mutation with additive effects ($1 - s$ in the heterozygote, $1 - 2s$ in the homozygote). In a diploid population with drift-effective population size $N_e$ this is,

$$p_{\text{fix}} := \frac{1 - \exp(2N_e s/N)}{1 - \exp(4N_e s)} \approx \frac{2N_e s}{N\left(\exp(4N_e s) - 1\right)} \tag{44}$$

(c.f. Durrett 2008, equation 7.21; Kimura 1957, equation 5.6). Then, Santiago and Caballero argue that the rate of the ratchet is the inverse of the averaging waiting time until a fixation, $T \approx 1/(UNp_{\text{fix}})$ (note that since $U$ is the *diploid* mutation rate, the total population mutation rate is $UN$ per generation), so

$$T \approx \frac{\exp(4N_e s) - 1}{2UsN_e}. \tag{45}$$

Since the ratchet rate depends on $N_e$, which in turn depends on the rate of the ratchet, we write the ratchet, or substitution, rate per $L$-basepair region per generation as,

$$R(N_e) := \frac{2UsN_e}{\exp(4N_e s) - 1}. \tag{46}$$

Under weak and strong background selection the variance input of mutations $V_m$ is the same, and using the $V_A$ derived above, the decay in variance is

$$Z_{\mathrm{WS}} = 1 - \frac{V_m}{V_A} = 1 - \frac{Us^2}{(U - R(N_e))s} = 1 - \frac{Us}{U - R(N_e)}. \tag{47}$$

At equilibrium, the $N_e$ that determines fixation probability is the asymptotic $N_{d,\infty}$. Consequently, Santiago and Caballero argue that the asymptotic reduction in fitness-effective population size uses the $Q^2_\infty$ and is given by the following system of non-linear equations,

$$R(N_{d,\infty}) = \frac{4UsN_{d,\infty}}{\exp(4N_{d,\infty}s) - 1} \tag{48}$$

$$N_{d,\infty} \approx N \exp\left(-\frac{V_A}{(1-Z)(2-(2-M)Z)}\right) \tag{49}$$

The solution to these equations is the asymptotic equilibrium $\widetilde{N}_d$ and equilibrium ratchet rate $\widetilde{R}$. This sets the equilibrium variance, $\widetilde{V} = Us - 2\widetilde{R}s$, or $\widetilde{V} = (U - 2\widetilde{R})s$. Note that because $p_{\text{fix}} < 1/2N$ for deleterious variation, $2R < U$, or the diploid mutation rate must be greater than or equal to twice the per-basepair per-generation substitution rate for equilibrium variance $\widetilde{V} > 0$ for $s > 0$.

### 2 Statistical Methods

The previous section outlined the basis of quantitative genetic linked selection models that predict the fitness-effective population size under polygenic selection and linkage. Our work here only considers the contribution of deleterious mutations to fitness variation. In this section, we outline more details about the statistical models used to construct the genome-wide B and B' maps that are used in model fitting.

#### 2.1 Models of Linked Selection

Both the classic BGS and the SC16 model we extend have the same parametric form, since both (1) model the impact of linked selection as a rescaling of effective population size, and (2) both assume multiplicative fitness across segments. Thus, the total reduction experienced by a neutral site at position $x$ due to segments across the genome is the product of the individual reductions,

$$\begin{aligned} B(x, \Psi) &= \prod_{g \in G} \exp\left(\int_0^1 b(\mu(s, k(g)), s, L_g, r_g, d(x, g)) ds\right) \\ &= \exp\left(\sum_{g \in G} \int_0^1 b(\mu(s, k(g)), s, L_g, r_g, d(x, g)) ds\right) \end{aligned} \tag{50}$$

where, $b(\mu(s, k(g)), s, L_g, r_g, d(x, g))$ is the log reduction due to segment $g$, and there are $S$ total segments. Here, $\mu(s, k(g))$ is the per-basepair per-genome rate that mutations with selection coefficient $s$ enter segments with feature class $k(g)$ (e.g. it incorporates the distribution of fitness effects and mutation rate), $L_g$ and $r_g$ are the length in basepairs and recombination rate per basepair of segment $g$, and the recombination distance between focal site $x$ and segment $g$ is $d(x, g)$. The functional form of $b$ varies depending on whether the classic BGS model is used (Supplementary Section 2.3), or our form of Santiago and Caballero's (2016) equations (see Supplementary Section 2.4).

Under both negative selection models implemented in our software, the diversity in window $v$ is $\bar{\pi}(v, \Psi) = \bar{B}(v, \Psi)\pi_0$. Here, $\bar{B}(v|\Psi)$ is the predicted reduction in diversity due to BGS in window $v$, given background selection parameters $\Psi$. In practice, we pre-calculate $B(x|\Psi)$ at fixed sites $x$ across the genome, and take the average of the fixed sites $B$ values within window $v$ for the average $\bar{B}(v|\Psi)$.

#### 2.2 Discretization of Parameters and Annotation Feature Classes

In practice to fit negative selection models we must discretize both $\mu(s, k(g))$ and $s$ in Supplementary Equation (50). We define $\mu_{i,k(g)}$ to be the rate of mutations entering the population per basepair per generation with selection coefficient $s_i$ for the feature class $k(g)$ of segment $g$. This $\mu_{i,k(g)}$ term reflects the product of the mutation rate and the distribution of deleterious fitness effects for selection coefficient $s_i$ and feature class $k(g)$. Then, the reduction at position $x$ is,

$$B(x, \Psi) = \exp\left(\sum_{g}^{S}\sum_{i}^{m_s} b(\mu_{i,\,k(g)}, s_i, L_g, r_g, d(x, g))\right). \tag{51}$$

Because it is computationally-intensive to numerically solve the system of two non-linear equations (Equation 6 in the main paper) to calculate $b$ at each segment, these are pre-computed for an $n_s$-element grid of selection coefficients and an $n_\mu$-element grid of mutation rates–DFE products. Likewise, when we calculate $b$ under classic BGS models, we pre-calculate segment components across these same grids, and combine them for each $x$.

The number of annotation features $K$ is variable and set by the annotation data specified. There are two ways to parameterize the mutation rates and DFE for all annotation classes in background selection models. First, for the *free-mutation* parameterization, each annotation feature class has a free mutation rate parameter for each of these selection coefficients (as in Supplementary Equation 50). We experimented with this approach, but ultimately found its results less intuitive and it has more degrees of freedom.

Instead, we opted for a *simplex* parameterization, which has a single shared mutation rate across all features, so the mutation parameter $\mu_{i,\,k(g)}$ above is given by the product of the mutation rate and the DFE weight matrix. For example, for a $\log_{10}$ selection grid and three features, CDS, UTRs, and PhastCons (PC), we'd have a DFE matrix like,

$$\mathbf{M} = \mu\mathbf{W} = \begin{bmatrix} w_{10^{-6},\text{CDS}} & w_{10^{-6},\text{UTR}} & w_{10^{-6},\text{PC}} \\ w_{10^{-5},\text{CDS}} & w_{10^{-5},\text{UTR}} & w_{10^{-5},\text{PC}} \\ \vdots & \vdots & \vdots \\ w_{10^{-1},\text{CDS}} & w_{10^{-1},\text{UTR}} & w_{10^{-1},\text{PC}} \end{bmatrix} \tag{52}$$

which has $2 + K \times (n_s - 1)$ free parameters, since each column must sum to one. This imposes the constraint that $\sum_i w_{i,k} = 1$, requiring constrained MLE optimization. We implement this using softmax, which describe further in Supplementary Section 3.12.

#### 2.3 Reductions Under the Classic BGS Model

The classic BGS model, which assumes deleterious mutations cannot fix, expresses the reduction for a focal site in the middle of a segment (Hudson and Kaplan 1995; Hudson and Kaplan 1994; Nordborg et al. 1996); here we consider a focal site directly to an arbitrary side of a segment under a deleterious mutations, and extend the model to handle a focal segment arbitrarily far from the segment. To simplify notation, we will only consider a single feature class in this and the next section. The log reduction under the classic mutation-selection-balance BGS model is,

$$b(\mu, s, L, r, 0) = -\int_0^L \frac{\mu(l)}{s(1 + (1-s)r(l)/s)^2} dl \tag{53}$$

(c.f. McVicker et al. 2009 p. 11, Hudson and Kaplan 1995 equation 5,), where $\mu(l)$ is the *haploid* per-basepair mutation rate at $l$ and $r(l)$ is the recombination fraction between the focal site (at position 0) and basepair $l$. Assuming constant per-basepair recombination and mutation rates, $r(l) = r_{\text{BP}} l$ and $\mu(l) = \mu$,

$$b(\mu, s, L, r_{\text{BP}}, 0) = -\frac{\mu L}{s + (1-s) r_{\text{BP}} L} \tag{54}$$

(c.f. Hudson and Kaplan 1995 equation 8; their model considers a neutral site in the middle of the segment and drops small terms). Often per-region rates are defined $U = 2\mu L$ and $M = r_{\text{BP}} L$ are the segment-wide mutation diploid mutation rates (mutations per diploid segment per generation) and recombination length (Morgans).

For computational efficiency, we adjust this model by setting $r(l) = d + r_{\text{BP}} l$ in Supplementary Equation (53) and integrating, where $d$ is the recombination fraction between the focal site $x$ and the start of segment $g$. This allows us to pre-compute the local effects of the segment, and substitute in $b$ as the focal site changes. This integral substitution has a closed form,

$$b(\mu, s, L, r_{\mathrm{BP}}, d) = -\frac{\mu L}{(d(1-s)+s)(s+(d+r_{\mathrm{BP}}L)(1-s))}. \tag{55}$$

Then, terms can be collected in powers of $d$ and pre-computed for all segments for the grid of $\mu$ and $s$. Our implementation of classic BGS in `bgspy` pre-computes these segment components, and then calculates the final $b$ value for an $x$ and $g$ by calculating the distance $d(x, g) = |M(x) - M(g)|$, where $M(x)$ is the cumulative recombination map length at position $x$ in Morgans. Since $r(l)$ could vary by position, segments with differing recombination rates are split in two of the same annotation class by their recombination rate. This has no effect on the calculation due to the multiplicative fitness, but ensures more accurate estimates of the reduction factor $B$.

#### 2.4 Reduction Under the Modified Santiago and Caballero Model

Under Santiago and Caballero's model, the correct effective population size to gauge reduction in heterozygosity or coalescent times is given by their equation 4 (Santiago and Caballero 2016). This equation considers how the inflation factor $Q_t^2$ that sets the fitness-effective population size varies over $t$. This is because statistics such as heterozygosity depend on the cumulative autocorrelation factor up to some time $t$, $Q_t^2$ (Santiago and Caballero 1998, p. 2111). This reflects the fact that effective population size experienced by a mutation newly associated with a fitness background is larger than the asymptotic effective population size, since not much autocorrelation has accumulated in the stochastic perturbations caused by drift yet. From a backwards-time perspective, this reflects the fact that pairwise diversity (which is approximately heterozygosity when $2N\mu \ll 1$) depends on the pairwise coalescence rate per generation, which is not constant under linked selection.

We experimented with calculating this full sum and integral approximations to it in our methods, but it seemed to make little difference in our application. Additionally, it was too costly to calculate computationally for genome-wide calculations of the B' map. Instead, we solve for the asymptotic $\widetilde{N}_{e,\infty}$ for each segment, and use that to calculate the log reduction factor under the SC16 model, $b'$.

Note that the asymptotic model above determines the reduction in effective population size experienced by a focal site in the middle of a $L$-basepair segment, but B maps require the reduction $b'$ experienced by a focal site $x$ at an arbitrary recombination distance apart. For each segment, we pre-compute the solution to the system of two equations in Supplementary Equation (48) for the mutation and selection coefficient grids, giving us $\widetilde{V}_a$ and $\widetilde{Q}_\infty^2$ for each set of parameters. Then,

$$b'(\mu, s, L, r, d) = -\frac{\widetilde{V}_a(\mu, s, r)}{2} Q^2(\mu, s, r) \tag{56}$$

$$Q^2(\mu, s, r) = \left( \frac{1}{(1 - (1 - d)\widetilde{Z}} \right)^2 \tag{57}$$

where $\widetilde{Z} = 1 - Us/(U - \widetilde{R})$ and $\widetilde{V}_a = Us - 2\widetilde{R}s$. As before, $d$ is the recombination fraction between the segment and the focal neutral site.

##### 2.5 Data Summary Matrices

Our underlying data for all likelihood and pairwise diversity estimates is the allele count matrix $\mathbf{C}$ with dimensions $L \times 2$, where $L$ is the chromosome length. This is transformed to a pairwise summary matrix with identical dimensions, $\mathbf{Y}$. The first column of $\mathbf{Y}$ is the number of pairwise comparisons between chromosomes that are identical, and the second column is the number that are different. Both of these columns are combinatoric summaries of the raw allele counts matrix needed for the binomial likelihood and pairwise diversity estimates. Let $[c_1, c_2]$ be a row of $\mathbf{C}$ for basepair $l$ (the $l$ index is omitted for clarity), and $n = c_1 + c_2$. Then, the (1) total number of pairwise combinations of chromosomes $n_T$, (2) the number of pairwise with identical alleles $n_S$, and (3) the number of pairwise combinations with differing alleles $n_D$ are respectively,

$$\begin{aligned} n_T &= \frac{n(n-1)}{2} \\ n_S &= \binom{c_1}{2} + \binom{c_2}{2} \\ n_D &= n_T - n_S \end{aligned} \tag{58}$$

which would be stored in row $\mathbf{Y}_l = [n_S, n_D]$. Note that the per-site $\mathbf{Y}$ handles non-polymorphic sites and missing data. Non-polymorphic sites have $n_S = \binom{n}{2}$ and $n_D = 0$, and missing data has $n_S = n_D = 0$.

##### 2.6 Pairwise Diversity Estimates

The pairwise diversity at site $l$ across the $n$ sampled chromosomes can be calculated from row $l$ of the $\mathbf{Y}$ matrix as follows,

$$\pi_l = \frac{n_D}{n_T} \tag{59}$$

which is identical to the more common expression of this estimator,

$$\pi_l = \frac{2}{n(n-1)} \sum_{i<j}^{n} k_{i,j} \tag{60}$$

where $k_{i,j}$ is 1 if the alleles at this site differ at site $l$, and 0 otherwise.

There are three ways to aggregate $\pi_l$ across all sites. The first is,

$$\pi^{(1)} = \frac{1}{L} \sum_{i=1}^{L} \frac{n_{D,i}}{n_{T,i}} \tag{61}$$

which if the number of samples across loci is constant, simplifies to an unweighted average across sites. Second, one can take a weighted average, with weights determined by the total number of samples present at a site,

$$\pi^{(2)} = \frac{1}{\sum_{i=1}^{L} n_i} \sum_{i=1}^{L} n_i \frac{n_{D,i}}{n_{T,i}} \tag{62}$$

Third, one can weight by the number of pairwise comparisons at a site, $n_{T,i}$, rather than total number of samples, $n_i$, which leads to a ratio of sums,

$$\pi^{(3)} = \frac{\sum_{i=1}^{L} n_{D,i}}{\sum_{i=1}^{L} n_{T,i}}. \tag{63}$$

We predominantly use the estimator $\pi^{(3)}$, as it corresponds to how we summarize the matrix $\mathbf{Y}$ across windows for our likelihood. All methods have mean squared errors and biases very close to one another.

Note, however, that estimates of pairwise diversity often condition on the accessible bases, and thus treat this as fixed. However, the number of accessible bases varies across the chromosome; this can lead to a source of apparent bias during block-bootstrap estimates of uncertainty. In this case, pairwise diversity is a ratio estimator, and is thus biased, since by Jensen's inequality $\mathbb{E}(y/x) \geq \mathbb{E}(y)/\mathbb{E}(x)$ for random variables $x$ and $y$.

### 3 Analysis Details

#### 3.1 Human Genomic Data

In order to try to ensure accurate estimation of pairwise diversity, which is a ratio estimator that is sensitivity to its denominator, we use the complete per-basepair genotype calls (gVCF) produced by Illumina's DRAGEN pipeline for the high-coverage 1000 Genome samples (Byrska-Bishop et al. 2022; Illumina, Inc. 2020). The original samples were sequenced to 30x by the New York Genome Center (Byrska-Bishop et al. 2022). This allows for filtering to be applied to the entire genome at once, rather than just variants, so the denominator does not need to be estimated separately.

#### 3.2 Filtering gVCFs

The gVCFs were filtered using a custom Python tool, `gvcf2counts.py` (available in the repository associated with the manuscript at `tools/gvcf2counts.py`), which reads the gVCFs, filters them according to the criteria below, and outputs a Numpy `.npy` file of reference and alternative allele counts for each chromosome (hereafter, "allele counts matrices").

Genotypes are included in the allele count if and only if:

1. The variant call is set to `PASS` in the VCF.
2. The `QUAL > 50`.
3. The `GQ > 30` (or `RGQ > 30` for invariant sites).

Because the data underlying the counts files are per-basepair resolution gVCFs, each chromosome's allele counts matrix is of size $l \times 2$, where $l$ is the total chromosome length. Basepairs that fail these filtering requirements lead to a row of zero counts, e.g. no observed reference and alternative allele counts, and thus do not effect the data that goes into the binomial likelihood or $\pi$ estimates used in figures.

#### 3.3 Non-accessible and Masked Regions

The allele counts matrices include many basepairs that may have allele counts that pass the genotype call filters, but are still need to be filtered out because the region of the genome may produce unreliable estimates. The following filters are applied based on masking regions:

1. **Non-accessible regions**: masks out centromeres (`acen` entries in `cytoBand.txt`), with 5 Mbp padding on either side. The file of passing masks is `data/annotation/no_centro.bed`.
2. **Reference masking**: soft and hard-masked regions in the human GRCh38 reference genome are also masked. Soft-masked regions were determined by Ensembl (Cunningham et al. 2022),

which uses Repeat Masker (Smit et al. 2015) and Dust (Morgulis et al. 2006). We also only used sites that pass the 1000 Genomes strict filter (1000 Genomes Project Consortium et al. 2015).

#### 3.4 Putatively Neutral Tracks

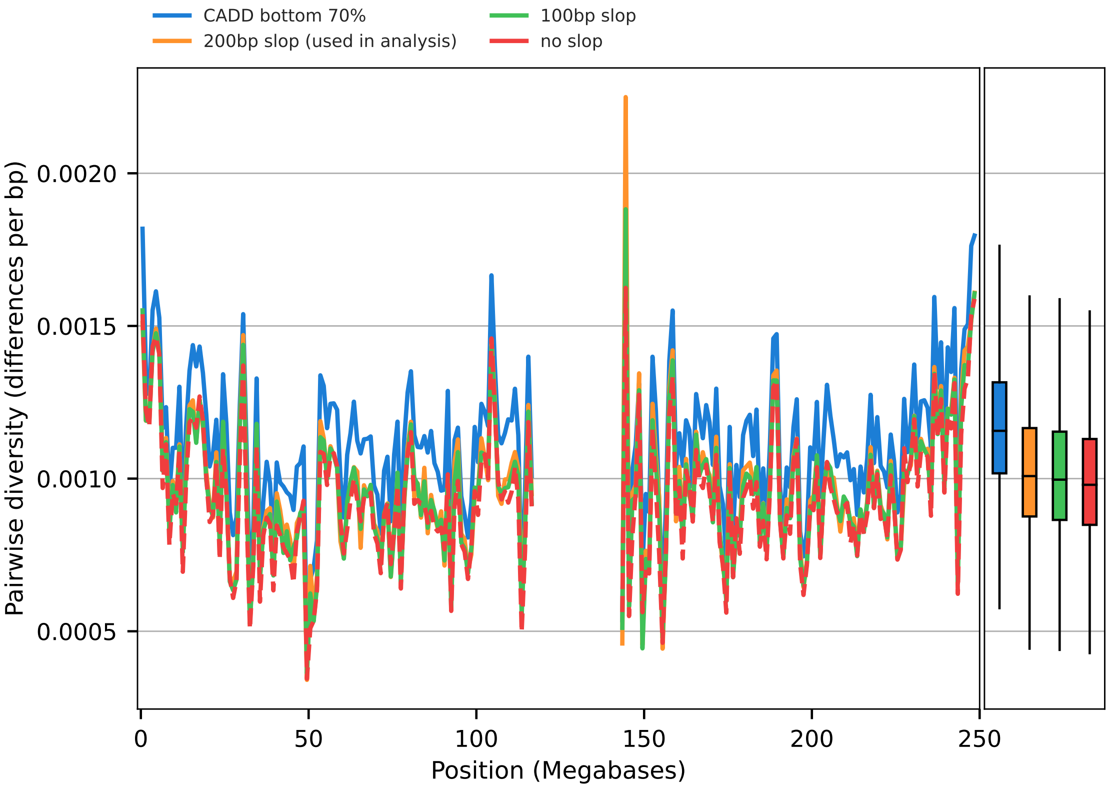

Figure 1: Estimates of chromosome 1 YRI diversity in non-overlapping megabase windows, under different neutral tracks (colors) and the filtering criteria described in Supplementary Section 3.3. To the right of the chromosome subfigure are the genome-wide boxplots of megabase-scale diversity.

Linked selection theory typically assumes that focal sites are not under purifying selection themselves. Consequently, linked selection statistical methods often only consider the fraction of the genome not *a priori* under selective constraint (Murphy et al. 2022, e.g. Appendix Section 3.1). Our methods follow suit, and first create a track consisting of the union (i.e. `bedtools merge`) of: exons and coding sequences from Ensembl Version 107 `Homo_sapiens.GRCh38.107.chr.gff3.gz`]Cunningham2022-vk and PhastCons regions `phastConsElements100way.bed.gz`]Siepel2005-wh. Then, we remove 200 basepairs of buffer around each of these regions (i.e. `bedtools slop`). We show in Supplementary Figure 1 that average diversity (we only consider YRI samples for this analysis) is fairly insensitive to the buffer width, but increasing slop widths around putatively conserved segments does lead to higher diversity estimates. This is expected, since increasing the slop width considers a smaller subset of sites, further away from putatively conserved regions. For our analyses we choose the 200bp slop version, which covers 63.66% of the autosomal reference sequence and consists of 1.26 million contiguous genomic regions.

Additionally, we compared our neutral track (and the slop width alternatives) against the bottom 70% of CADD site scores (this works out to be 64.70% of the autosomal reference sequence). The correlation between these two tracks is extraordinarily high (Pearson $r = 0.98$, p-value = 0), but the average pairwise diversity is 13% higher (two-tailed t-test p-value $1.29 \times 10^{-71}$) in the CADD bottom 70% neutral track. However, we did not use the CADD bottom 70% neutral track in our analysis of human population genomic data. Our concern was that because CADD models are trained by differentiating high-frequency derived SNPs in humans from simulated variants at every basepair, there could be circularity in basing our neutral regions on predictions based on modeling frequency data. Moreover, this neutral track would be close to a complement of the CADD 6% track, which we use as one of our annotation models, and might lead to issues. Additionally, the CADD bottom 70% neutral track is much less sparse than the track we used in our analysis, with a total of $\approx 20$ times more contiguous regions than the neutral track used in analysis (25.38 million total). Many of these ranges are at very small spatial scales (average: $\approx 7$ basepairs).

Additionally, we ran one test model with the CADD bottom 70% as the neutral track as a test. Using the CADD bottom 70% neutral track with the CADD 6% annotation model, the in-sample was $R^2 = 63.99\%$; by comparison, the in-sample $R^2 = 68.03\%$ using the constructed neutral track with 200bp slop. Overall, future work calibrating the neutral track is needed.

These neutral tracks are all produced by the Snakemake file `data/fit_annotation/Snakefile` in this study's GitHub repository (`http://github.com/vsbuffalo/bprime`). We summarize the percentage of basepairs after masking per chromosome in Supplementary Table 1.

#### 3.5 Window-based Summaries and filtering

Based on exploratory data analyses, we noticed some rare windows that were outliers. Given that maximum likelihood estimation can be easily biased by outliers, we took a conservative approach and removed the top 0.5% of high-diversity windows. The results of this can be seen in Supplementary Figure 2. We also remove 1cM from each end of the recombination map. These window-based filters are in addition to the accessible and neutral sequence masked described in Supplementary Sections 3.2 and 3.3.

Table 1: Accessible and putatively neutral sequence mask statistics, for the criteria used in the main model fits.

| Chromosome | Accessible (%) | Neutral (%) | Intersection (%) | Total Basepairs ($\times 10^6$) |
|---|---|---|---|---|
| 1 | 39.7 | 62.0 | 15.7 | 39.09 |
| 2 | 45.3 | 60.1 | 18.9 | 45.77 |
| 3 | 43.2 | 59.6 | 16.9 | 33.51 |
| 4 | 42.7 | 65.9 | 20.2 | 38.42 |
| 5 | 42.4 | 62.2 | 18.1 | 32.86 |
| 6 | 43.8 | 61.4 | 18.4 | 31.43 |
| 7 | 41.9 | 64.0 | 18.6 | 29.64 |
| 8 | 42.3 | 66.1 | 20.2 | 29.32 |
| 9 | 36.2 | 65.4 | 14.6 | 20.21 |
| 10 | 42.9 | 62.2 | 18.8 | 25.15 |
| 11 | 41.3 | 61.5 | 17.0 | 22.96 |
| 12 | 40.6 | 60.4 | 15.9 | 21.19 |
| 13 | 38.8 | 70.7 | 18.8 | 21.50 |
| 14 | 36.5 | 64.6 | 13.8 | 14.77 |
| 15 | 34.4 | 66.3 | 13.5 | 13.77 |
| 16 | 36.0 | 66.0 | 15.5 | 14.00 |
| 17 | 37.2 | 58.8 | 14.2 | 11.82 |
| 18 | 40.3 | 66.1 | 19.6 | 15.75 |
| 19 | 28.1 | 67.5 | 11.5 | 6.74 |
| 20 | 36.1 | 62.5 | 15.4 | 9.92 |
| 21 | 29.8 | 76.9 | 17.3 | 8.08 |
| 22 | 25.5 | 75.5 | 12.6 | 6.40 |
| weighted average | 40.1 | 63.6 | 17.1 | - |

Table 2: $R^2$ and mutation rate estimates for all primary models fit on the default DFE grid (up to selection $s = 10^{-1}$). $R^2_{\text{IS}}$ is the in-sample $R^2$, and $R^2_{\text{LOCO}}$ is the leave-one-chromosome-out $R^2$.

| **Model** | **Track type** | **Pop.** | **B' $R^2_{\text{LOCO}}$** | **B' $R^2_{\text{IS}}$** | **B $R^2_{\text{IS}}$** | **B' $\hat{\mu} \times 10^{-8}$** | **B $\hat{\mu} \times 10^{-8}$** |
|---|---|---|---|---|---|---|---|
| PhastCons Priority | sparse | YRI | 67.29 | 67.36 | 62.82 | 2.0039 | 0.2070 |
| PhastCons Priority | full | YRI | 67.27 | 67.36 | 58.26 | 2.0037 | 0.1037 |
| CADD 6% | sparse | YRI | 66.66 | 66.81 | 66.79 | 1.6565 | 1.6328 |
| CADD 6% | full | YRI | 66.50 | 66.68 | 58.76 | 1.5556 | 0.1074 |
| CADD 8% | sparse | YRI | 65.30 | 65.58 | 65.56 | 1.2229 | 1.1997 |
| CADD 8% | full | YRI | 65.13 | 65.41 | 59.16 | 1.1260 | 0.1067 |
| Feature Priority | sparse | YRI | 63.61 | 64.28 | 60.43 | 3.3528 | 0.2259 |
| Feature Priority | full | YRI | 63.06 | 64.28 | 56.08 | 3.3490 | 0.1000 |
| PhastCons Priority | sparse | CEU | 61.04 | 62.25 | 58.37 | 2.3133 | 0.2160 |
| PhastCons Priority | full | CEU | 61.04 | 62.25 | 53.95 | 1.9117 | 0.1044 |
| CADD 6% | sparse | CEU | 60.37 | 61.80 | 61.79 | 1.6797 | 1.6670 |
| CADD 6% | full | CEU | 60.24 | 61.77 | 53.99 | 1.6347 | 0.1096 |
| CADD 8% | sparse | CEU | 59.38 | 60.92 | 60.91 | 1.2547 | 1.2418 |
| CADD 8% | full | CEU | 59.24 | 60.88 | 54.31 | 1.2078 | 0.1090 |
| PhastCons Priority | sparse | CHB | 58.84 | 60.45 | 56.55 | 1.9202 | 0.2240 |
| PhastCons Priority | full | CHB | 58.76 | 60.45 | 52.93 | 1.8689 | 0.1101 |
| CADD 6% | sparse | CHB | 58.67 | 60.49 | 60.48 | 1.7454 | 1.7334 |
| CADD 6% | full | CHB | 58.48 | 60.40 | 52.76 | 1.5981 | 0.1166 |
| CADD 8% | sparse | CHB | 57.91 | 59.78 | 59.76 | 1.3093 | 1.3022 |
| CADD 8% | full | CHB | 57.74 | 59.69 | 53.14 | 1.2038 | 0.1161 |
| Feature Priority | sparse | CEU | 57.24 | 59.11 | 56.35 | 3.0442 | 0.2313 |
| Feature Priority | full | CEU | 57.14 | 59.11 | 52.21 | 3.0441 | 0.1004 |
| Feature Priority | sparse | CHB | 56.00 | 57.88 | 54.76 | 3.1722 | 0.2362 |
| Feature Priority | full | CHB | 55.63 | 57.88 | 51.39 | 3.1722 | 0.1056 |

Table 3: $R^2$ and mutation rate estimates for all primary models fit with the strong selection grid (up to $s = 10^{-1}$). Note that the repeated values in the last rows are correct and due to rounding; see the main analysis notebook in the GitHub repository for more information. $R^2_{\text{IS}}$ is the in-sample $R^2$, and $R^2_{\text{LOCO}}$ is the leave-one-chromosome-out $R^2$.

| **Model** | **Track type** | **Pop.** | **B'** $R^2_{\text{LOCO}}$ | **B'** $R^2_{\text{IS}}$ | **B** $R^2_{\text{IS}}$ | **B'** $\hat{\mu} \times 10^{-8}$ | **B** $\hat{\mu} \times 10^{-8}$ |
|---|---|---|---|---|---|---|---|
| PhastCons Priority | sparse | YRI | 68.45 | 68.16 | 65.23 | 1.8636 | 0.2975 |
| PhastCons Priority | full | YRI | 68.19 | 68.12 | 63.67 | 1.7082 | 0.1666 |
| CADD 6% | full | YRI | 68.08 | 68.08 | 62.96 | 1.4612 | 0.1721 |
| CADD 6% | sparse | YRI | 68.04 | 68.03 | 68.01 | 2.0912 | 2.0716 |
| CADD 8% | full | YRI | 66.98 | 67.11 | 63.61 | 1.0126 | 0.1713 |
| CADD 8% | sparse | YRI | 66.67 | 67.01 | 66.98 | 1.5688 | 1.5473 |
| Feature Priority | full | YRI | 64.90 | 65.51 | 61.23 | 4.1951 | 0.1620 |
| Feature Priority | sparse | YRI | 64.90 | 65.51 | 63.16 | 4.1795 | 0.3070 |
| PhastCons Priority | sparse | CEU | 61.98 | 63.02 | 60.02 | 2.0194 | 0.3036 |
| PhastCons Priority | full | CEU | 61.78 | 63.01 | 58.57 | 2.3499 | 0.1679 |
| CADD 6% | full | CEU | 61.74 | 62.94 | 57.42 | 1.5365 | 0.1739 |
| CADD 6% | sparse | CEU | 61.66 | 62.86 | 62.84 | 2.1215 | 2.1084 |
| CADD 8% | full | CEU | 60.80 | 62.12 | 57.95 | 1.1572 | 0.1742 |
| CADD 8% | sparse | CEU | 60.59 | 62.06 | 62.04 | 1.6030 | 1.5876 |
| CADD 6% | full | CHB | 59.62 | 61.31 | 55.31 | 1.5083 | 0.1727 |
| CADD 6% | sparse | CHB | 59.54 | 61.21 | 61.20 | 2.1242 | 2.1170 |
| PhastCons Priority | sparse | CHB | 59.44 | 61.08 | 57.88 | 5.8874 | 0.3060 |
| PhastCons Priority | full | CHB | 59.29 | 61.08 | 56.71 | 2.5573 | 0.1694 |
| CADD 8% | full | CHB | 58.88 | 60.63 | 55.86 | 1.1128 | 0.1733 |
| CADD 8% | sparse | CHB | 58.65 | 60.57 | 60.55 | 1.6098 | 1.6001 |
| Feature Priority | sparse | CEU | 58.58 | 60.30 | 58.41 | 4.1041 | 0.3110 |
| Feature Priority | full | CEU | 58.58 | 60.30 | 56.75 | 4.1001 | 0.1648 |
| Feature Priority | sparse | CHB | 56.48 | 58.73 | 56.60 | 3.7195 | 0.3136 |
| Feature Priority | full | CHB | 56.48 | 58.73 | 54.95 | 3.7552 | 0.1654 |

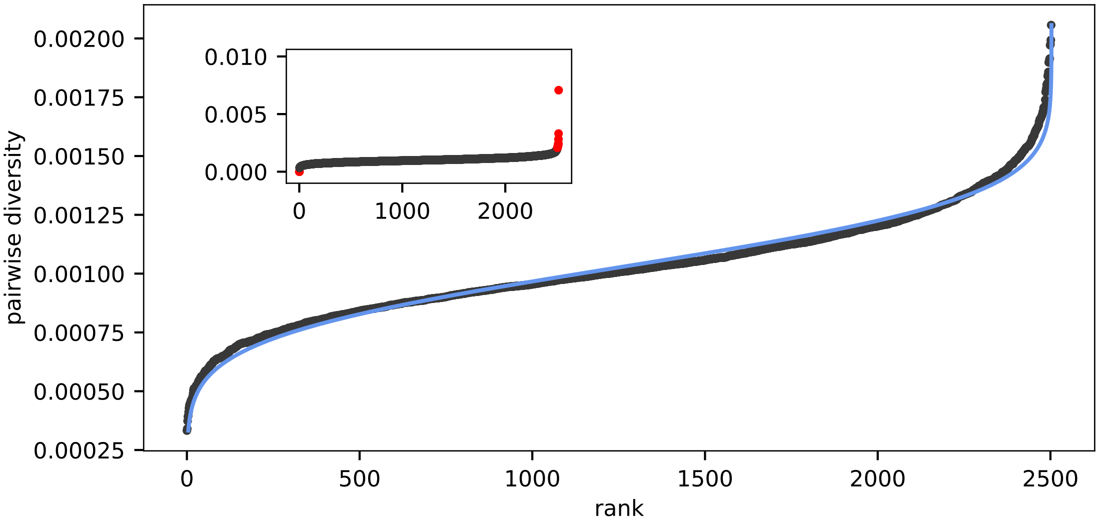

Figure 2: The distribution of diversity across the genome at the megabase scale, with outliers trimmed. The blue line is the best-fitting normal CDF. We note that this implies the distribution of megabase-scale diversity is slightly fat-tailed compared to a normal distribution. The inset figure is the untrimmed data, with trimmed points shown in red.

##### 3.6 Model Comparisons

##### 3.7 Table of $R^2$ Values

##### 3.8 Table of $R^2$ Values (Strong Selection Grid)

##### 3.9 Derivation of Approximate Coalescence $R^2_{\textbf{coal}}$

Our approach models the observed megabase-scale diversity $y_i$ in window $i$ as the predicted level under our negative selection model, $\pi_0 B_i$, plus some random residual error $\varepsilon_i$,

$$y_i = \pi_0 B_i + \varepsilon_i \tag{64}$$

where each $B_i = f(X_i|\Psi)$ for some data $X_i$ (i.e. the recombinaton map and annotation model) and parameters $\Psi$. To compare our models and assess model goodness-of-fit, we use the out-sample $R^2_{\text{LOCO}}$ which is calculated by fitting the genome-wide model leaving one chromosome out, and calculating the residual variance between predictions and observed values for this out-sample chromosome.

In general, $R^2$ is calculated for a model fit across windows as,

$$R^2 = 1 - \frac{V_{\text{res}}}{V_{\text{tot}}} \tag{65}$$

where $V_{\text{res}}$ is the mean squared error across windows and $V_{\text{tot}}$ is the total variance in the observed values. Suppose we have estimates $\widehat{\pi}_0 \widehat{B}_i$ for $\pi_0 B_i$ from our model. Then,

$$V_{\text{res}} = \frac{1}{n} \sum_{i=1}^{n} (y_i - \widehat{\pi}_0 \widehat{B}_i)^2 \tag{66}$$

$$= \underbrace{\frac{1}{n} \sum_{i=1}^{n} \varepsilon_i^2}_{\text{irreducible error}} + \underbrace{\left( \frac{1}{n} \sum_{i=1}^{n} (\pi_0 B_i - \widehat{\pi}_0 \widehat{B}_i) \right)^2}_{\text{bias squared}}. \tag{67}$$

The total variance among the observed diversity values is,

$$V_{\text{tot}} = V_{\text{res}} + V_{\text{model}} + 2\widehat{\pi}_0 \operatorname{cov}_i(\varepsilon_i, \widehat{B}_i) \tag{68}$$

where

$$V_{\text{model}} = \frac{\widehat{\pi}_0^2}{n} \sum_{i=1}^{n} \left( \widehat{B}_i - \frac{1}{n} \sum_{i=1}^{n} \widehat{B}_i \right)^2. \tag{69}$$

Thus, our $R^2$ depends on (1) the bias of our predictors, (2) the covariance between residuals and predictors, and (3) the irreducible error due to variance in diversity within windows. However, it is of interest to approximate the irreducible error under theoretic models that assume all irreducible error is due entirely to the variance of coalescence times in a window, assuming that all of the effects of negative selection on the variance can be thought of as a local reduction in the effective population size to $B_i N_e$ and demographic effects amount to a simple rescaling of $N_e$. This also assumes mutation rates are constant across the genome, and do not contribute to the irreducible variance. We call this theoretic goodness-of-fit under drift $R^2_{\text{drift}}$, and it provides a rough approximation of the irreducible error in our model due to "coalescence noise" around the modeled reductions due to negative selection.

Then,

$$\mathbb{E}(\varepsilon_i^2) = \mathrm{V}(\pi_i) \tag{70}$$

since the mean residual is zero. The variance in pairwise diversity $\mathrm{V}(\pi_i)$ is equivalent to the variance in the number of segregating sites for a sample of two, $\text{V}(\pi) = \text{V}(S_2)$. The variance in coalescence times in a window of width $w$ basepairs with population-scaled recombination rate $\rho = 4B_i N_e r w$ is,

$$\text{V}(S_2) = \theta + \theta^2 \frac{2}{\rho^2} \int_0^\rho (\rho - x) f_2(x) dx \tag{71}$$

(Wakeley 2009, eq. 7.20) where,

$$f_2(\rho) = \frac{\rho + 18}{\rho^2 + 13\rho + 18}. \tag{72}$$

Since this is a rough approximation, we set $\theta = w\pi_0 B_i$ and fix $\rho = w\gamma\pi_0 B_i$ where $\gamma$ is the ratio of recombination to mutation rates, $r/\mu$. In humans, $\gamma \approx 1$, but we explore different ratios to assess sensitivity of this calculation (Supplementary Figure 3B).

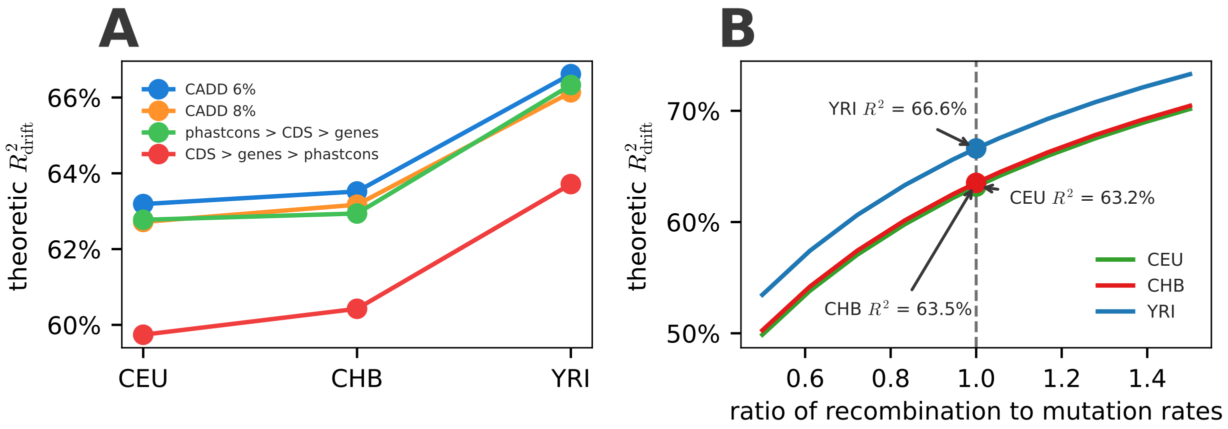

Figure 3: (A) the theoretic $R^2_{\text{drift}}$ using different plugin estimators for $B_i$ under different models. This shows that among the best-fitting models, the predicted $R^2_{\text{drift}}$ varies little; most variation is among populations due to differing $\pi_0$. (B) The $R^2_{\text{drift}}$ across different populations, for varying ratio of recombination to mutation rates (x-axis).

This back-of-the-envelope calculation finds that the upper bound of $R^2$ would be around 67% for Yoruba, and 64% for European and Han Chinese models (Figure 3 This rough approximation assumes constant demography, which for bottlenecked out-of-Africa populations assumes that the variance in coalescence times is determined entirely by reducing the effective population size. However, the variance in coalescence times is *higher* in bottlenecked populations (CEU and CHB) compared to Yoruba, which we confirm with simple simulations using the `OutOfAfrica_3G09` model from `stdpopsim` (Adrion et al. 2020; Gutenkunst et al. 2009). This higher than expected variance would inflate $SS_{\text{res}}$, which would decrease $R^2_{\text{drift}}$, bringing it closer to the observed $R^2_{\text{LOCO}}$. A more accurate approximation of the $R^2_{\text{drift}}$ could be found with simulations with more realistic demography and varying coalescence rates along the genome.

##### 3.10 Predicted Rate of Fitness Change

Each fixation of a mutant with fitness effect $-s$ reduces the population fitness to $1-2s$ since all individuals are homozygotes for the deleterious mutation and we assume additive effects at a site. For each segment $l$, the maximum likelihood parameter estimates of our model imply a prediction of the deleterious substitution rate per segment $\widehat{R}_{l,i}$ for each selection coefficient $s_i$ in the $m_s$-element DFE grid (according to which feature class it belongs to).

Since we assume multiplicative fitness effects, the log population fitness (considering only fixed sites) for segment $l$ in the next generation is,

$$\log(L_l) = \sum_{i=1}^{m_s} \log(1-2s_i)\widehat{R}_{l,i}. \tag{73}$$

Since we assume multiplicative fitness, the total genome-wide log population fitness in the next generation is

$$\log(L) = \sum_{l} \log L_l. \tag{74}$$

This amounts to a predicted population fitness loss per generation of

$$\begin{aligned} \widehat{\Delta}_W &= 1 - L \\ &= 1 - \exp\left(\sum_l \sum_{i=1}^{m_s} \log(1-2s_i)\widehat{R}_{l,i}\right). \end{aligned} \tag{75}$$

We calculate this value for all segments within a feature class, and then sum over all features to arrive at our genome-wide estimate. We use a parametric bootstrap to carry forward the estimation uncertainty to the predicted fitness loss per generation. Briefly, we do this by sampling new estimates of the DFE per selection coefficient and feature from a normal distribution centered on the ML point estimate and with the jackknife standard deviation.

##### 3.11 Likelihood

Our model is essentially a generalized nonlinear model with a binomial link function. This is the form used by Elyashiv et al. (2016) and Murphy et al. (2022). These models fit the observed number of pairwise nucleotide differences in genomic windows to the expected pairwise diversity under some evolutionary model. We only consider models of purifying selection here, so the mean function for position $x$ is the product of the proportion by which selection reduces diversity at position $x$, $B(x)$, and genome-wide neutral diversity $\pi_0$,

$$\pi(x) = B(x, \Phi)\pi_0 \tag{76}$$

$\Phi$ is the set of parameters (i.e. the DFE matrix $\mathbf{W}$ and mutation rate $\mu$), and $\pi_0 = 4N_e\mu$ is determined by the genome-wide drift-effective population size $N_e$, set by only reproductive and demographic processes. Regional mutation rate heterogeneity can be accounted for with a regional mutation rate scaling function $m(x)$, $\pi(x) = B(x, \Phi)m(x)\pi_0$, but we do not explore mutation-rate heterogeneity, as it is unclear how to differentiate variance in mutation rate from the substitution rate heterogeneity we find across the genome.

The likelihood of the set of parameters $\Psi = \{\pi_0, \Phi\}$ can be written in the form of,

$$\log \mathcal{L}(\Psi) = \sum_{v \in \mathcal{V}} \sum_{i \neq j \in \mathcal{S}} \log(P(O_{i,j}(v)|\Psi)) \tag{77}$$

(c.f. Elyashiv et al. 2016; McVicker et al. 2009; Murphy et al. 2022) where $\mathcal{V}$ is the set of putatively neutral sites, $\mathcal{S}$ is the set of samples, and $\Psi$ are the BGS parameters. The indicator variable $O_{i,j}(v)$ is 1 if samples $i$ and $j$ are different at putatively neutral site $v$, and zero otherwise. While the theoretic $\pi(v)$ gives the average number of pairwise differences, for small values, this is approximately the heterozygosity probability, so we can write

$$P(O_{i,j}(v)|\Psi) = \begin{cases} \pi(v), & O_{i,j}(v, \Psi) = 1 \\ 1 - \pi(v), & O_{i,j}(v, \Psi) = 0 \end{cases}. \tag{78}$$

(c.f. Elyashiv et al. 2016).

As in Section 2.6, the number of pairwise differences and the total number of pairwise comparison at a site are sufficient statistics for the likelihood. Then, the binomial log-likelihood for the data at site $v$ is,

$$\ell_v(\Psi) = \log(\pi(v, \Psi))n_{\mathrm{D},v} + \log(1 - \pi(v, \Psi))n_{\mathrm{S},v} \tag{79}$$

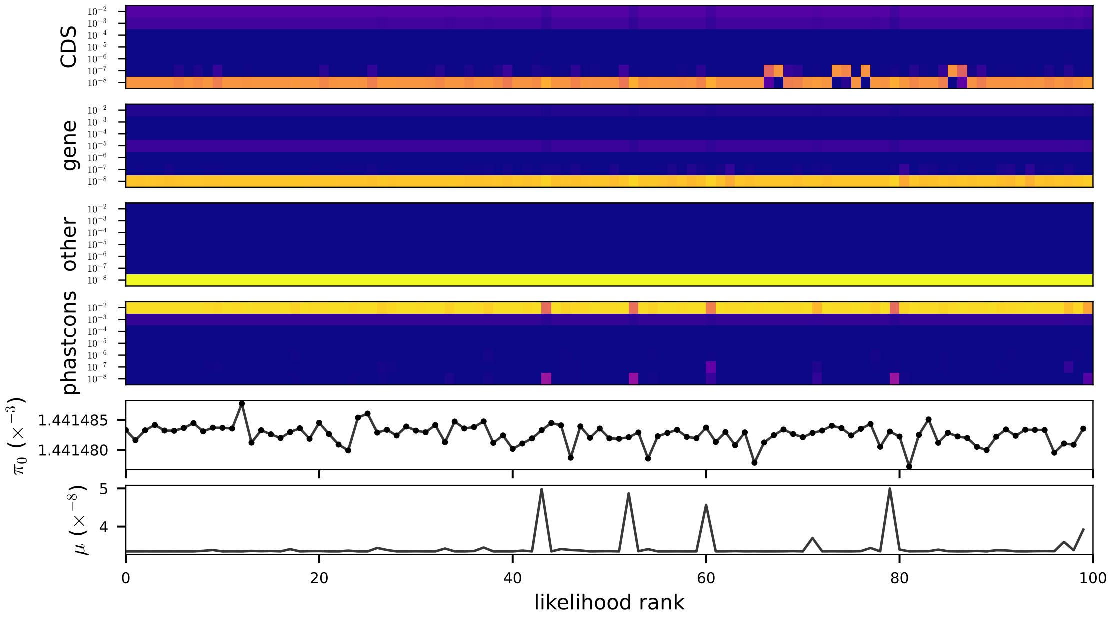

Figure 4: An optimization diagnostic plot showing the 100 top optima. Heatmap colors indicate probability mass for the DFE parameters (top four rows). Estimates of $\pi_0$ and $\mu$ are shown in the bottom two rows. This is the optimization plot for YRI Feature Priority model.

##### 3.12 Optimization and Diagnostics

As described in the Methods section we use the BOBYQA algorithm implemented by the NLopt library for numeric optimization to find maximum likelihood estimates (Johnson 2007; Powell 2009). The DFE is a simplex, and thus requires constrained optimization. We compared constrained optimization algorithms to an unconstrained softmax approach, finding the latter vastly outperformed the former. Bounds for $\pi_0$ and $\mu$ can be set manually in `bgspy`; our main models used $10^{-8} \leq \mu \leq 5 \times 10^{-8}$ and $0.0005 \leq \pi_0 \leq 0.008$. All main model fits were not at bounds. For locally rescaled fits, we found our $\widehat{\mu}$ estimate was continually the upper bound for the PhastCons Priority model; for these fits we increased this first to $\mu = 8 \times 10^{-8}$ and then again to $\mu = \times 10^{-7}$, find it still always reached the upper bound. This did not occur for CADD 6% locally rescaled models.

We only show one optimization diagnostic plot here (Supplementary Figure 4), but the can be easily calculated from the model pickle files in the Data Dryad repository (`https://doi.org/10.5061/dryad.qnk98sfnv`). In one case, our optimization diagnostics indicated that the best optima across 10,000 random starts was one of few of the top 100 hits that were hitting either an alternate likelihood mode, or the optimization routine had numeric issues. In this one case, we removed the top hit and used the second, which was the consensus among the other top 100 optima.

##### 3.13 The scale of processes

We can observe $\widehat{\pi}(x)$ at a per-basepair resolution. However, for a variety of reasons, we do not want to fit the composite likelihood model to the per-basepair scale of data. First, this would be computationally infeasible. Second, the mean function $\pi(x)$ varies on a natural scale that is itself a free parameter of the model. Our model can be written as,

$$\ell(\Psi, h) = \sum_b \sum_{v \in \mathcal{V}_b} \ell_v(\Psi) \tag{80}$$

$$= \sum_b \left[ \log(\bar{\pi}(b, \Psi)) \sum_{v \in \mathcal{V}_b} n_{\mathrm{D},v} + \log(1 - \bar{\pi}(b, \Psi)) \sum_{v \in \mathcal{V}_b} n_{\mathrm{S},v} \right] \tag{81}$$

$$= \sum_b \left[ \log(\bar{\pi}(b, \Psi)) Y_{\mathrm{D},b} + \log(1 - \bar{\pi}(b, \Psi)) Y_{\mathrm{S},b} \right] \tag{82}$$

$$\tag{83}$$

where $h$ is the bandwidth or window size, $b$ the bin index for windows of width $h$, $\mathcal{V}_b$ is the set of putatively neutral sites in bin $b$, $\bar{\pi}(b|\Psi)$ are the average diversity in bin $b$, and $Y_{\mathrm{S},b}$ and $Y_{\mathrm{S},b}$ are the sums across putatively neutral sites within a bin. Note that by binning, we sum the pairwise summaries of the data $\mathbf{Y}$ across sites, so the likelihood across bins is naturally weighted by the quantity of observed data.

This model corresponds to a binomial likelihood for the observed data summarized at genomic scale $h$. Thus an alternative way to express this model is as,

$$Y_{D,b} \sim \text{Binom}(\bar{\pi}(b, \Psi), Y_{D,b} + Y_{S,b}). \tag{84}$$

Here, $\bar{\pi}(b, \Psi)$ is assumed to be the *probability* of sampling two different alleles, rather than the average *number* of pairwise differences; these are approximately equal when $\pi$ is small. This corresponds to an identity link function; one could alternatively use a two-alleles finite sites model link function of the form, $\pi/(1 + 2\pi)$. We experimented with this and found there was little difference between these link functions, so we opted for the simpler identity link function.

##### 3.14 Out-Sample Error Estimation and $\widehat{R}^2_{\text{LOCO}}$

To estimate out-sample prediction error, we fit each model leaving one chromosome out (LOCO) and calculating the out-of-sample prediction error on these excluded chromosomes. Each fit used 10,000 random starting positions as the main fits. The out-sample prediction error (i.e. sum of residuals squared, $\widehat{\text{SSR}}_{\text{LOCO}}$) for all bins in chromosome $C$ left out is,

$$\widehat{\text{SSR}}_{\text{LOCO}}(C) = \sum_{b \in C} (\widehat{\pi}_{b,(C)} - \pi_b)^2 \tag{85}$$

where $\widehat{\pi}_{b,(C)}$ denotes the predictions for window $b$ fit on data excluding chromosome $C$. The average out-sample error $\widehat{\text{SSR}}_{\text{LOCO}}$ across chromosomes is then the average of $\widehat{\text{SSR}}_{\text{LOCO}}(C)$ across chromosomes, weighted by the number of bins per chromosome. We then calculate the out-sample coefficient of determination $\widehat{R}^2_{\text{LOCO}}$

$$\widehat{R}^2_{\text{LOCO}} = 1 - \frac{\widehat{\text{SSR}}_{\text{LOCO}}}{\widehat{\text{SST}}_{\text{LOCO}}} \tag{86}$$

where,

$$\widehat{\text{SST}}_{\text{LOCO}}(C) = \sum_{b \in C} (\widehat{\pi}_{b,(C)} - \bar{\pi})^2 \tag{87}$$

and $\bar{\pi}$ is the average diversity across all windows.

#### 3.15 Jackknife Procedures

We use a block jackknife procedure to calculate the uncertainty of our estimates. This approach is unfortunately computationally intensive, since a 10Mbp block width requires ≈ 280 blocks to cover 2.8Gbp of autosomal sequence and each fit requires thousands of random starts to reliably find consistent optima. We use a parallelization strategy where a set number of evenly-spaced positions along the genome that are chosen to approximately cover all 2.8Gbps of autosomal sequence. Each of these positions is sent out across a computing cluster, which excludes the 10Mbp block there and fits on the remaining data. Then, using these jackknife samples we calculated the jackknife standard error for each of parameters (Wasserman 2004).

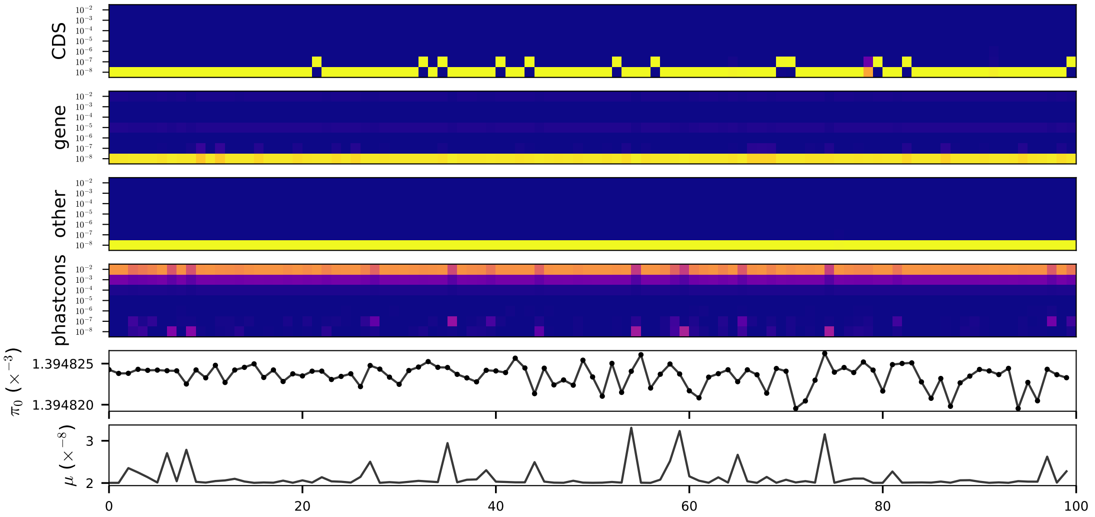

Figure 5: An optimization diagnostic plot showing an example of the bimodal likelihood optima, or a numeric optimization issue. The first four rows are the DFE estimates, where color indicates the parameter value (brighter colors closer to 1, darker colors closer to 0). The $\pi_0$ and $\mu$ estimates are shown in the bottom two rows. Note that different parameter combinations co-occur, indicating in the top 100 optima, there is an apparent other mode that varies with mutation rate.

In one case, for our PhastCons Priority model mutation rate estimtes, our jackknife standard error estimates were very large. Upon inspection of optimization diagnostic plots (see Supplementary Materials Section 5) and histograms of jackknife estimates, these looked like they were due to rare aberrant optima. These were either caused by a numeric optimization issue or the presence of multiple local optima. Given that this looked to be artificially inflating the uncertainty compared to all other models, we filtered out these jackknife results before computing the standard error.

### 4 Simulations

All of our forward simulations used SLiM version 4.0.1 (Haller and Messer 2019, 2023) and all Eidos scripts are available in the `slim_sims/` directory in the GitHub repository (`https://github.com/vsbuffalo/bprime`). These Eidos scripts take command line arguments for the parameters. Our simulation pipelines use YAML configuration files that specify the parameter grids and other settings, which are read and dispatched across a cluster using Snakemake (Köster and Rahmann 2012).

#### 4.1 Segment Simulations

To confirm the SC16 theory, we simulated a 100 kbp segment in a population of $N = 1000$ diploids across grids of mutation and selection parameters. The Eidos script for these simulations can be found at `slim_sims/region/region.slim` and the analysis notebook at `notebooks/region_simulations.ipynb` in the GitHub repository (`https://github.com/vsbuffalo/bprime`). We simulated over grids of mutation rates ($\mu \in \{10^{-9.5}, 10^{-9}, 10^{-8.5}, 10^{-8}\}$) and selection coefficients ($s \in \{10^{-1}, 10^{-1.5}, 10^{-2}, \ldots, 10^{-4.5}\}$). Selection was additive within a site and multiplicative across sites, and recombination was fixed at $10^{-8}$ per basepair.

We replicated $10,000$ simulations for each parameter combination to average and compare against theory. Each simulation recorded the number of fixations and the fitness variance per generation. At the end $10N$ generations, the ancestral recombination graph was recorded. Then, downstream Python scripts calculate branch-length diversity using `tskit` (Kelleher et al. 2018), and we estimate the reduction factor in a window centered in the region with $B = \pi/4N$ (since branch length diversity implies $\mu = 1$).

#### 4.2 Whole-chromosome Simulations

Our whole-chromosome simulations use realistic human recombination maps and putatively conserved segments. These annotation differ slightly from our primary model fits, since these simulations were written before the method was finalized. We use the HapMap recombination map (International HapMap Consortium et al. 2007), PhastCons regions (Siepel et al. 2005), and UTRs as an approximate set of putatively conserved segments.

Our first set of simulations were used to validate that our methods were producing accurate classic B and B' maps. We replicated 100 whole-chromosome simulations over grids of mutation rates ($\mu \in \{10^{-10}, 10^{-9.5}, \ldots, 10^{-8}, 10^{-7.5}\}$) and selection coefficients ($s \in \{10^{-1}, 10^{-1.5}, 10^{-2}, \ldots, 10^{-6}\}$). For each parameter combination, we calculate mean branch diversity in 10 kbp windows across the 100 replicates to create the average simulation reduction map.

We then compare this average simulation map to our method's theoretic B' maps (see Figure 1 D and E). We calculate the MSE per 10 kbp window to estimate the total rate across the chromosome.

We can assess the accuracy relative to a theoretic lower bound of the MSE if the noise were due to coalescence noise. We approximate this by noting that the MSE can be decomposed into a squared bias term and a variance term. If our methods are unbiased in estimating the true reduction at a position, the MSE is just the stochastic variance due to the $n = 100$ replicates we simulate and the coalescence noise in each one. Tajima (1983) showed that variance in diversity due to coalescence noise is much larger than sampling noise, and is given by $\mathrm{V}(\pi) \approx 2\theta^2/9$ in an infinite sample at a non-recombining locus. One can derive an analogous equation for the variance of average simulation $\bar{B}$ across $n$ replicate simulations as, $\mathrm{V}(\bar{B}) \approx 2B^2/9n$. We note that this assumes no recombination but this approximation is fine over short scales (such as the 10 kbp scale used here).

### 5 Additional Results and Figures

#### 5.1 Locally Rescaled Fit Results

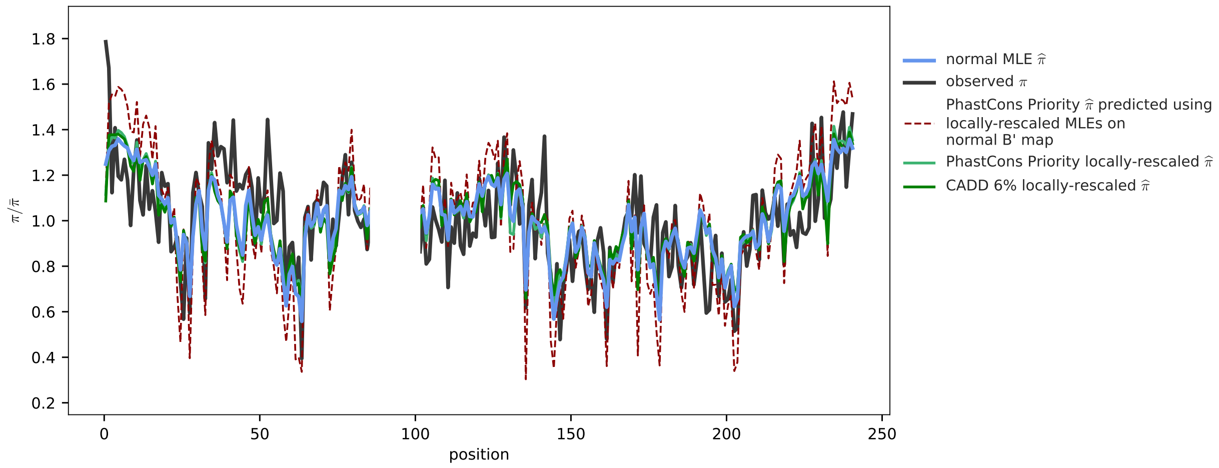

Figure 6: Chromosome 2 predictions on the YRI samples, with locally rescaled model fits. The observed data is the dark gray line, and the normal MLE for the PhastCons Priority model is the blue line. The locally rescaled predictions are the green line. The dashed red line are the prediction using the standard B' map (without local rescaling) and the maximum likelihood estimates from the locally rescaled fits. The large discrepancy in this shows that estimates are highly dependent on the B' map.

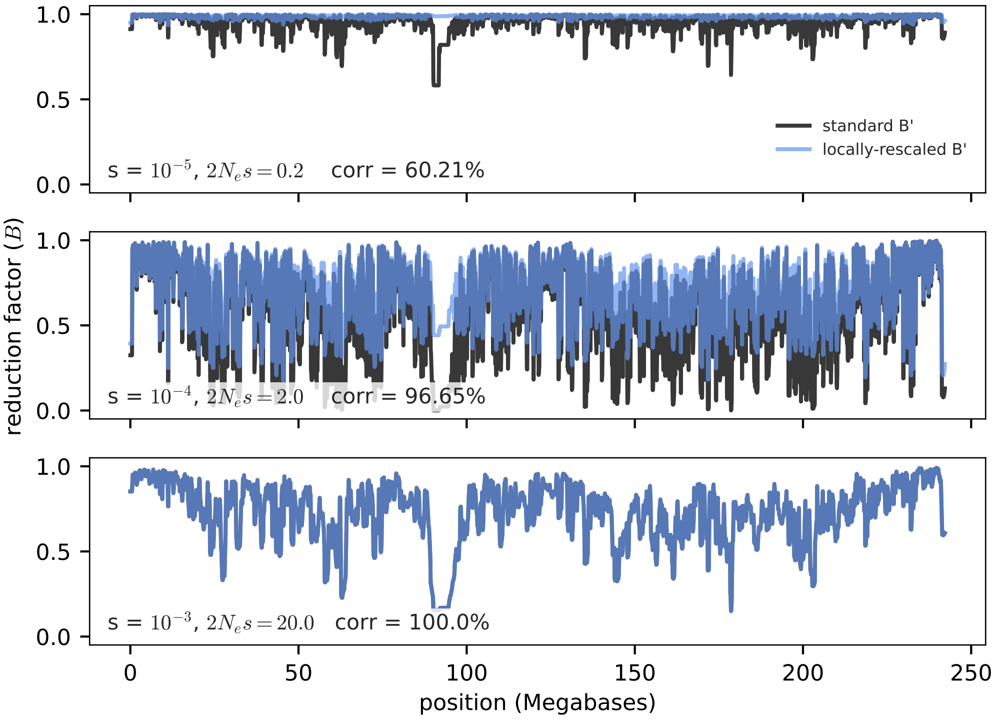

Figure 7: The standard (dark gray) and locally rescaled (blue) B' maps for different $\mu = 1.6 \times 10^{-8}$ and three different selection coefficients. For $s = 10^{-5}$ ($2N_e s = 0.2$), locally rescaling alters the predicted reduction so that it is essentially insignificant ($B \approx 1$). For mid-strength selection $s = 10^{-4}$ ($2N_e s = 2$), there is only a very slight difference between standard and locally rescaled B' maps. Finally, for strong selection $s = 10^{-3}$ ($2N_e s = 20$), local rescaling does not change the B' maps, as expected.

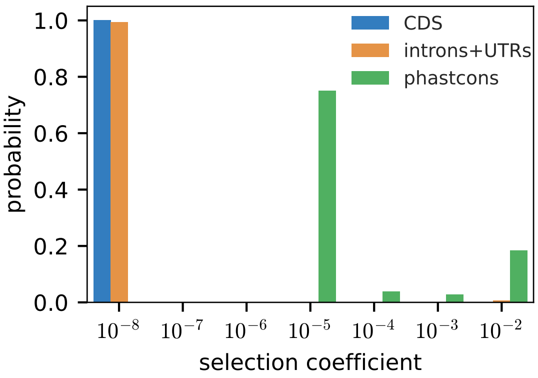

Figure 8: The DFE for YRI samples with local rescaling. The maximum likelihood mutation rate estimate for this is $\widehat{\mu} = 8 \times 10^{-8}$, which is the upper boundary of the range used during optimization.

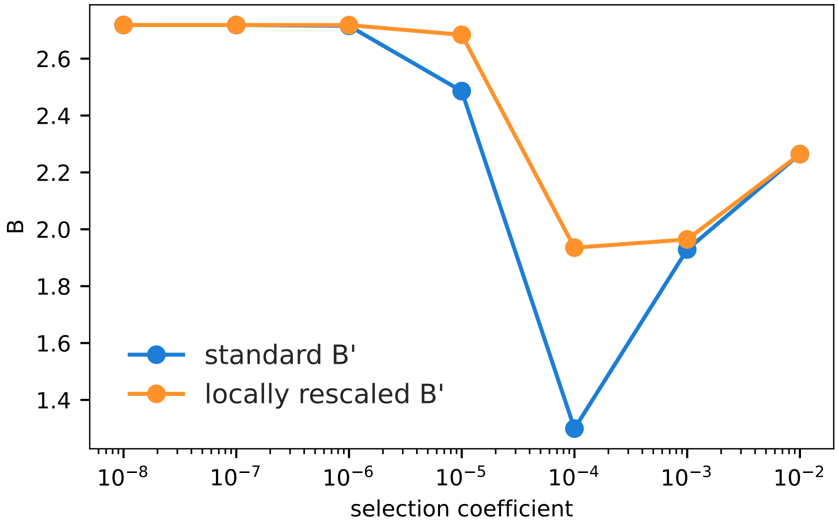

Figure 9: Genome-wide average $B(x)$ values across selection coefficients for $\mu = 1.58 \times 10^{-8}$, for both the standard (blue) and locally rescaled B' (orange) maps. This indicates that locally rescaling the B' maps only in practice changes how deep the "U" is.

##### 5.1.1 PhastCons Priority Locally Rescaled Model Fit Results

Below are the model fit results for locally rescaled model runs, with the upper mutation rate boundary set to $\mu = 8 \times 10^{-8}$. Each model fit is the direct output from a `bgspy`. The `w grid` and `t grid` are the grids the B and B' maps are built on; note that the mutation grid `w` is an exponential that is inferred later, so optimization is done continuously over this. Each model output shows the maximum likelihood estimates, with uncertainty where available.

#### CHB

```
SimplexModel (interpolated w): 6 x 7 x 3
    w grid: [1.000e-11 6.310e-11 3.981e-10 2.512e-09 1.585e-08 1.000e-07] (before interpolation)
    t grid: [1.e-08 1.e-07 1.e-06 1.e-05 1.e-04 1.e-03 1.e-02]
    window size: 1.0 Mbp

  ML estimates: (whole genome)
  standard error method: None
  negative log-likelihood: 61820142829.01206
  number of successful starts: 10000 (100.0% total)
  pi_0 = 0.00106289
  pi  = 0.000752601
  mut. rate  = 8e-08
  Ne = 26,572 (if mu=1e-8), Ne = 13,286 (if mu=2e-8)
  R² = 60.8648% (in-sample)  59.1654% (out-sample)
  W =
            CDS    gene    phastcons
  ------  -----  ------  -----------
  1e-08       1   0.991        0
  1e-07       0   0            0
  1e-06       0   0            0
  1e-05       0   0            0.767
  0.0001      0   0            0.03
  0.001       0   0            0.069
  0.01        0   0.008        0.134
```

### CEU

SimplexModel (interpolated w): 6 x 7 x 3

w grid: [1.000e-11 6.310e-11 3.981e-10 2.512e-09 1.585e-08 1.000e-07] (before interpolation)

t grid: [1.e-08 1.e-07 1.e-06 1.e-05 1.e-04 1.e-03 1.e-02]

window size: 1.0 Mbp

ML estimates: (whole genome)

standard error method: None

negative log-likelihood: 250647896892.40094

number of successful starts: 10000 (100.0% total)

pi_0 = 0.001479

pi  = 0.00106447

mut. rate  = 8e-08

Ne = 36,974 (if mu=1e-8), Ne = 18,487 (if mu=2e-8)

$R^2$ = 68.7046% (in-sample)  67.8854% (out-sample)

W =

| | CDS | gene | phastcons |
|---|---|---|---|
| 1e-08 | 1 | 0.994 | 0 |
| 1e-07 | 0 | 0 | 0 |
| 1e-06 | 0 | 0 | 0 |
| 1e-05 | 0 | 0 | 0.751 |
| 0.0001 | 0 | 0 | 0.039 |
| 0.001 | 0 | 0 | 0.028 |
| 0.01 | 0 | 0.006 | 0.183 |

### YRI

SimplexModel (interpolated w): 6 x 7 x 3

w grid: [1.000e-11 6.310e-11 3.981e-10 2.512e-09 1.585e-08 1.000e-07] (before interpolation)

t grid: [1.e-08 1.e-07 1.e-06 1.e-05 1.e-04 1.e-03 1.e-02]

window size: 1.0 Mbp

ML estimates: (whole genome)

standard error method: None

negative log-likelihood: 197824525643.2897

number of successful starts: 10000 (100.0% total)

pi_0 = 0.00111021

pi = 0.000801495

mut. rate = 8e-08

Ne = 27,755 (if mu=1e-8), Ne = 13,877 (if mu=2e-8)

$R^2$ = 62.5591% (in-sample) 61.367% (out-sample)

W =

| | CDS | gene | phastcons |
|---|---|---|---|
| 1e-08 | 1 | 0.989 | 0 |
| 1e-07 | 0 | 0.003 | 0 |
| 1e-06 | 0 | 0 | 0 |
| 1e-05 | 0 | 0 | 0.774 |
| 0.0001 | 0 | 0 | 0.025 |
| 0.001 | 0 | 0 | 0.067 |
| 0.01 | 0 | 0.008 | 0.134 |

##### 5.1.2 CADD 6% Locally Rescaled Model Fit Results

#### CHB

```
SimplexModel (interpolated w): 6 x 7 x 1
  w grid: [1.000e-11 6.310e-11 3.981e-10 2.512e-09 1.585e-08 1.000e-07] (before interpolation)
  t grid: [1.e-08 1.e-07 1.e-06 1.e-05 1.e-04 1.e-03 1.e-02]
  window size: 1.0 Mbp

ML estimates: (whole genome)
standard error method: None
negative log-likelihood: 61820130669.36475
number of successful starts: 10000 (100.0% total)
pi_0 = 0.00102026
pi  = 0.000752601
mut. rate  = 7.291e-08
Ne = 25,506 (if mu=1e-8), Ne = 12,753 (if mu=2e-8)
R² = 60.6057% (in-sample)
W =
          cadd6
------  -------
1e-08     0.009
1e-07     0
1e-06     0
1e-05     0.749
0.0001    0.014
0.001     0.067
0.01      0.161
```

#### CEU

```
SimplexModel (interpolated w): 6 x 7 x 1
  w grid: [1.000e-11 6.310e-11 3.981e-10 2.512e-09 1.585e-08 1.000e-07] (before interpolation)
  t grid: [1.e-08 1.e-07 1.e-06 1.e-05 1.e-04 1.e-03 1.e-02]
  window size: 1.0 Mbp

ML estimates: (whole genome)
standard error method: None
negative log-likelihood: 197825902519.15137
number of successful starts: 10000 (100.0% total)
pi_0 = 0.00107426
pi0  = 0.000801495
mut. rate  = 5.991e-08
Ne = 26,856 (if mu=1e-8), Ne = 13,428 (if mu=2e-8)
R² = 62.0668% (in-sample)
W =
          cadd6
------  -------
1e-08     0.001
1e-07     0
1e-06     0
1e-05     0.71
0.0001    0.027
0.001     0.065
0.01      0.197
```

#### YRI

SimplexModel (interpolated w): 6 x 7 x 1
  w grid: [1.000e-11 6.310e-11 3.981e-10 2.512e-09 1.585e-08 1.000e-07] (before interpolation)
  t grid: [1.e-08 1.e-07 1.e-06 1.e-05 1.e-04 1.e-03 1.e-02]
  window size: 1.0 Mbp

ML estimates: (whole genome)
standard error method: None
negative log-likelihood: 250648005868.94275
number of successful starts: 10000 (100.0% total)
pi0 = 0.0014366
pi  = 0.00106447
mut. rate  = 6.024e-08
Ne = 35,914 (if mu=1e-8), Ne = 17,957 (if mu=2e-8)
$R^2$ = 68.6747% (in-sample)
W =

| | cadd6 |
|---|---|
| 1e-08 | 0 |
| 1e-07 | 0 |
| 1e-06 | 0 |
| 1e-05 | 0.707 |
| 0.0001 | 0.048 |
| 0.001 | 0.012 |
| 0.01 | 0.233 |

#### 5.2 Strong Selection Grids

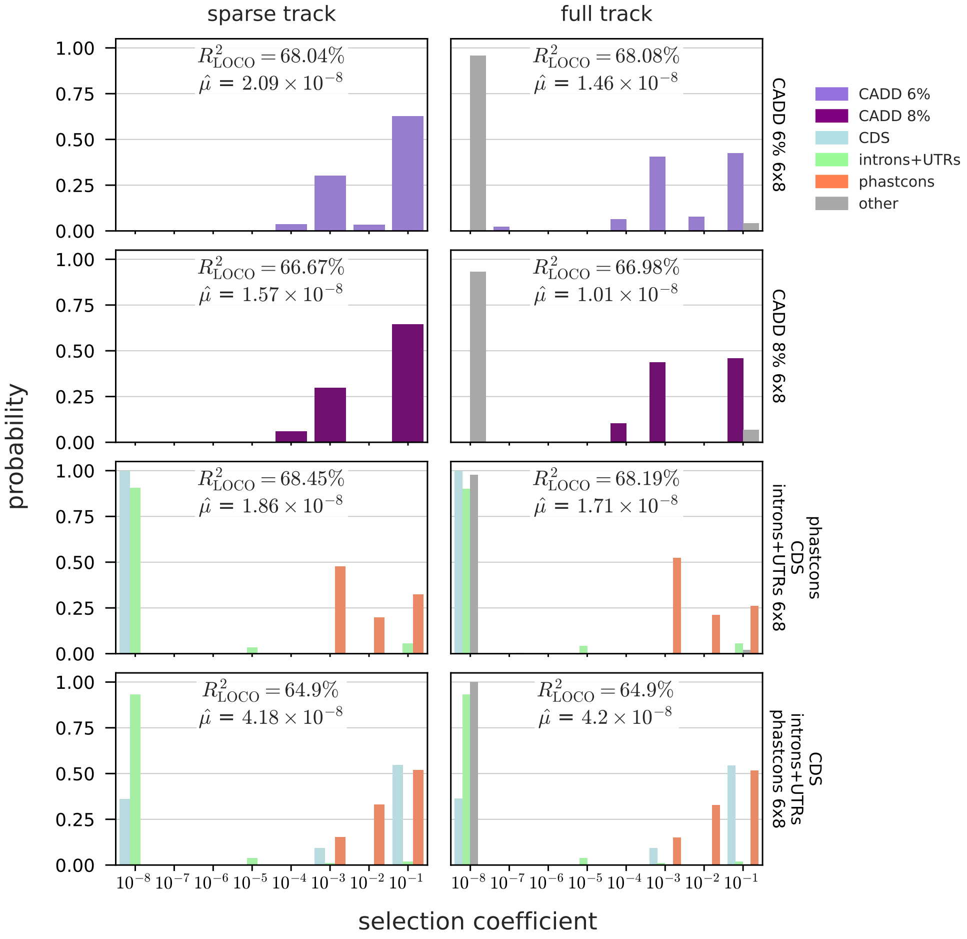

Figure 10: The DFE estimates for the strong selection grid (up to $s = 10^{-1}$).

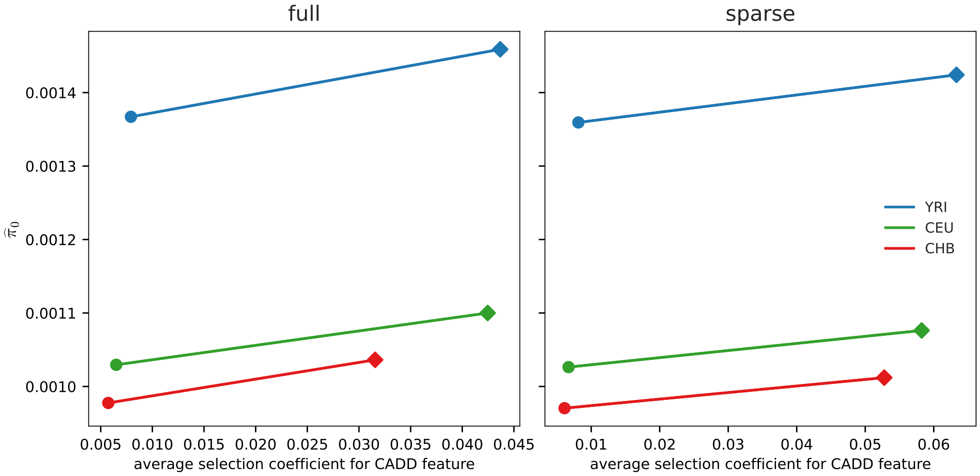

Figure 11: The maximum likelihood estimates of $\pi_0$ and average selection coefficient implied by the estimated DFE for the CADD 6% models. Diamonds indicate estimates under the strong selection grid (up to $s = 10^{-1}$) and circles indicate estimates under the default grid (up to $s = 10^{-2}$).

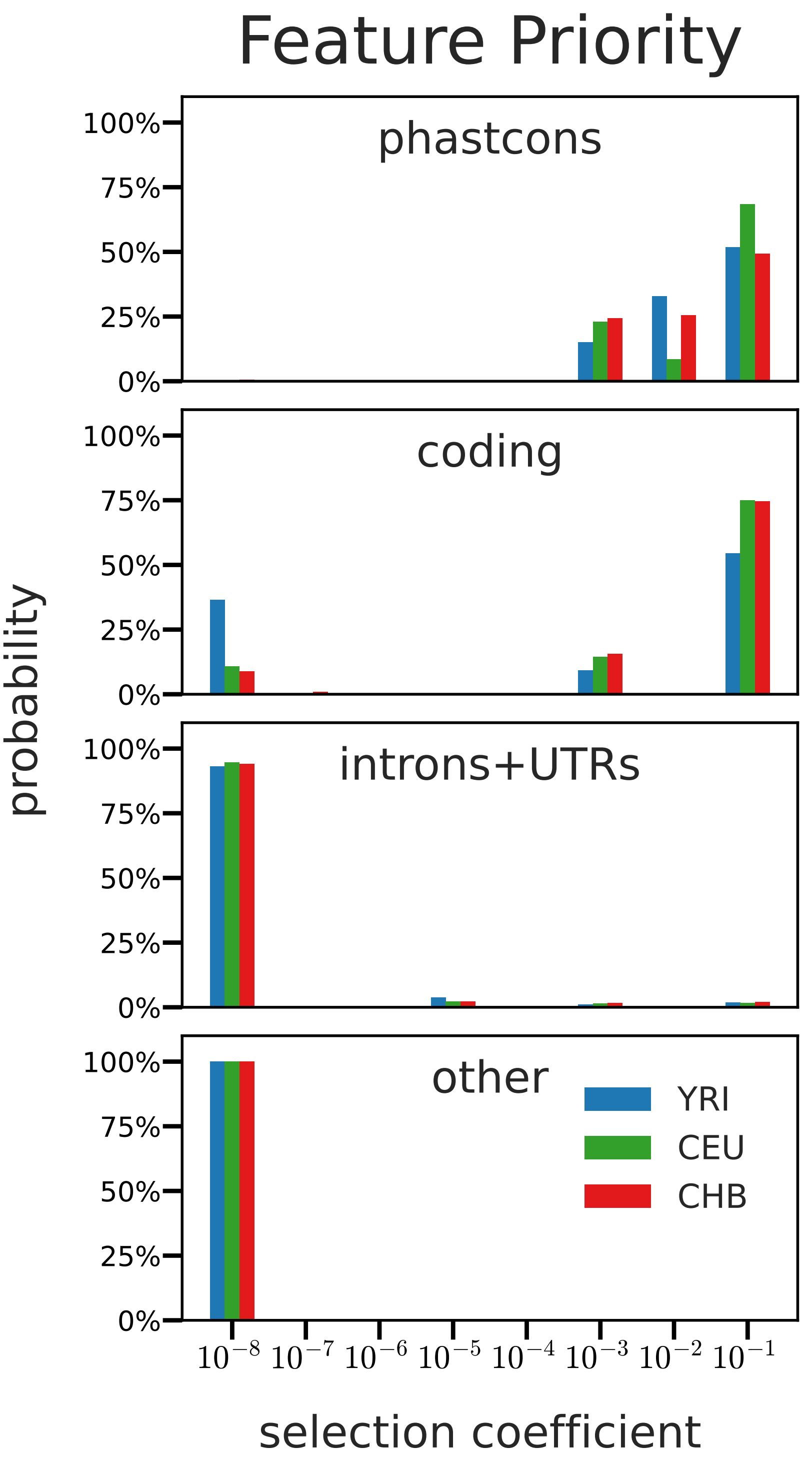

Figure 12: The DFE estimates for the strong selection grid (up to $s = 10^{-1}$) under the Feature Priority model, across populations.

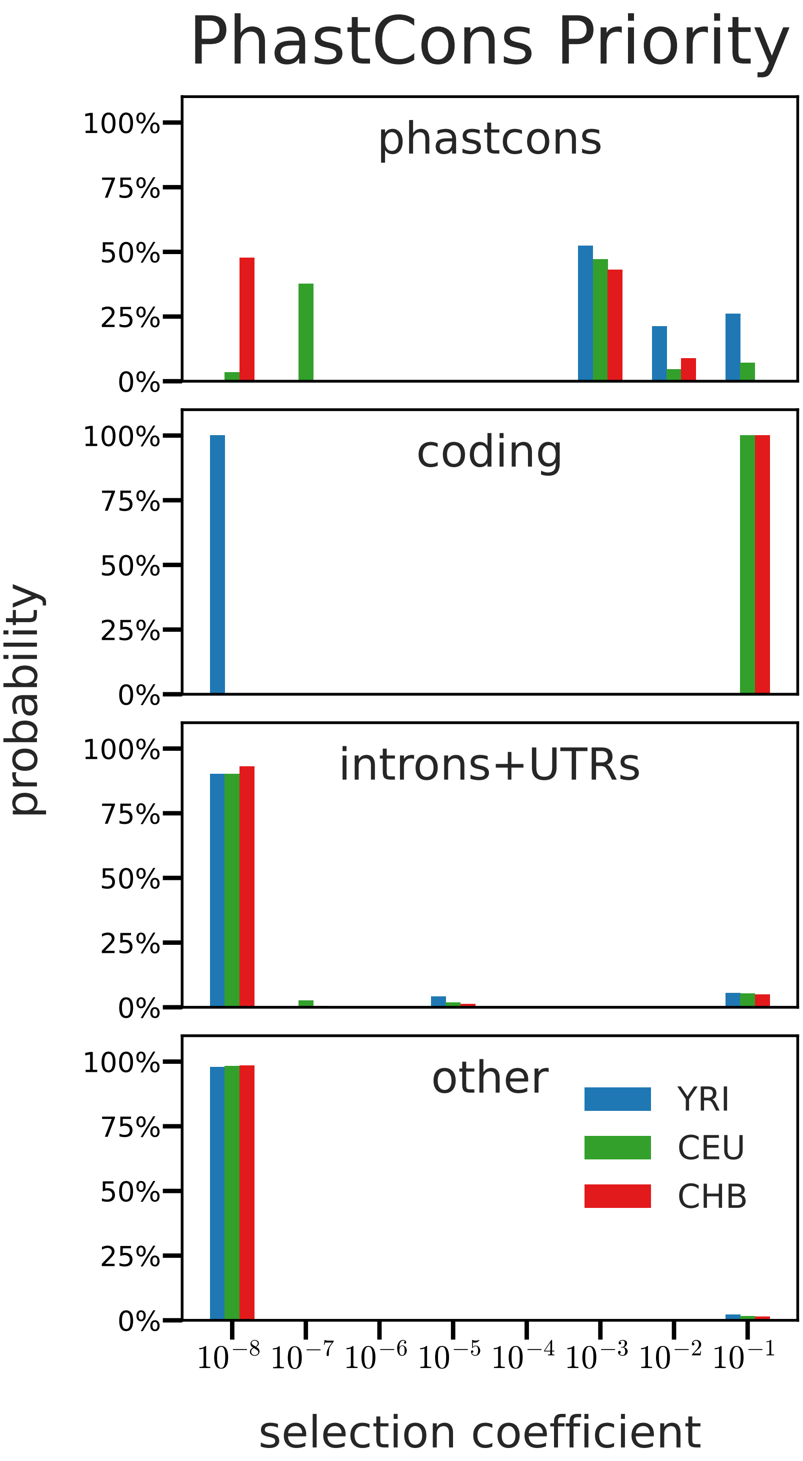

Figure 13: The DFE estimates for the strong selection grid (up to $s = 10^{-1}$) under the PhastCons Priority model, across populations.

##### 5.3 Demographic Expansion and Recessive Simulation Tests

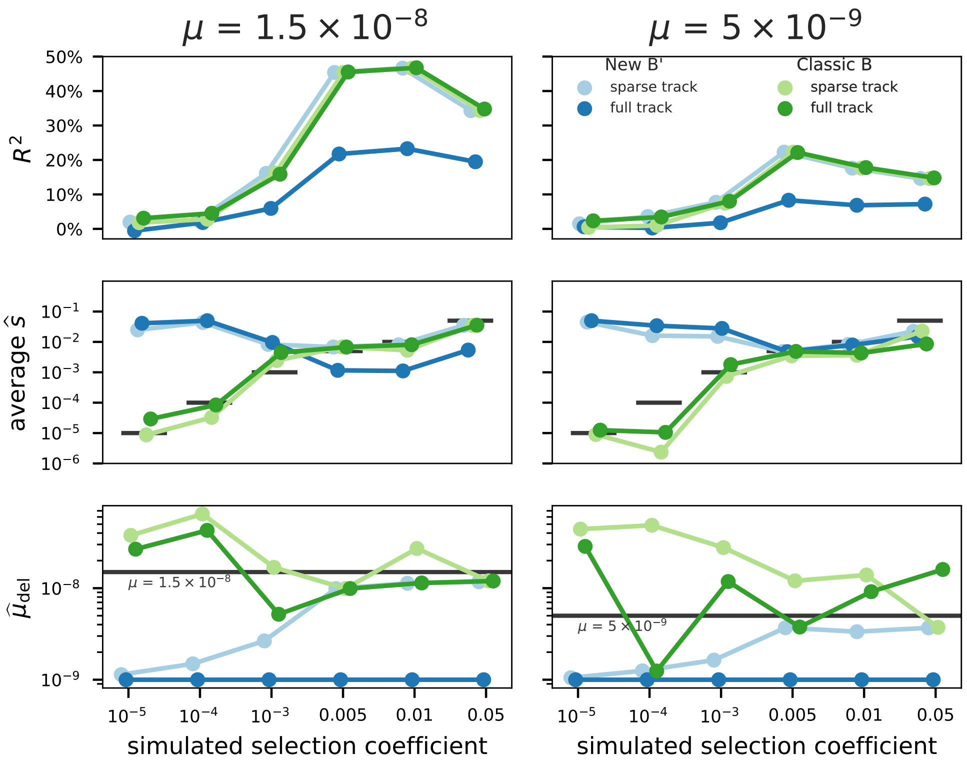

Figure 14: Simulations to diagnose the effect of population exponential expansion on our composite likelihood estimates. The simulated population expansion occurs at a rate of 0.4% a generation, starting $9.3N$ (out of $10N$) generations in. These parameters were meant to approximate the European expansion parameters estimated by Gutenkunst et al. (2009). Overall, we see little impact on estimated parameters.

##### 5.4 Average Selection Coefficients

There are the average selection coefficients ($\bar{s}$) calculated from the estimated DFEs, across both default and constrained grids.

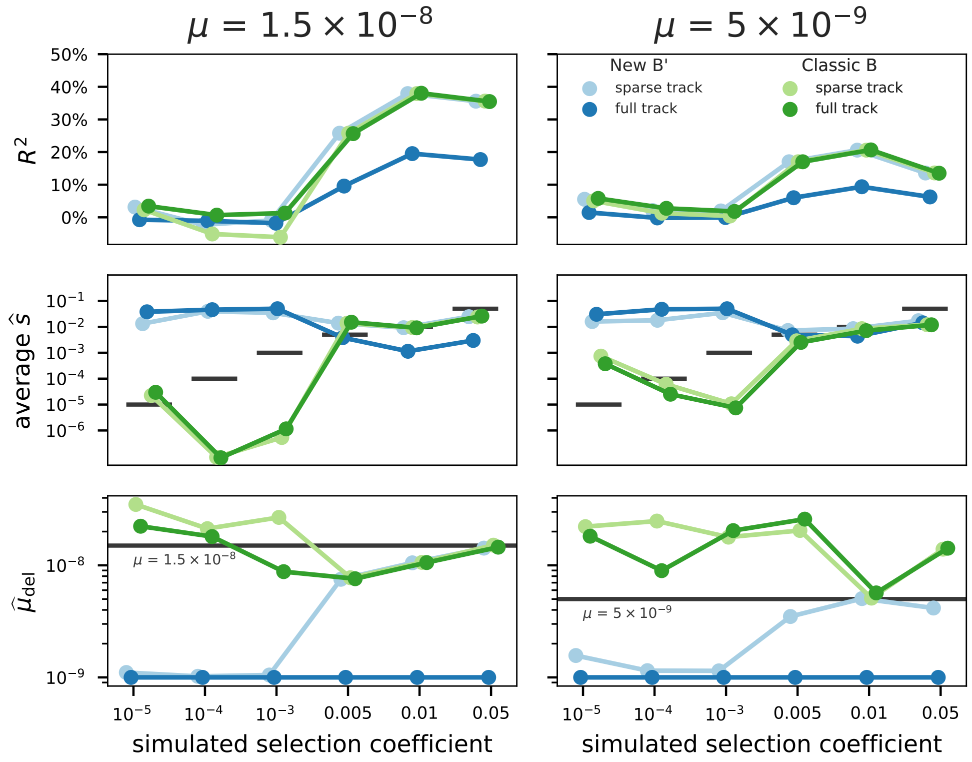

Figure 15: Simulations to diagnose the effect of non-additive mutations. These simulations use a dominance coefficient of $h = 0.3$, i.e. the selective effects are 1 ancestral state, $1 + hs$ for heterozygotes, and $1 + s$ for homozygotes. We find that slightly recessive mutations have little effect on the estimated selection coefficient when selection is strong, consistent with the fact that only the selective effect in heterozygotes matters when mutations are kept at low frequency by strong selection. However, our B' method underestimates weak selection coefficients when mutation effects are recessive. This is because recessive mutations are removed far more efficiently once they reach appreciable frequencies and are found in heterozygotes. Since our theory assumes additive effects, it accommodates by estimating weaker selection coefficients.

| Annotation Model | Population | Feature | $\bar{s}$ (default grid) | $\bar{s}$ (constrained grid) |
|---|---|---|---|---|
| PhastCons Priority | CEU | CDS | 0.100000 | 0.000000 |
| PhastCons Priority | CEU | gene | 0.005308 | 0.000226 |
| PhastCons Priority | CEU | phastcons | 0.008076 | 0.006278 |
| PhastCons Priority | CHB | CDS | 0.100000 | 0.000000 |
| PhastCons Priority | CHB | gene | 0.004896 | 0.000296 |
| PhastCons Priority | CHB | phastcons | 0.001327 | 0.005887 |
| PhastCons Priority | YRI | CDS | 0.000000 | 0.000000 |
| PhastCons Priority | YRI | gene | 0.005556 | 0.000163 |
| PhastCons Priority | YRI | phastcons | 0.028642 | 0.007687 |
| Feature Priority | CEU | CDS | 0.075041 | 0.005284 |
| Feature Priority | CEU | gene | 0.001654 | 0.000315 |
| Feature Priority | CEU | phastcons | 0.069563 | 0.008772 |
| Feature Priority | CHB | CDS | 0.074797 | 0.005211 |
| Feature Priority | CHB | gene | 0.001974 | 0.000271 |
| Feature Priority | CHB | phastcons | 0.051988 | 0.008507 |
| Feature Priority | YRI | CDS | 0.054503 | 0.001576 |
| Feature Priority | YRI | gene | 0.001912 | 0.000336 |
| Feature Priority | YRI | phastcons | 0.055153 | 0.009370 |
| CADD 6% | CEU | cadd6 | 0.042458 | 0.006500 |
| CADD 6% | CHB | cadd6 | 0.031565 | 0.005733 |
| CADD 6% | YRI | cadd6 | 0.043671 | 0.007926 |
| CADD 8% | CEU | cadd8 | 0.041254 | 0.006017 |
| CADD 8% | CHB | cadd8 | 0.029052 | 0.005068 |
| CADD 8% | YRI | cadd8 | 0.046212 | 0.007543 |

#### 5.5 Residual Covariate Diagnostics

#### 5.6 Additional DFE Figures

#### 5.7 Spatial Residuals

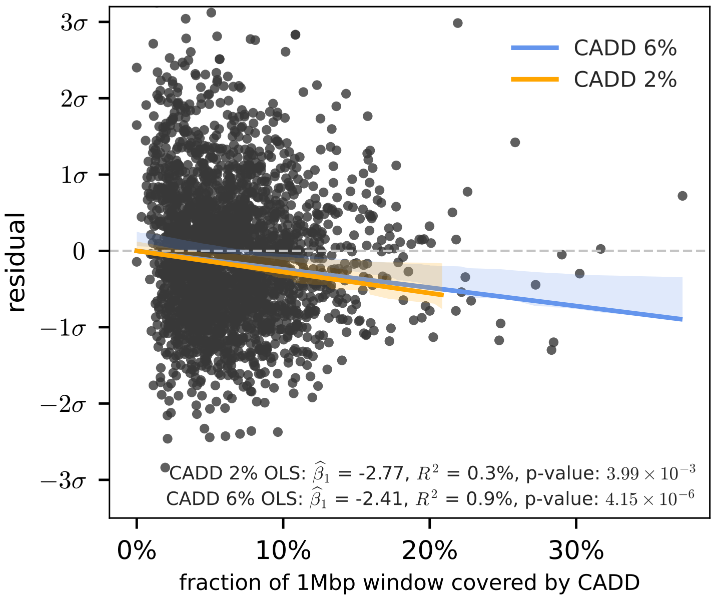

Figure 16: Residuals from the CADD 6% sparse track plotted against the fraction of basepairs in a window annotated by a CADD region.

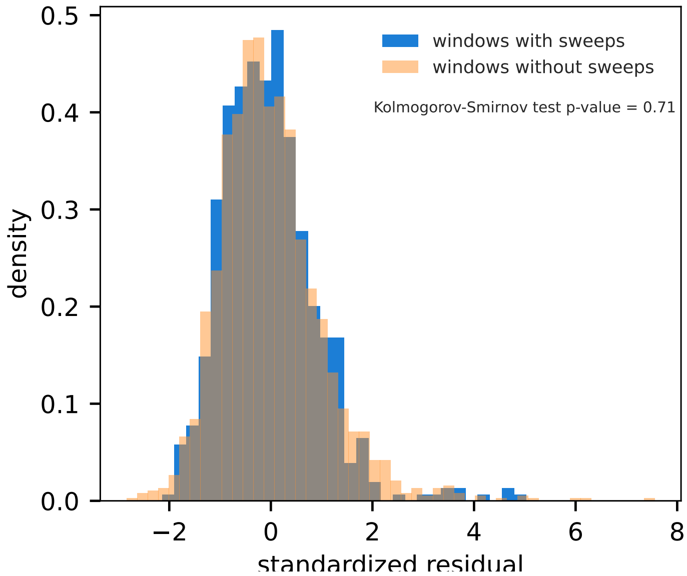

Figure 17: The distributions of residuals in windows containing hard or soft sweeps (blue) and not containing sweeps (orange) found by (Schrider and Kern 2017).

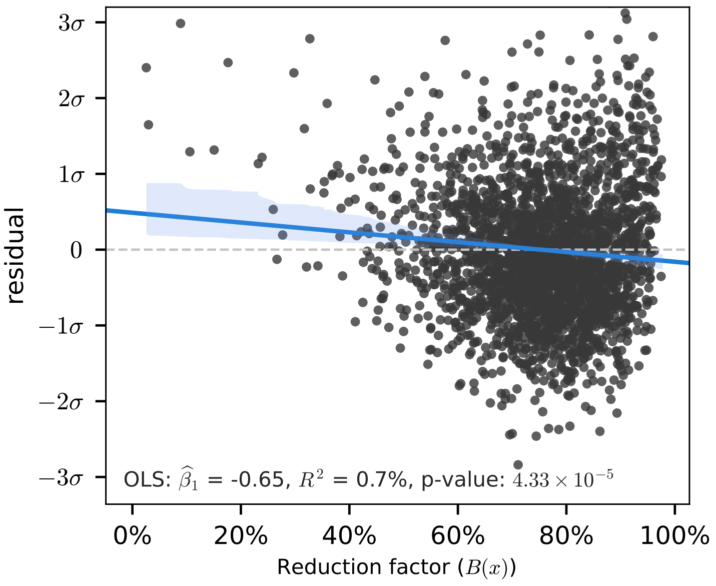

Figure 18: Residuals from the CADD 6% model plotted against the predicted $B(x)$ value.

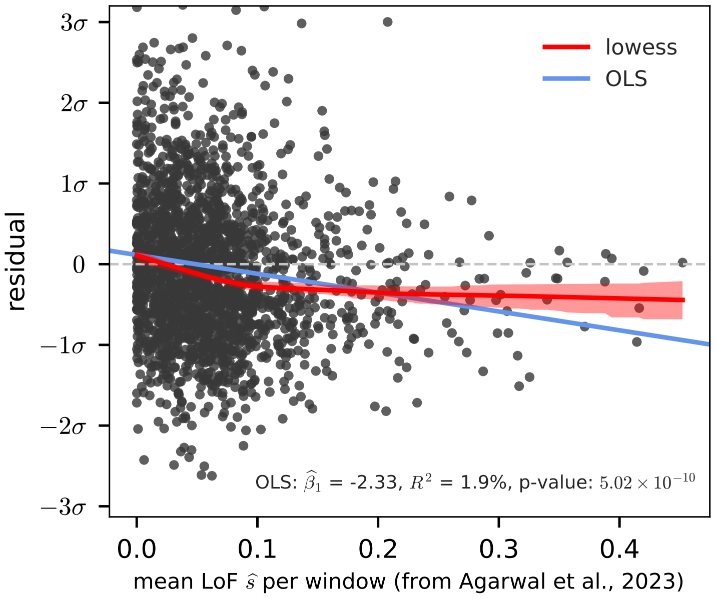

Figure 19: Residuals from the PhastCons Priority model plotted against the average LoF selection coefficient across genes in a megabase window (estimated from Agarwal et al. 2023). These results show that the remaining likely selection signal is not dependent on the model used (PhastCons Priority versus CADD 6%).

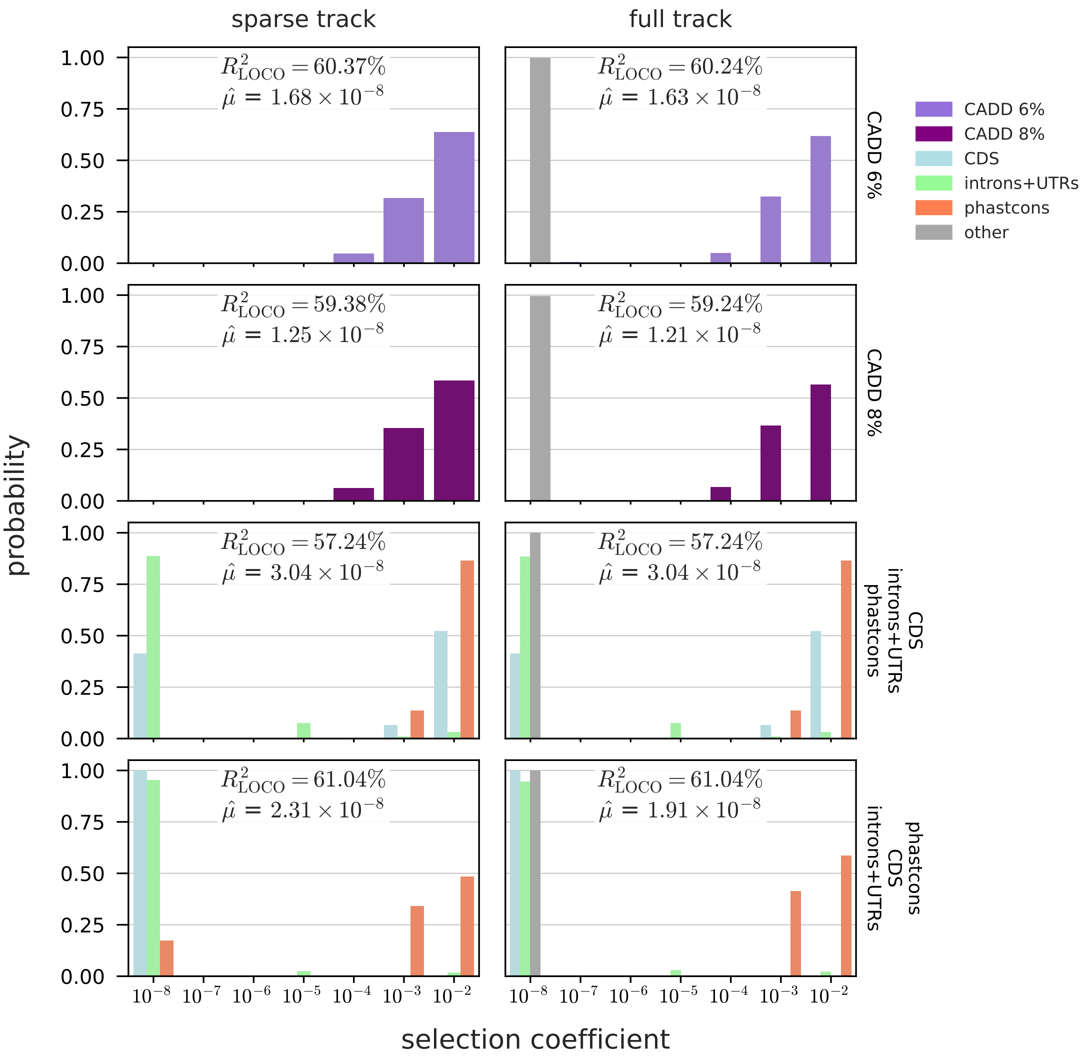

Figure 20: The DFE across models for the CEU reference sample (analogous to the DFE for YRI samples in Figure 3.

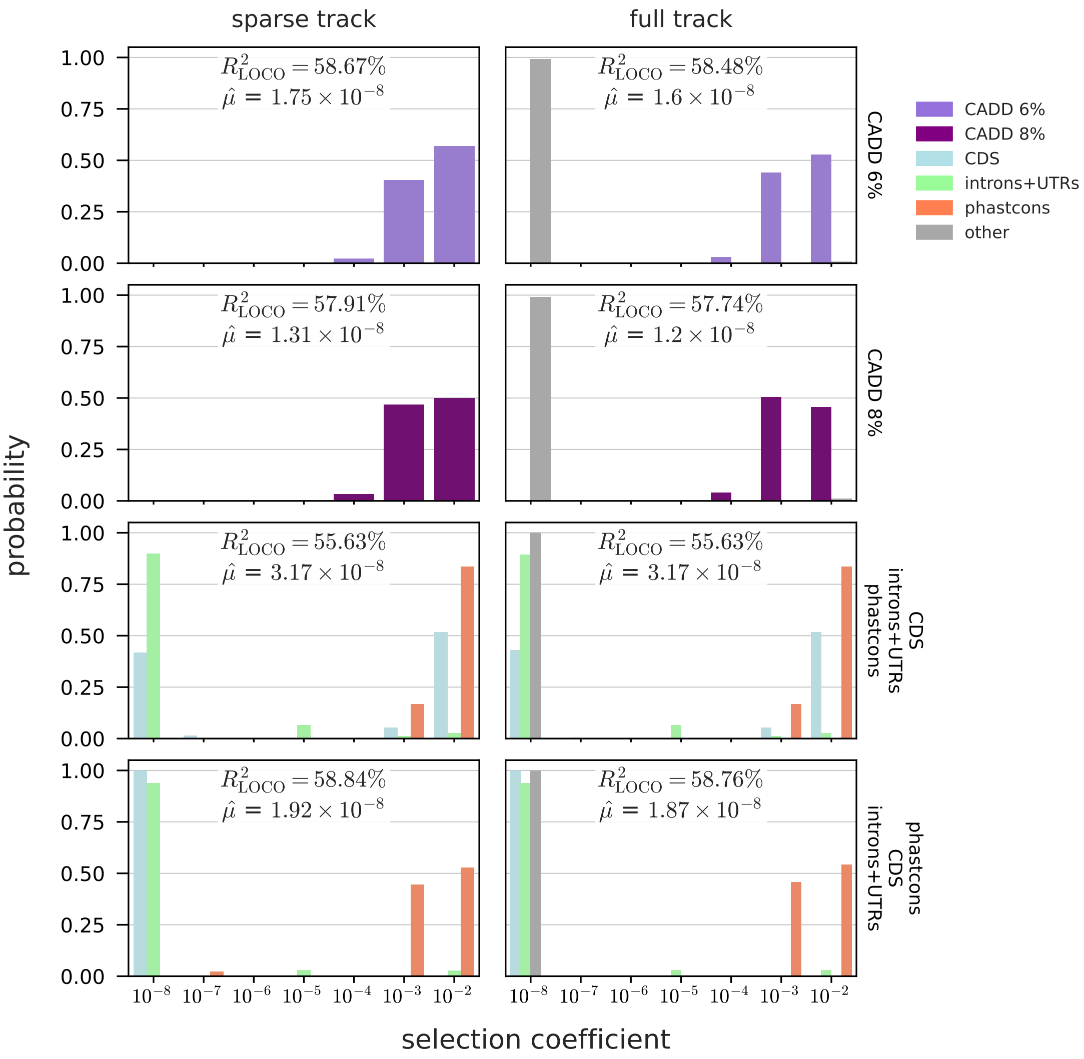

Figure 21: The DFE across models for the CHB reference sample (analogous to the DFE for YRI samples in Figure 3.

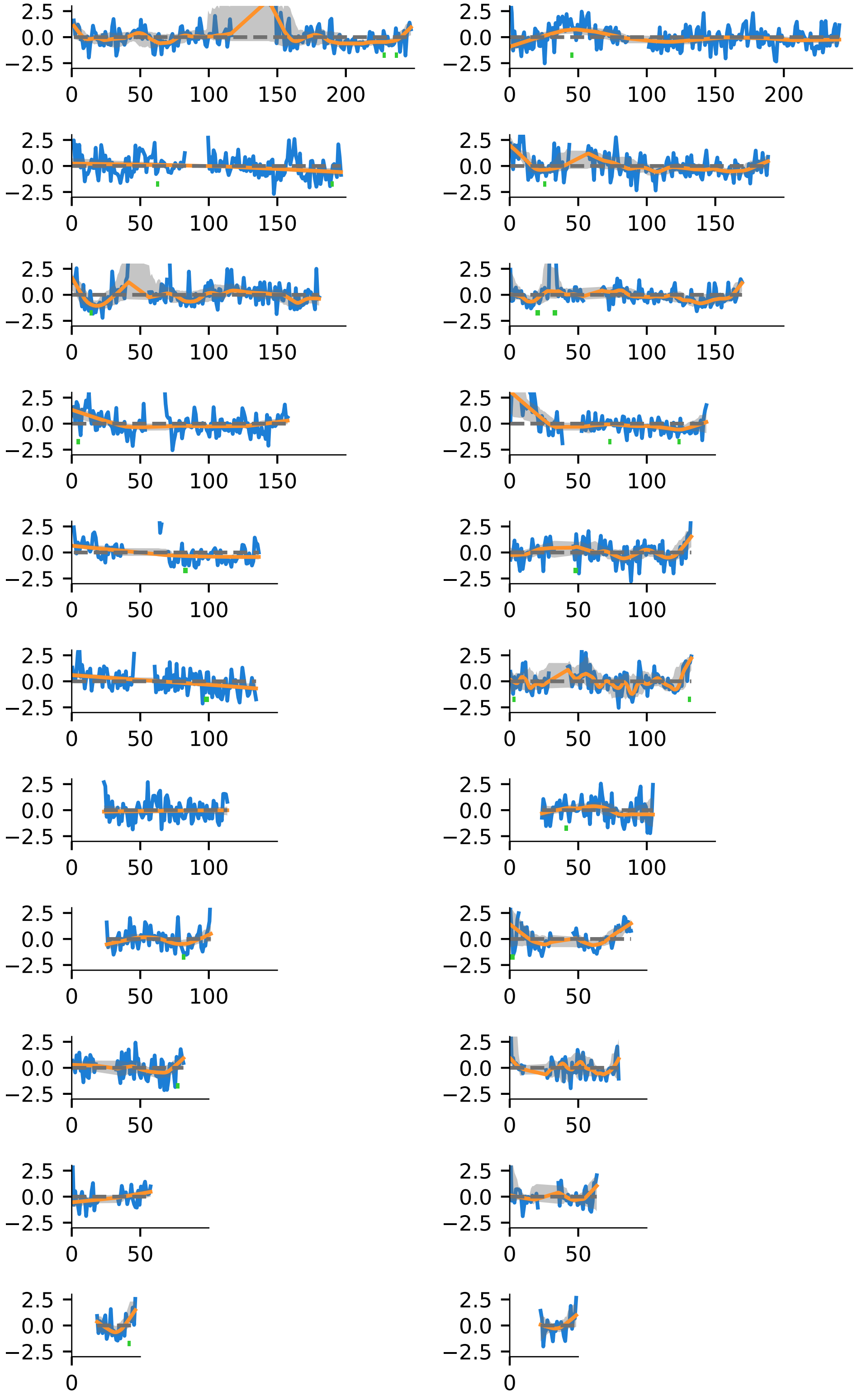

Figure 22: Blue lines represent the residual between predicted an observed diversity levels along chromosomes. The orange line is a lowess curve, with bandwidth chosen by cross-validation.

#### 5.8 Chromosome Fit Figures

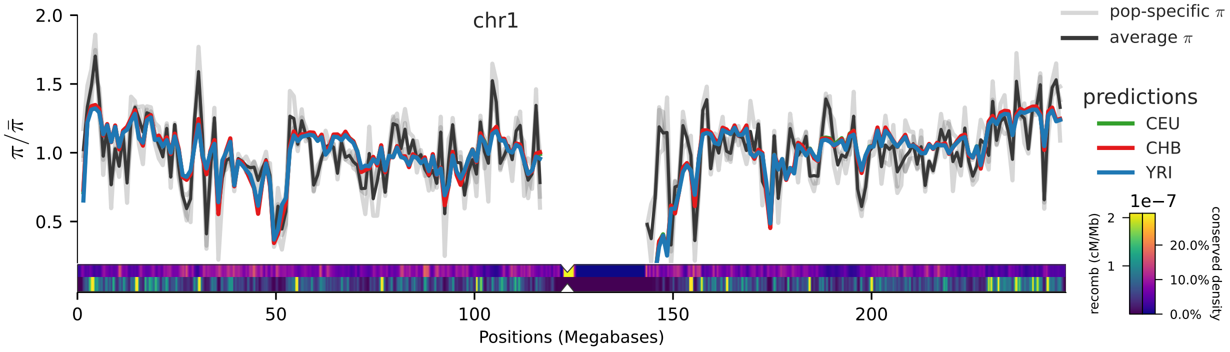

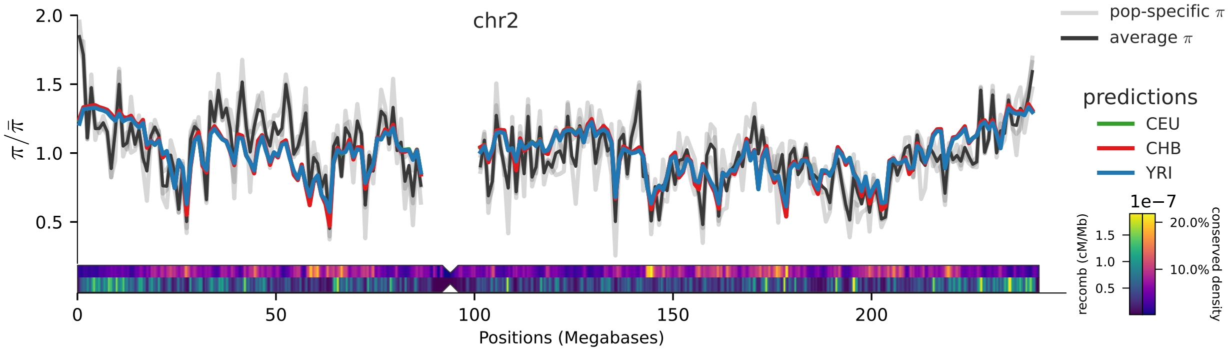

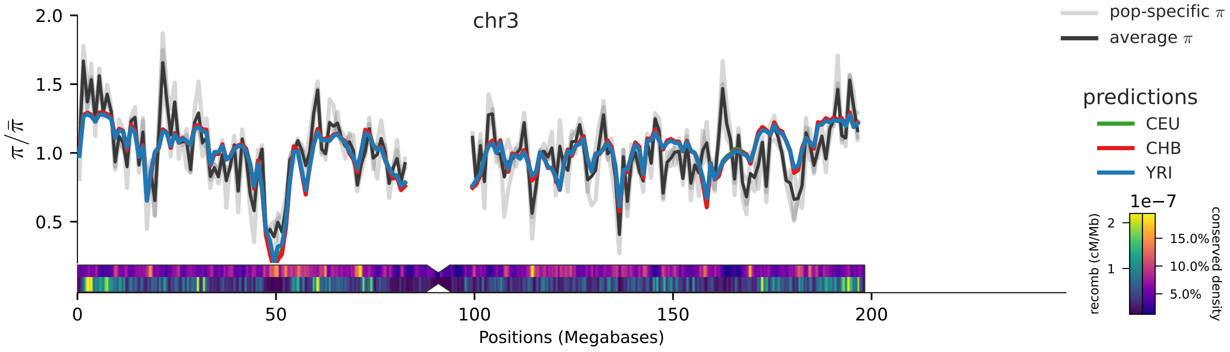

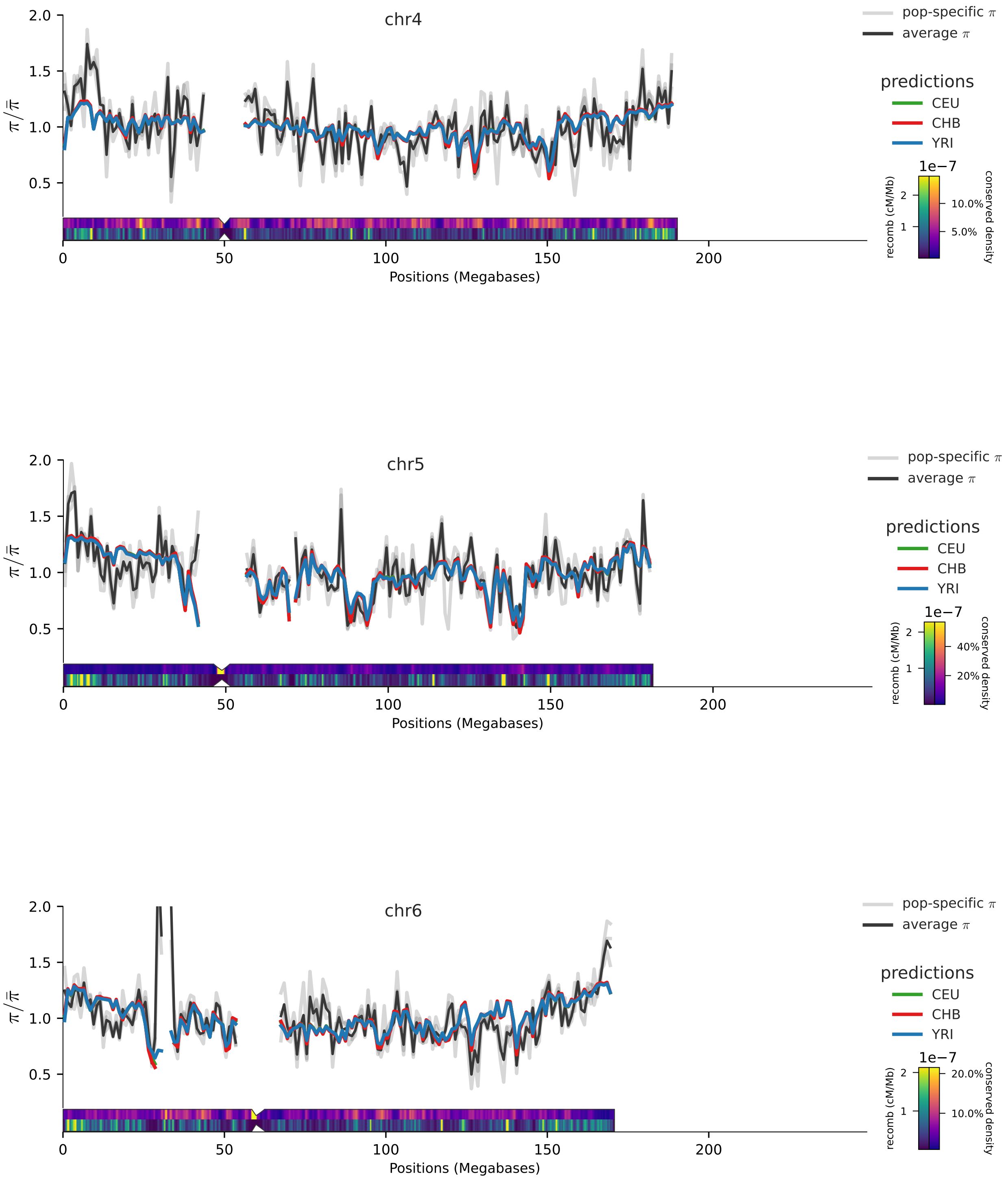

Figure 23: Note: we mask the MHC region during model fit.

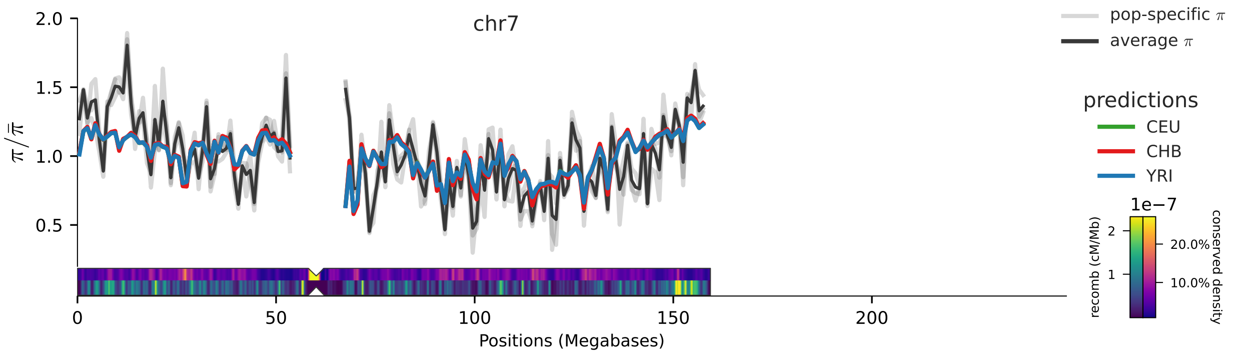
chr7
2.0
1.5
1.0
0.5
$\pi/\bar{\pi}$
0
50
100
150
200
Positions (Megabases)
pop-specific $\pi$
average $\pi$
predictions
CEU
CHB
YRI
1e−7
recomb (cM/Mb)
conserved density
20.0%
10.0%

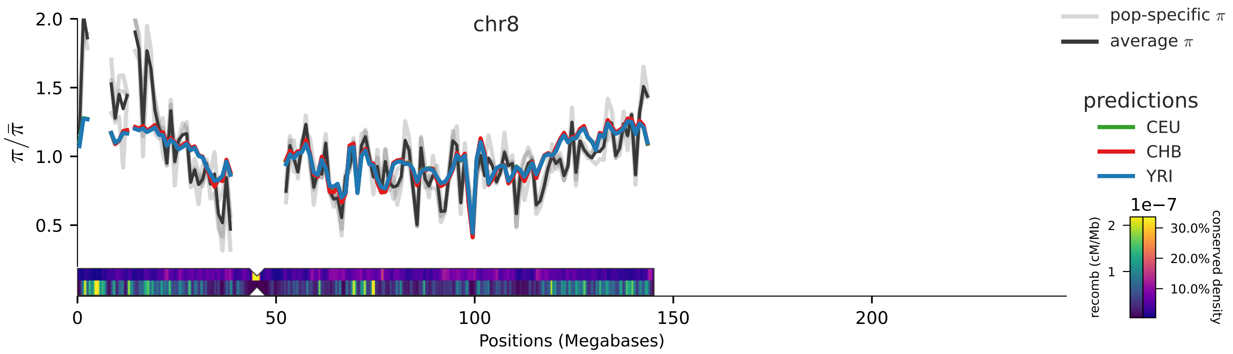
chr8
2.0
1.5
1.0
0.5
$\pi/\bar{\pi}$
0
50
100
150
200
Positions (Megabases)
pop-specific $\pi$
average $\pi$
predictions
CEU
CHB
YRI
1e−7
recomb (cM/Mb)
conserved density
30.0%
20.0%
10.0%

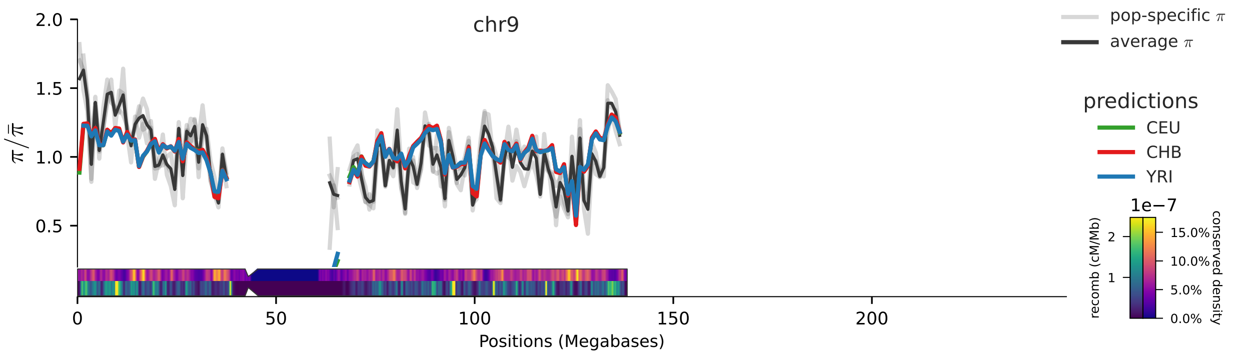
chr9
2.0
1.5
1.0
0.5
$\pi/\bar{\pi}$
0
50
100
150
200
Positions (Megabases)
pop-specific $\pi$
average $\pi$
predictions
CEU
CHB
YRI
1e−7
recomb (cM/Mb)
conserved density
15.0%
10.0%
5.0%
0.0%

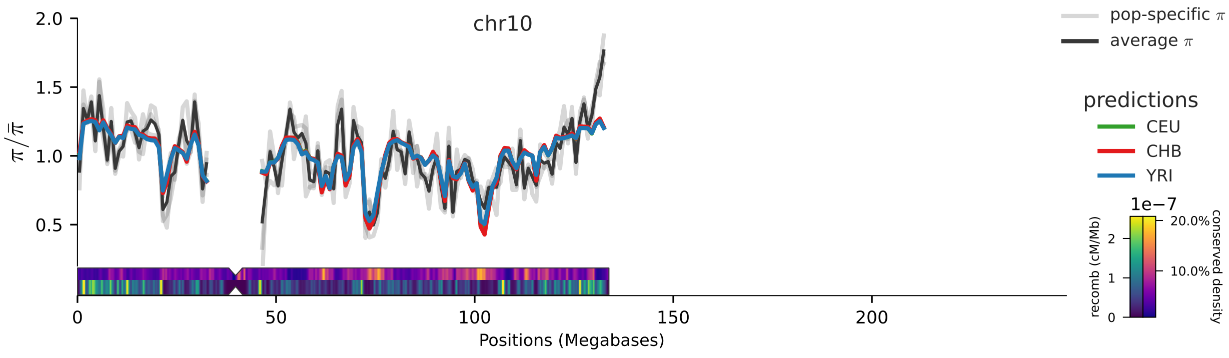
chr10
2.0
1.5
1.0
0.5
$\pi/\bar{\pi}$
0
50
100
150
200
Positions (Megabases)
pop-specific $\pi$
average $\pi$
predictions
CEU
CHB
YRI
1e−7
recomb (cM/Mb)
0
1
2
20.0%
10.0%
conserved density

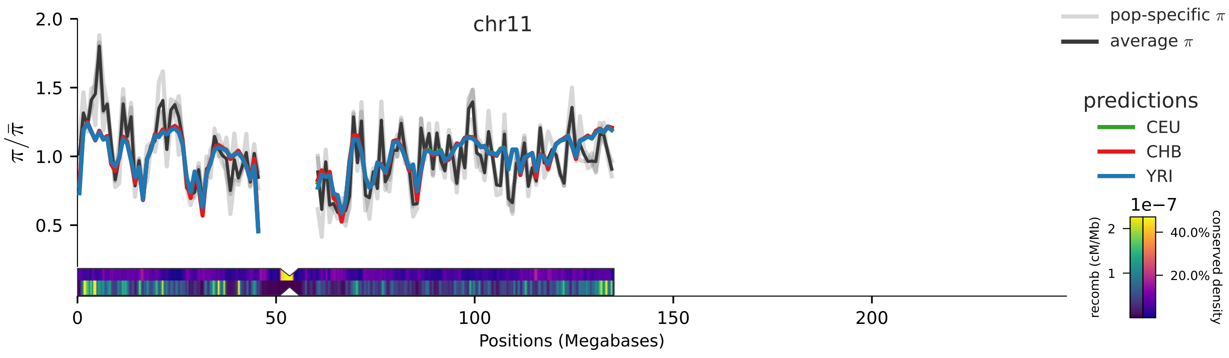
chr11
2.0
1.5
1.0
0.5
$\pi/\bar{\pi}$
0
50
100
150
200
Positions (Megabases)
pop-specific $\pi$
average $\pi$
predictions
CEU
CHB
YRI
1e−7
recomb (cM/Mb)
1
2
40.0%
20.0%
conserved density

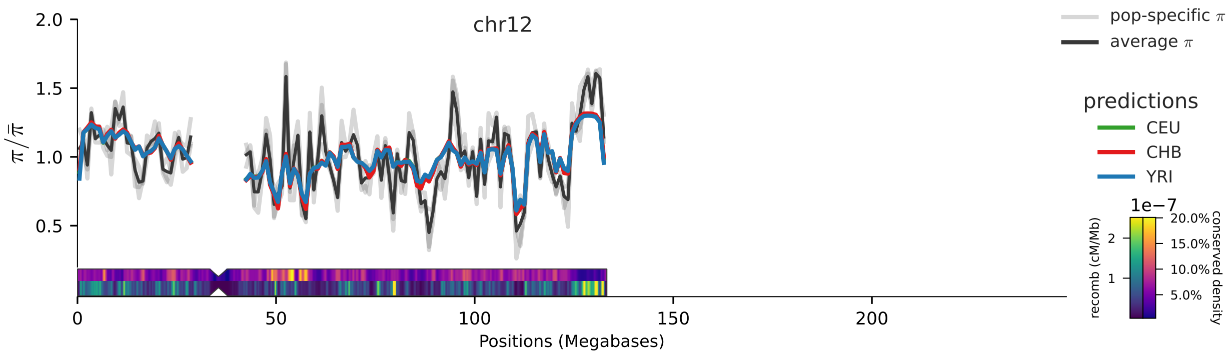
chr12
2.0
1.5
1.0
0.5
$\pi/\bar{\pi}$
0
50
100
150
200
Positions (Megabases)
pop-specific $\pi$
average $\pi$
predictions
CEU
CHB
YRI
1e−7
recomb (cM/Mb)
1
2
20.0%
15.0%
10.0%
5.0%
conserved density

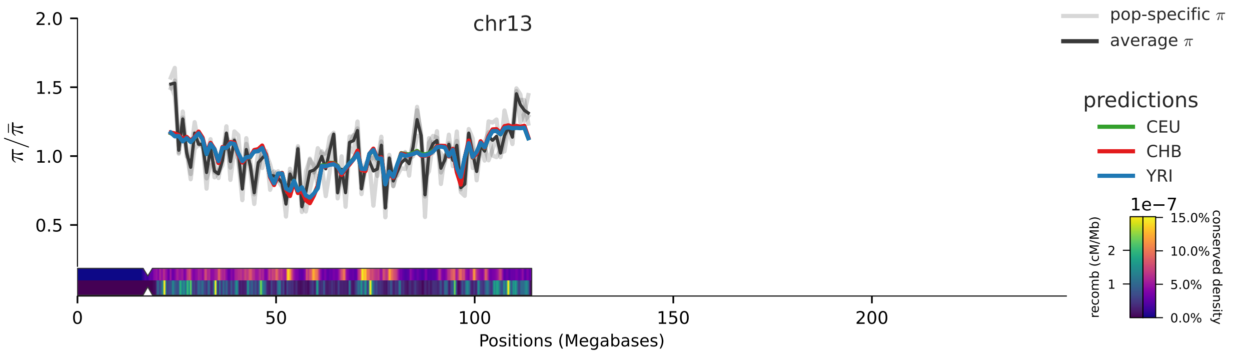
chr13
2.0
1.5
1.0
0.5
$\pi/\bar{\pi}$
0
50
100
150
200
Positions (Megabases)
pop-specific $\pi$
average $\pi$
predictions
CEU
CHB
YRI
1e−7
recomb (cM/Mb)
conserved density
15.0%
10.0%
5.0%
0.0%

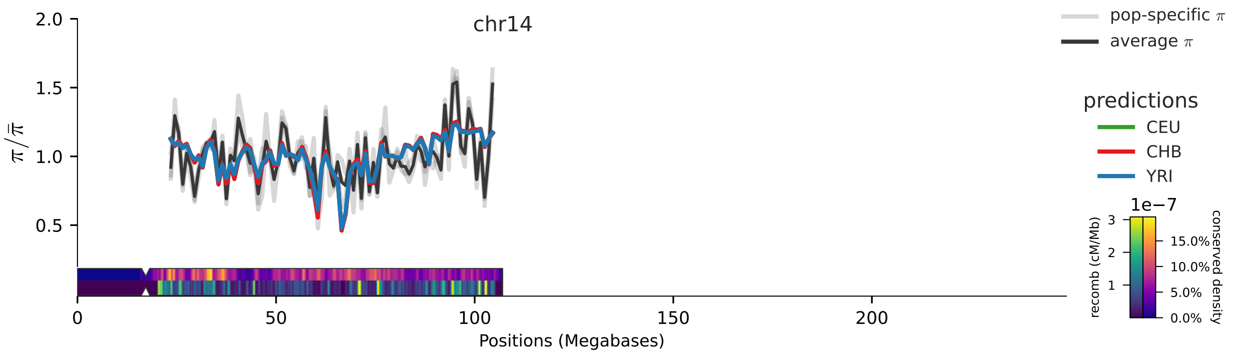
chr14
2.0
1.5
1.0
0.5
$\pi/\bar{\pi}$
0
50
100
150
200
Positions (Megabases)
pop-specific $\pi$
average $\pi$
predictions
CEU
CHB
YRI
1e−7
recomb (cM/Mb)
conserved density
15.0%
10.0%
5.0%
0.0%

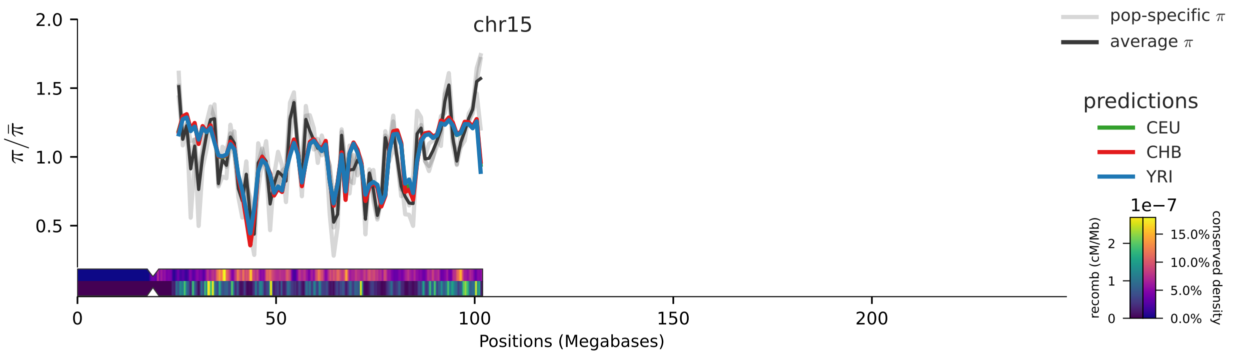
chr15
2.0
1.5
1.0
0.5
$\pi/\bar{\pi}$
0
50
100
150
200
Positions (Megabases)
pop-specific $\pi$
average $\pi$
predictions
CEU
CHB
YRI
1e−7
recomb (cM/Mb)
conserved density
15.0%
10.0%
5.0%
0.0%

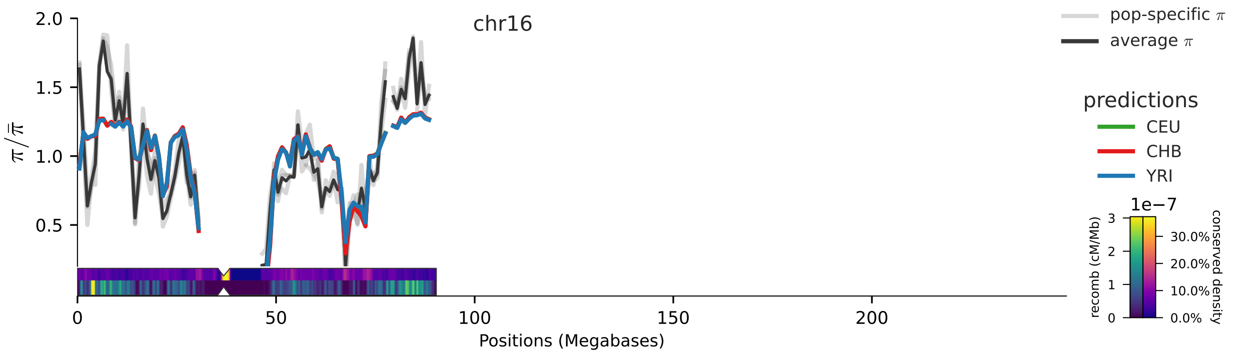
chr16
2.0
1.5
1.0
0.5
0
50
100
150
200
Positions (Megabases)
pop-specific π
average π
predictions
CEU
CHB
YRI
1e−7
recomb (cM/Mb)
3
2
1
0
30.0%
20.0%
10.0%
0.0%
conserved density

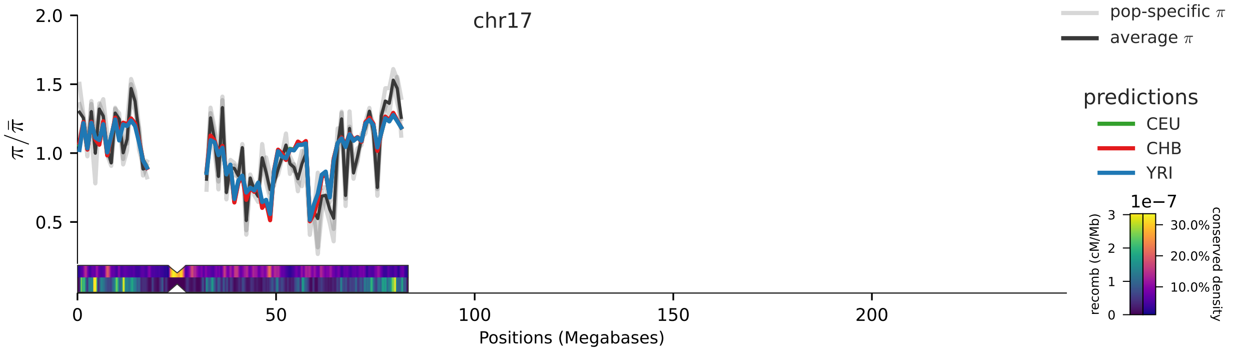
chr17
2.0
1.5
1.0
0.5
0
50
100
150
200
Positions (Megabases)
pop-specific π
average π
predictions
CEU
CHB
YRI
1e−7
recomb (cM/Mb)
3
2
1
0
30.0%
20.0%
10.0%
conserved density

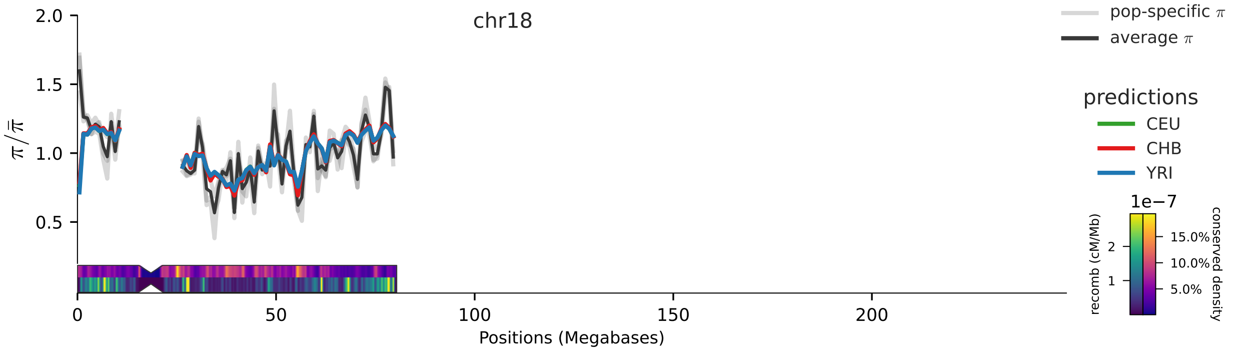
chr18
2.0
1.5
1.0
0.5
0
50
100
150
200
Positions (Megabases)
pop-specific π
average π
predictions
CEU
CHB
YRI
1e−7
recomb (cM/Mb)
2
1
15.0%
10.0%
5.0%
conserved density

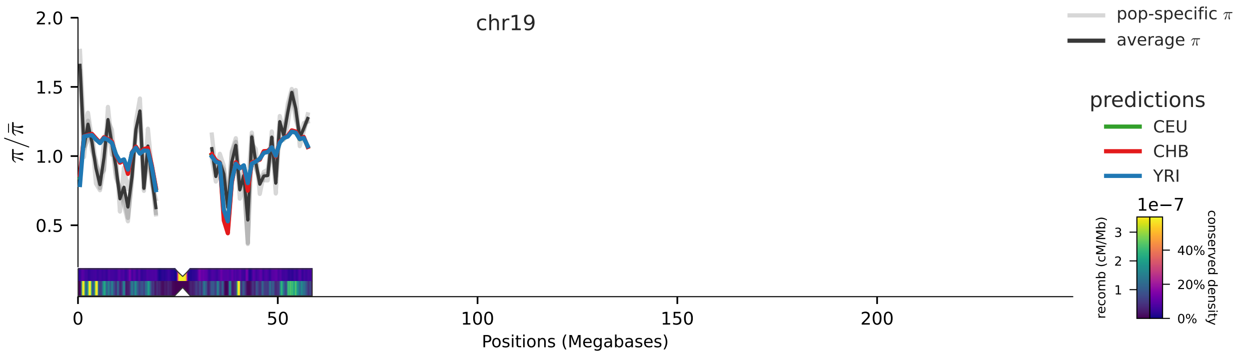
chr19
pop-specific $\pi$
average $\pi$
predictions
CEU
CHB
YRI
1e−7
recomb (cM/Mb)
conserved density
$\pi/\bar{\pi}$
Positions (Megabases)

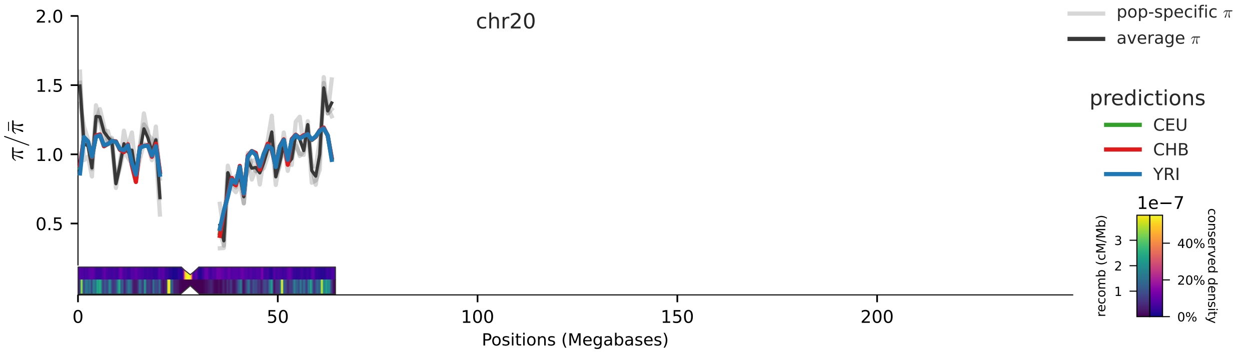
chr20
pop-specific $\pi$
average $\pi$
predictions
CEU
CHB
YRI
1e−7
recomb (cM/Mb)
conserved density
$\pi/\bar{\pi}$
Positions (Megabases)

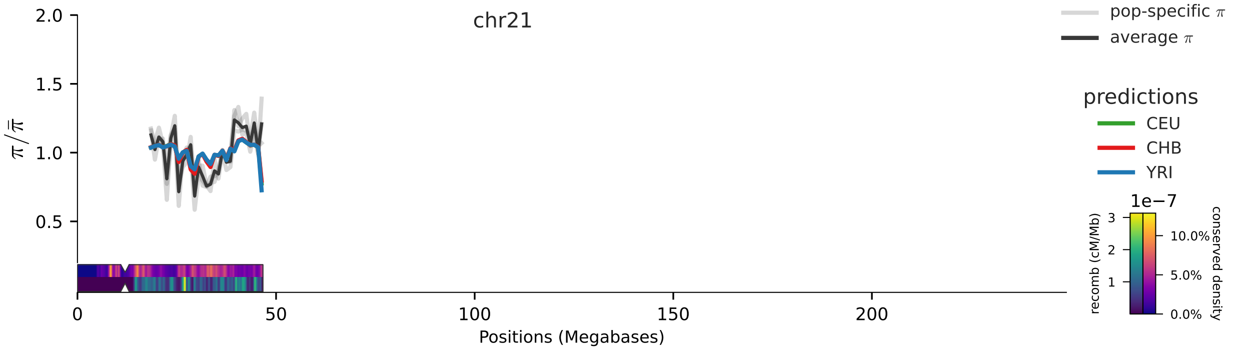
chr21
pop-specific $\pi$
average $\pi$
predictions
CEU
CHB
YRI
1e−7
recomb (cM/Mb)
conserved density
$\pi/\bar{\pi}$
Positions (Megabases)

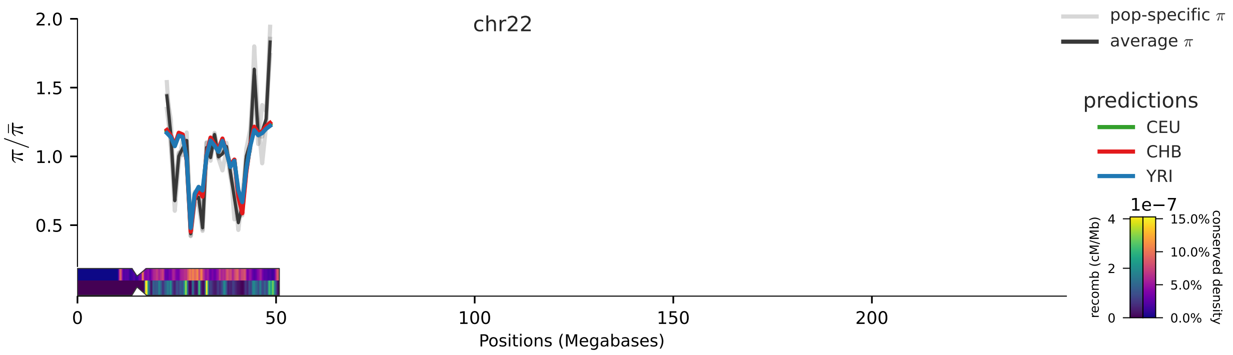
chr22
pop-specific $\pi$
average $\pi$
predictions
CEU
CHB
YRI
1e−7
recomb (cM/Mb)
conserved density
0
2
4
0.0%
5.0%
10.0%
15.0%
$\pi/\bar{\pi}$
2.0
1.5
1.0
0.5
0
50
100
150
200
Positions (Megabases)

### 6 All Model Fits

#### 6.1 Phastcons Priority (strong sel. grid), Full Track

##### CEU

ML estimates: (whole genome)
standard error method: moving block jackknife
negative log-likelihood: 197825873893.4953
number of successful starts: 10000 (100.0% total)
pi_0 = 0.0011279
pi  = 0.000801495
mut. rate = 2.35e-08
Ne = 28,197 (if mu=1e-8), Ne = 14,098 (if mu=2e-8)
R2 = 63.0106% (in-sample)  61.7759% (out-sample)
W =

| | CDS | gene | other | phastcons |
|---|---|---|---|---|
| 1e-08 | 0 | 0.901 | 0.984 | 0.034 |
| 1e-07 | 0 | 0.026 | 0 | 0.376 |
| 1e-06 | 0 | 0 | 0 | 0.001 |
| 1e-05 | 0 | 0.019 | 0 | 0 |
| 0.0001 | 0 | 0 | 0 | 0 |
| 0.001 | 0 | 0 | 0 | 0.472 |
| 0.01 | 0 | 0 | 0 | 0.046 |
| 0.1 | 1 | 0.053 | 0.016 | 0.071 |

#### CHB

ML estimates: (whole genome)

standard error method: moving block jackknife

negative log-likelihood: 61821783504.67356

number of successful starts: 10000 (100.0% total)

pi_0 = 0.00106467

pi  = 0.000752601

mut. rate = 2.557e-08

Ne = 26,616 (if mu=1e-8), Ne = 13,308 (if mu=2e-8)

R2 = 61.0847% (in-sample)  59.2923% (out-sample)

W =

| | CDS | gene | other | phastcons |
|---|---|---|---|---|
| 1e-08 | 0 | 0.931 | 0.985 | 0.478 |
| 1e-07 | 0 | 0.005 | 0 | 0.002 |
| 1e-06 | 0 | 0 | 0 | 0 |
| 1e-05 | 0 | 0.013 | 0 | 0 |
| 0.0001 | 0 | 0 | 0 | 0 |
| 0.001 | 0 | 0.003 | 0 | 0.43 |
| 0.01 | 0 | 0 | 0 | 0.09 |
| 0.1 | 1 | 0.049 | 0.015 | 0 |

#### YRI

ML estimates: (whole genome)

standard error method: moving block jackknife

negative log-likelihood: 250656989872.19766

number of successful starts: 10000 (100.0% total)

pi_0 = 0.00147277

pi  = 0.00106447

mut. rate = 1.708e-08

Ne = 36,819 (if mu=1e-8), Ne = 18,409 (if mu=2e-8)

R2 = 68.1177% (in-sample)  68.1947% (out-sample)

W =

| | CDS | gene | other | phastcons |
|---|---|---|---|---|
| 1e-08 | 1 | 0.902 | 0.979 | 0 |
| 1e-07 | 0 | 0 | 0 | 0.004 |
| 1e-06 | 0 | 0 | 0 | 0.001 |
| 1e-05 | 0 | 0.042 | 0 | 0 |
| 0.0001 | 0 | 0 | 0 | 0 |
| 0.001 | 0 | 0 | 0 | 0.523 |
| 0.01 | 0 | 0 | 0 | 0.212 |
| 0.1 | 0 | 0.056 | 0.021 | 0.26 |

#### 6.2 Phastcons Priority (strong sel. grid), Sparse Track

##### CEU

```
ML estimates: (whole genome)
standard error method: None
negative log-likelihood: 197826612684.87143
number of successful starts: 10000 (100.0% total)
pi_0 = 0.00112308
pi  = 0.000801495
mut. rate = 2.019e-08
Ne = 28,076 (if mu=1e-8), Ne = 14,038 (if mu=2e-8)
R2 = 63.0152% (in-sample)  61.9834% (out-sample)
W =
          CDS    gene    phastcons
------  -----  ------  -----------
1e-08   0       0.913        0.227
1e-07   0       0            0.004
1e-06   0.004   0            0
1e-05   0       0.019        0
0.0001  0       0            0
0.001   0       0            0.546
0.01    0       0            0.061
0.1     0.996   0.069        0.161
```

##### CHB

```
ML estimates: (whole genome)
standard error method: None
negative log-likelihood: 61822016128.79776
number of successful starts: 10000 (100.0% total)
pi_0 = 0.00106069
pi  = 0.000752601
mut. rate = 5.887e-08
Ne = 26,517 (if mu=1e-8), Ne = 13,258 (if mu=2e-8)
R2 = 61.0804% (in-sample)  59.4419% (out-sample)
W =
          CDS    gene    phastcons
------  -----  ------  -----------
1e-08   0.617   0.968        0.744
1e-07   0.01    0            0.02
1e-06   0       0            0
1e-05   0       0.005        0
0.0001  0       0            0
0.001   0       0.001        0.185
0.01    0       0            0.051
0.1     0.373   0.026        0
```

#### YRI

```
ML estimates: (whole genome)
standard error method: None
negative log-likelihood: 250657908564.79294
number of successful starts: 10000 (100.0% total)
pi_0 = 0.00146801
pi  = 0.00106447
mut. rate = 1.864e-08
Ne = 36,700 (if mu=1e-8), Ne = 18,350 (if mu=2e-8)
R2 = 68.1582% (in-sample)  68.4462% (out-sample)
W =
          CDS    gene    phastcons
------  -----  ------  -----------
1e-08       1   0.907        0
1e-07       0   0.001        0
1e-06       0   0            0
1e-05       0   0.035        0
0.0001      0   0            0
0.001       0   0            0.477
0.01        0   0            0.199
0.1         0   0.057        0.323
```

#### 6.3 Feature Priority (strong sel. grid), Full Track

##### CEU

```
ML estimates: (whole genome)
standard error method: moving block jackknife
negative log-likelihood: 197848627167.32043
number of successful starts: 10000 (100.0% total)
pi_0 = 0.00115698
pi  = 0.000801495
mut. rate = 4.1e-08
Ne = 28,924 (if mu=1e-8), Ne = 14,462 (if mu=2e-8)
R2 = 60.3026% (in-sample)  58.5808% (out-sample)
W =
          CDS    gene    other    phastcons
------  -----  ------  -------  -----------
1e-08   0.107   0.946        1        0
1e-07   0       0.001        0        0
1e-06   0       0            0        0
1e-05   0       0.023        0        0
0.0001  0       0            0        0
0.001   0.144   0.014        0        0.229
0.01    0       0            0        0.086
0.1     0.749   0.016        0        0.685
```

#### CHB

ML estimates: (whole genome)

standard error method: moving block jackknife

negative log-likelihood: 61827828856.16418

number of successful starts: 10000 (100.0% total)

pi_0 = 0.00108991

pi  = 0.000752601

mut. rate = 3.755e-08

Ne = 27,247 (if mu=1e-8), Ne = 13,623 (if mu=2e-8)

R2 = 58.7328% (in-sample)  56.4805% (out-sample)

W =

| | CDS | gene | other | phastcons |
|---|---|---|---|---|
| 1e-08 | 0.088 | 0.94 | 1 | 0.006 |
| 1e-07 | 0.01 | 0 | 0 | 0.003 |
| 1e-06 | 0 | 0 | 0 | 0 |
| 1e-05 | 0 | 0.023 | 0 | 0 |
| 0.0001 | 0 | 0 | 0 | 0 |
| 0.001 | 0.156 | 0.017 | 0 | 0.244 |
| 0.01 | 0 | 0 | 0 | 0.254 |
| 0.1 | 0.746 | 0.02 | 0 | 0.492 |

#### YRI

ML estimates: (whole genome)

standard error method: moving block jackknife

negative log-likelihood: 250679665159.79324

number of successful starts: 10000 (100.0% total)

pi_0 = 0.00151818

pi  = 0.00106447

mut. rate = 4.195e-08

Ne = 37,954 (if mu=1e-8), Ne = 18,977 (if mu=2e-8)

R2 = 65.5097% (in-sample)  64.9049% (out-sample)

W =

| | CDS | gene | other | phastcons |
|---|---|---|---|---|
| 1e-08 | 0.364 | 0.931 | 1 | 0.003 |
| 1e-07 | 0.001 | 0 | 0 | 0 |
| 1e-06 | 0 | 0 | 0 | 0 |
| 1e-05 | 0 | 0.039 | 0 | 0 |
| 0.0001 | 0 | 0 | 0 | 0 |
| 0.001 | 0.091 | 0.011 | 0 | 0.151 |
| 0.01 | 0 | 0 | 0 | 0.328 |
| 0.1 | 0.544 | 0.019 | 0 | 0.517 |

#### 6.4 Feature Priority (strong sel. grid), Sparse Track

##### CEU

```
ML estimates: (whole genome)
standard error method: moving block jackknife
negative log-likelihood: 197848626646.91736
number of successful starts: 10000 (100.0% total)
pi_0 = 0.00115698
pi  = 0.000801495
mut. rate = 4.104e-08
Ne = 28,924 (if mu=1e-8), Ne = 14,462 (if mu=2e-8)
R2 = 60.3027% (in-sample)  58.5823% (out-sample)
W =
          CDS    gene    phastcons
------  -----  ------  -----------
1e-08   0.107   0.947        0.001
1e-07   0       0            0
1e-06   0       0            0.001
1e-05   0       0.023        0
0.0001  0       0            0
0.001   0.144   0.014        0.229
0.01    0       0            0.086
0.1     0.748   0.016        0.684
```

##### CHB

```
ML estimates: (whole genome)
standard error method: moving block jackknife
negative log-likelihood: 61827828699.43107
number of successful starts: 10000 (100.0% total)
pi_0 = 0.00108991
pi  = 0.000752601
mut. rate = 3.719e-08
Ne = 27,247 (if mu=1e-8), Ne = 13,623 (if mu=2e-8)
R2 = 58.7329% (in-sample)  56.4805% (out-sample)
W =
          CDS    gene    phastcons
------  -----  ------  -----------
1e-08   0.089   0.938        0
1e-07   0       0.001        0
1e-06   0       0            0
1e-05   0       0.023        0
0.0001  0       0            0
0.001   0.157   0.017        0.246
0.01    0       0            0.257
0.1     0.754   0.02         0.497
```

#### YRI

```
ML estimates: (whole genome)
standard error method: moving block jackknife
negative log-likelihood: 250679664591.4161
number of successful starts: 10000 (100.0% total)
pi_0 = 0.00151817
pi  = 0.00106447
mut. rate = 4.179e-08
Ne = 37,954 (if mu=1e-8), Ne = 18,977 (if mu=2e-8)
R2 = 65.5099% (in-sample)  64.9049% (out-sample)
W =
          CDS    gene    phastcons
------  -----  ------  -----------
1e-08   0.36    0.931        0
1e-07   0.002   0            0
1e-06   0       0            0
1e-05   0       0.039        0
0.0001  0       0            0
0.001   0.092   0.011        0.151
0.01    0       0            0.329
0.1     0.546   0.019        0.519
```

#### 6.5 CADD 6% (strong selection DFE grid), Full Track

##### CEU

```
ML estimates: (whole genome)
standard error method: moving block jackknife
negative log-likelihood: 197826953189.02365
number of successful starts: 20000 (100.0% total)
pi_0 = 0.00109999
pi  = 0.000801495
mut. rate = 1.537e-08
Ne = 27,499 (if mu=1e-8), Ne = 13,749 (if mu=2e-8)
R2 = 62.9394% (in-sample)  61.7433% (out-sample)
W =
          cadd6    other
------  -------  -------
1e-08     0        0.965
1e-07     0        0
1e-06     0        0
1e-05     0        0
0.0001    0.026    0
0.001     0.555    0
0.01      0        0
0.1       0.419    0.035
```

#### CHB

```
ML estimates: (whole genome)
standard error method: moving block jackknife
negative log-likelihood: 61821323176.26884
number of successful starts: 20000 (100.0% total)
pi_0 = 0.00103624
pi  = 0.000752601
mut. rate = 1.508e-08
Ne = 25,905 (if mu=1e-8), Ne = 12,952 (if mu=2e-8)
R2 = 61.3115% (in-sample)  59.6228% (out-sample)
W =
          cadd6    other
------  -------  -------
1e-08     0        0.961
1e-07     0.007    0
1e-06     0.009    0
1e-05     0        0
0.0001    0.004    0
0.001     0.67     0
0.01      0        0
0.1       0.309    0.039
```

#### YRI

```
ML estimates: (whole genome)
standard error method: moving block jackknife
negative log-likelihood: 250659830283.90668
number of successful starts: 20000 (100.0% total)
pi_0 = 0.00145893
pi  = 0.00106447
mut. rate = 1.461e-08
Ne = 36,473 (if mu=1e-8), Ne = 18,236 (if mu=2e-8)
R2 = 68.0839% (in-sample)  68.0751% (out-sample)
W =
          cadd6    other
------  -------  -------
1e-08     0        0.958
1e-07     0.023    0
1e-06     0.001    0
1e-05     0        0
0.0001    0.065    0
0.001     0.407    0
0.01      0.078    0
0.1       0.425    0.041
```

#### 6.6 CADD 8% (strong selection DFE grid), Full Track

##### CEU

```
ML estimates: (whole genome)
standard error method: moving block jackknife
negative log-likelihood: 197834651568.3971
number of successful starts: 10000 (100.0% total)
pi_0 = 0.00109606
pi  = 0.000801495
mut. rate = 1.157e-08
Ne = 27,401 (if mu=1e-8), Ne = 13,700 (if mu=2e-8)
R2 = 62.1245% (in-sample)  60.8009% (out-sample)
W =
          cadd8    other
------  -------  -------
1e-08     0        0.952
1e-07     0.004    0
1e-06     0        0
1e-05     0        0
0.0001    0.052    0
0.001     0.537    0
0.01      0        0
0.1       0.407    0.048
```

##### CHB

```
ML estimates: (whole genome)
standard error method: moving block jackknife
negative log-likelihood: 61823378586.32404
number of successful starts: 10000 (100.0% total)
pi_0 = 0.00103245
pi  = 0.000752601
mut. rate = 1.113e-08
Ne = 25,811 (if mu=1e-8), Ne = 12,905 (if mu=2e-8)
R2 = 60.6265% (in-sample)  58.8768% (out-sample)
W =
          cadd8    other
------  -------  -------
1e-08     0        0.944
1e-07     0        0.001
1e-06     0        0
1e-05     0        0
0.0001    0.026    0
0.001     0.69     0
0.01      0        0
0.1       0.284    0.055
```

#### YRI

ML estimates: (whole genome)
standard error method: moving block jackknife
negative log-likelihood: 250669695762.61414
number of successful starts: 10000 (100.0% total)
pi_0 = 0.00145867
pi  = 0.00106447
mut. rate = 1.013e-08
Ne = 36,466 (if mu=1e-8), Ne = 18,233 (if mu=2e-8)
R2 = 67.1076% (in-sample)  66.9835% (out-sample)
W =

| | cadd8 | other |
|---|---|---|
| 1e-08 | 0 | 0.932 |
| 1e-07 | 0.002 | 0 |
| 1e-06 | 0 | 0 |
| 1e-05 | 0 | 0 |
| 0.0001 | 0.103 | 0 |
| 0.001 | 0.438 | 0 |
| 0.01 | 0 | 0 |
| 0.1 | 0.458 | 0.068 |

#### 6.7 Phastcons Priority, Full Track

##### CEU

ML estimates: (whole genome)
number jackknife samples: 264
standard error method: moving block jackknife
negative log-likelihood: 197835157725.57724
number of successful starts: 10000 (100.0% total)
pi_0 = 0.00105552 (0.00106, 0.00106)
pi  = 0.000801495
mut. rate = 1.912e-08  (1.32e-08, 2.5e-08)
Ne = 26,387 (if mu=1e-8), Ne = 13,193 (if mu=2e-8)
R2 = 62.2547% (in-sample)  61.0392% (out-sample)
W =

| | CDS | gene | other | phastcons |
|---|---|---|---|---|
| 1e-08 | 1.000 (0.9744, 1.0256) | 0.947 (0.8398, 1.0542) | 1.000 (0.9959, 1.0041) | 0.000 (-0.2261, 0.2262) |
| 1e-07 | 0.000 (-0.0250, 0.0250) | 0.000 (-0.1053, 0.1053) | 0.000 (-0.0041, 0.0041) | 0.000 (-0.1200, 0.1200) |
| 1e-06 | 0.000 (-0.0021, 0.0021) | 0.000 (-0.0010, 0.0010) | 0.000 (-0.0001, 0.0001) | 0.000 (-0.0280, 0.0280) |
| 1e-05 | 0.000 (-0.0000, 0.0000) | 0.030 (0.0209, 0.0400) | 0.000 (-0.0000, 0.0000) | 0.000 (-0.0004, 0.0004) |
| 0.0001 | 0.000 (-0.0000, 0.0000) | 0.000 (-0.0000, 0.0000) | 0.000 (-0.0000, 0.0000) | 0.000 (-0.0000, 0.0001) |
| 0.001 | 0.000 (-0.0000, 0.0000) | 0.000 (-0.0000, 0.0000) | 0.000 (-0.0000, 0.0000) | 0.413 (0.2960, 0.5308) |
| 0.01 | 0.000 (-0.0001, 0.0001) | 0.023 (0.0160, 0.0292) | 0.000 (-0.0000, 0.0000) | 0.586 (0.4205, 0.7525) |

#### CHB

```
ML estimates: (whole genome)
number jackknife samples: 266
standard error method: moving block jackknife
negative log-likelihood: 61824235931.79921
number of successful starts: 10000 (100.0% total)
pi_0 = 0.00100129 (0.00100, 0.00100)
pi  = 0.000752601
mut. rate = 1.869e-08  (1.16e-08, 2.57e-08)
Ne = 25,032 (if mu=1e-8), Ne = 12,516 (if mu=2e-8)
R2 = 60.4486% (in-sample)  58.7636% (out-sample)
W =
        CDS                      gene                     other                    phastcons
------  -----------------------  -----------------------  -----------------------  -----------------------
1e-08   1.000 (0.8992, 1.1008)   0.940 (0.7946, 1.0845)   1.000 (0.9996, 1.0004)   0.000 (-0.2201, 0.2202)
1e-07   0.000 (-0.1006, 0.1006)  0.000 (-0.1420, 0.1420)  0.000 (-0.0003, 0.0003)  0.000 (-0.2016, 0.2017)
1e-06   0.000 (-0.0021, 0.0021)  0.000 (-0.0025, 0.0025)  0.000 (-0.0001, 0.0001)  0.000 (-0.0118, 0.0119)
1e-05   0.000 (-0.0000, 0.0000)  0.031 (0.0161, 0.0457)   0.000 (-0.0000, 0.0000)  0.000 (-0.0002, 0.0002)
0.0001  0.000 (-0.0000, 0.0000)  0.000 (-0.0000, 0.0000)  0.000 (-0.0000, 0.0000)  0.000 (-0.0000, 0.0000)
0.001   0.000 (-0.0000, 0.0000)  0.000 (-0.0000, 0.0000)  0.000 (-0.0000, 0.0000)  0.457 (0.2998, 0.6141)
0.01    0.000 (-0.0000, 0.0000)  0.030 (0.0193, 0.0399)   0.000 (-0.0000, 0.0000)  0.543 (0.3558, 0.7302)
```

#### YRI

```
ML estimates: (whole genome)
number jackknife samples: 264
standard error method: moving block jackknife
negative log-likelihood: 250667315874.309
number of successful starts: 10000 (100.0% total)
pi_0 = 0.00139482 (0.00139, 0.00140)
pi  = 0.00106447
mut. rate = 2.004e-08  (1.53e-08, 2.48e-08)
Ne = 34,870 (if mu=1e-8), Ne = 17,435 (if mu=2e-8)
R2 = 67.3564% (in-sample)  67.274% (out-sample)
W =
        CDS                      gene                     other                    phastcons
------  -----------------------  -----------------------  -----------------------  -----------------------
1e-08   1.000 (0.9646, 1.0354)   0.955 (0.8905, 1.0192)   1.000 (0.9972, 1.0028)   0.000 (-0.1515, 0.1516)
1e-07   0.000 (-0.0352, 0.0352)  0.000 (-0.0636, 0.0636)  0.000 (-0.0028, 0.0028)  0.000 (-0.1435, 0.1435)
1e-06   0.000 (-0.0008, 0.0008)  0.000 (-0.0009, 0.0011)  0.000 (-0.0000, 0.0000)  0.000 (-0.0101, 0.0101)
1e-05   0.000 (-0.0000, 0.0000)  0.029 (0.0192, 0.0384)   0.000 (-0.0000, 0.0000)  0.000 (-0.0002, 0.0002)
0.0001  0.000 (-0.0000, 0.0000)  0.000 (-0.0000, 0.0000)  0.000 (-0.0000, 0.0000)  0.027 (0.0108, 0.0435)
0.001   0.000 (-0.0000, 0.0000)  0.000 (-0.0000, 0.0000)  0.000 (-0.0000, 0.0000)  0.227 (0.1661, 0.2882)
0.01    0.000 (-0.0000, 0.0000)  0.016 (0.0124, 0.0201)   0.000 (-0.0000, 0.0000)  0.746 (0.5776, 0.9138)
```

#### 6.8 Phastcons Priority, Sparse Track

##### CEU

```
ML estimates: (whole genome)
standard error method: None
negative log-likelihood: 197835157553.76733
number of successful starts: 10000 (100.0% total)
pi_0 = 0.00105552
pi  = 0.000801495
mut. rate = 2.313e-08
Ne = 26,387 (if mu=1e-8), Ne = 13,193 (if mu=2e-8)
R2 = 62.2548% (in-sample)  61.0392% (out-sample)
W =
          CDS    gene    phastcons
------  -----  ------  -----------
1e-08       1   0.954        0.174
1e-07       0   0.002        0
1e-06       0   0            0
1e-05       0   0.025        0
0.0001      0   0            0
0.001       0   0            0.342
0.01        0   0.019        0.485
```

##### CHB

```
ML estimates: (whole genome)
standard error method: None
negative log-likelihood: 61824235897.32505
number of successful starts: 10000 (100.0% total)
pi_0 = 0.00100129
pi  = 0.000752601
mut. rate = 1.92e-08
Ne = 25,032 (if mu=1e-8), Ne = 12,516 (if mu=2e-8)
R2 = 60.4487% (in-sample)  58.8423% (out-sample)
W =
          CDS    gene    phastcons
------  -----  ------  -----------
1e-08       1   0.939        0.003
1e-07       0   0.002        0.022
1e-06       0   0            0.002
1e-05       0   0.03         0
0.0001      0   0            0
0.001       0   0            0.445
0.01        0   0.029        0.528
```

#### YRI

ML estimates: (whole genome)
standard error method: None
negative log-likelihood: 250667315734.13507
number of successful starts: 10000 (100.0% total)
pi_0 = 0.00139482
pi = 0.00106447
mut. rate = 2.004e-08
Ne = 34,870 (if mu=1e-8), Ne = 17,435 (if mu=2e-8)
R2 = 67.3565% (in-sample) 67.2897% (out-sample)
W =

| | CDS | gene | phastcons |
|---|---|---|---|
| 1e-08 | 1 | 0.954 | 0 |
| 1e-07 | 0 | 0.001 | 0 |
| 1e-06 | 0 | 0 | 0 |
| 1e-05 | 0 | 0.029 | 0 |
| 0.0001 | 0 | 0 | 0.027 |
| 0.001 | 0 | 0 | 0.227 |
| 0.01 | 0 | 0.016 | 0.746 |

#### 6.9 Feature Priority, Full Track

##### CEU

ML estimates: (whole genome)
number jackknife samples: 116
standard error method: moving block jackknife
negative log-likelihood: 197860660948.8551
number of successful starts: 10000 (100.0% total)
pi_0 = 0.00109138 (0.00109, 0.00109)
pi = 0.000801495
mut. rate = 3.044e-08 (2.79e-08, 3.3e-08)
Ne = 27,284 (if mu=1e-8), Ne = 13,642 (if mu=2e-8)
R2 = 59.1052% (in-sample) 57.1447% (out-sample)
W =

| | CDS | gene | other | phastcons |
|---|---|---|---|---|
| 1e-08 | 0.412 (0.2565, 0.5672) | 0.884 (0.8329, 0.9356) | 1.000 (0.9996, 1.0003) | 0.000 (-0.0666, 0.0666) |
| 1e-07 | 0.000 (-0.1509, 0.1511) | 0.003 (-0.0480, 0.0534) | 0.000 (-0.0003, 0.0004) | 0.000 (-0.0248, 0.0248) |
| 1e-06 | 0.000 (-0.0021, 0.0025) | 0.000 (-0.0005, 0.0005) | 0.000 (-0.0000, 0.0000) | 0.000 (-0.0054, 0.0054) |
| 1e-05 | 0.000 (-0.0000, 0.0000) | 0.074 (0.0608, 0.0875) | 0.000 (-0.0000, 0.0000) | 0.000 (-0.0001, 0.0001) |
| 0.0001 | 0.000 (-0.0000, 0.0000) | 0.000 (-0.0000, 0.0000) | 0.000 (-0.0000, 0.0000) | 0.000 (-0.0000, 0.0000) |
| 0.001 | 0.066 (0.0340, 0.0982) | 0.008 (0.0064, 0.0102) | 0.000 (-0.0000, 0.0000) | 0.136 (0.1225, 0.1503) |
| 0.01 | 0.522 (0.4497, 0.5939) | 0.031 (0.0276, 0.0336) | 0.000 (-0.0000, 0.0000) | 0.864 (0.7932, 0.9340) |

#### CHB

ML estimates: (whole genome)
number jackknife samples: 256
standard error method: moving block jackknife
negative log-likelihood: 61830715796.965416
number of successful starts: 10000 (100.0% total)
pi_0 = 0.00103388 (0.00103, 0.00104)
pi  = 0.000752601
mut. rate = 3.172e-08  (2.73e-08, 3.62e-08)
Ne = 25,847 (if mu=1e-8), Ne = 12,923 (if mu=2e-8)
R2 = 57.8792% (in-sample)  55.631% (out-sample)
W =

| | CDS | gene | other | phastcons |
|---|---|---|---|---|
| 1e-08 | 0.430 (0.1137, 0.7471) | 0.894 (0.8175, 0.9702) | 1.000 (0.9993, 1.0007) | 0.000 (-0.1107, 0.1107) |
| 1e-07 | 0.001 (-0.2978, 0.3001) | 0.004 (-0.0686, 0.0772) | 0.000 (-0.0007, 0.0007) | 0.000 (-0.0393, 0.0393) |
| 1e-06 | 0.000 (-0.0035, 0.0042) | 0.000 (-0.0006, 0.0006) | 0.000 (-0.0000, 0.0000) | 0.000 (-0.0053, 0.0053) |
| 1e-05 | 0.000 (-0.0000, 0.0000) | 0.064 (0.0437, 0.0848) | 0.000 (-0.0000, 0.0000) | 0.000 (-0.0001, 0.0001) |
| 0.0001 | 0.000 (-0.0000, 0.0000) | 0.000 (-0.0000, 0.0000) | 0.000 (-0.0000, 0.0000) | 0.000 (-0.0000, 0.0000) |
| 0.001 | 0.052 (0.0045, 0.1000) | 0.012 (0.0086, 0.0148) | 0.000 (-0.0000, 0.0000) | 0.166 (0.1391, 0.1926) |
| 0.01 | 0.516 (0.4059, 0.6258) | 0.026 (0.0216, 0.0302) | 0.000 (-0.0000, 0.0000) | 0.834 (0.7222, 0.9461) |

#### YRI

ML estimates: (whole genome)
number jackknife samples: 241
standard error method: moving block jackknife
negative log-likelihood: 250692005490.22156
number of successful starts: 10000 (100.0% total)
pi_0 = 0.00144148 (0.00144, 0.00144)
pi  = 0.00106447
mut. rate = 3.349e-08  (2.85e-08, 3.85e-08)
Ne = 36,037 (if mu=1e-8), Ne = 18,018 (if mu=2e-8)
R2 = 64.2844% (in-sample)  63.058% (out-sample)
W =

| | CDS | gene | other | phastcons |
|---|---|---|---|---|
| 1e-08 | 0.752 (0.4939, 1.0110) | 0.878 (0.8209, 0.9353) | 1.000 (0.9998, 1.0002) | 0.000 (-0.1066, 0.1066) |
| 1e-07 | 0.000 (-0.2548, 0.2552) | 0.000 (-0.0533, 0.0533) | 0.000 (-0.0002, 0.0002) | 0.000 (-0.0669, 0.0669) |
| 1e-06 | 0.000 (-0.0028, 0.0029) | 0.000 (-0.0005, 0.0005) | 0.000 (-0.0000, 0.0000) | 0.000 (-0.0087, 0.0087) |
| 1e-05 | 0.000 (-0.0000, 0.0000) | 0.085 (0.0721, 0.0986) | 0.000 (-0.0000, 0.0000) | 0.000 (-0.0002, 0.0002) |
| 0.0001 | 0.000 (-0.0000, 0.0000) | 0.000 (-0.0000, 0.0000) | 0.000 (-0.0000, 0.0000) | 0.000 (-0.0000, 0.0000) |
| 0.001 | 0.100 (0.0858, 0.1136) | 0.003 (0.0026, 0.0040) | 0.000 (-0.0000, 0.0000) | 0.070 (0.0577, 0.0823) |
| 0.01 | 0.148 (0.1266, 0.1688) | 0.033 (0.0285, 0.0379) | 0.000 (-0.0000, 0.0000) | 0.930 (0.8006, 1.0594) |

#### 6.10 Feature Priority, Sparse Track

##### CEU

```
ML estimates: (whole genome)
standard error method: None
negative log-likelihood: 197860660635.007
number of successful starts: 10000 (100.0% total)
pi_0 = 0.00109138
pi  = 0.000801495
mut. rate = 3.044e-08
Ne = 27,284 (if mu=1e-8), Ne = 13,642 (if mu=2e-8)
R2 = 59.1053% (in-sample)  57.2393% (out-sample)
W =
```

| | CDS | gene | phastcons |
|---|---|---|---|
| 1e-08 | 0.412 | 0.887 | 0 |
| 1e-07 | 0 | 0 | 0 |
| 1e-06 | 0.001 | 0 | 0 |
| 1e-05 | 0 | 0.074 | 0 |
| 0.0001 | 0 | 0 | 0 |
| 0.001 | 0.066 | 0.008 | 0.136 |
| 0.01 | 0.522 | 0.031 | 0.864 |

##### CHB

```
ML estimates: (whole genome)
standard error method: None
negative log-likelihood: 61830715695.97452
number of successful starts: 10000 (100.0% total)
pi_0 = 0.00103388
pi  = 0.000752601
mut. rate = 3.172e-08
Ne = 25,847 (if mu=1e-8), Ne = 12,923 (if mu=2e-8)
R2 = 57.8793% (in-sample)  55.9962% (out-sample)
W =
```

| | CDS | gene | phastcons |
|---|---|---|---|
| 1e-08 | 0.418 | 0.898 | 0 |
| 1e-07 | 0.014 | 0 | 0 |
| 1e-06 | 0 | 0 | 0 |
| 1e-05 | 0 | 0.064 | 0 |
| 0.0001 | 0 | 0 | 0 |
| 0.001 | 0.052 | 0.012 | 0.166 |
| 0.01 | 0.516 | 0.026 | 0.834 |

#### YRI

```
ML estimates: (whole genome)
standard error method: None
negative log-likelihood: 250692005023.91275
number of successful starts: 10000 (100.0% total)
pi_0 = 0.00144148
pi  = 0.00106447
mut. rate = 3.353e-08
Ne = 36,037 (if mu=1e-8), Ne = 18,018 (if mu=2e-8)
R2 = 64.2844% (in-sample)  63.6078% (out-sample)
W =
          CDS    gene    phastcons
------  -----  ------  -----------
1e-08   0.747   0.875        0
1e-07   0.006   0.004        0.001
1e-06   0       0            0
1e-05   0       0.085        0
0.0001  0       0            0
0.001   0.1     0.003        0.07
0.01    0.148   0.033        0.929
```

#### 6.11 CADD 6%, Full Track

##### CEU

```
ML estimates: (whole genome)
number jackknife samples: 280
standard error method: moving block jackknife
negative log-likelihood: 197839911915.8117
number of successful starts: 10000 (100.0% total)
pi_0 = 0.00102951 (0.00103, 0.00103)
pi  = 0.000801495
mut. rate = 1.635e-08  (1.2e-08, 2.07e-08)
Ne = 25,737 (if mu=1e-8), Ne = 12,868 (if mu=2e-8)
R2 = 61.767% (in-sample)  60.2433% (out-sample)
W =
        cadd6                    other
------  -----------------------  -----------------------
1e-08   0.002 (-0.1758, 0.1800)  0.997 (0.9918, 1.0019)
1e-07   0.006 (-0.1224, 0.1353)  0.000 (-0.0049, 0.0049)
1e-06   0.000 (-0.0079, 0.0084)  0.000 (-0.0001, 0.0001)
1e-05   0.000 (-0.0002, 0.0002)  0.000 (-0.0000, 0.0000)
0.0001  0.049 (0.0332, 0.0649)   0.000 (-0.0000, 0.0000)
0.001   0.325 (0.2429, 0.4072)   0.000 (-0.0000, 0.0000)
0.01    0.617 (0.4630, 0.7711)   0.003 (0.0022, 0.0042)
```

#### CHB

```
ML estimates: (whole genome)
number jackknife samples: 279
standard error method: moving block jackknife
negative log-likelihood: 61824516495.02472
number of successful starts: 10000 (100.0% total)
pi_0 = 0.000977579 (0.00098, 0.00098)
pi  = 0.000752601
mut. rate = 1.598e-08  (1.15e-08, 2.05e-08)
Ne = 24,439 (if mu=1e-8), Ne = 12,219 (if mu=2e-8)
R2 = 60.4032% (in-sample)  58.4847% (out-sample)
W =
        cadd6                    other
------  -----------------------  -----------------------
1e-08   0.000 (-0.1922, 0.1922)  0.992 (0.9829, 1.0002)
1e-07   0.001 (-0.1431, 0.1441)  0.001 (-0.0076, 0.0088)
1e-06   0.000 (-0.0068, 0.0068)  0.000 (-0.0001, 0.0001)
1e-05   0.000 (-0.0002, 0.0002)  0.000 (-0.0000, 0.0000)
0.0001  0.030 (0.0129, 0.0476)   0.000 (-0.0000, 0.0000)
0.001   0.440 (0.3238, 0.5568)   0.000 (-0.0000, 0.0000)
0.01    0.529 (0.3902, 0.6676)   0.008 (0.0056, 0.0101)
```

#### YRI

```
ML estimates: (whole genome)
number jackknife samples: 280
standard error method: moving block jackknife
negative log-likelihood: 250675195472.4508
number of successful starts: 10000 (100.0% total)
pi_0 = 0.00136706 (0.00137, 0.00137)
pi  = 0.00106447
mut. rate = 1.556e-08  (1.25e-08, 1.86e-08)
Ne = 34,176 (if mu=1e-8), Ne = 17,088 (if mu=2e-8)
R2 = 66.6755% (in-sample)  66.5047% (out-sample)
W =
        cadd6                    other
------  -----------------------  -----------------------
1e-08   0.000 (-0.1074, 0.1079)  0.994 (0.9892, 0.9985)
1e-07   0.006 (-0.1023, 0.1133)  0.000 (-0.0045, 0.0045)
1e-06   0.000 (-0.0038, 0.0038)  0.000 (-0.0001, 0.0001)
1e-05   0.000 (-0.0001, 0.0001)  0.000 (-0.0000, 0.0000)
0.0001  0.099 (0.0783, 0.1204)   0.000 (-0.0000, 0.0000)
0.001   0.115 (0.0848, 0.1448)   0.000 (-0.0000, 0.0000)
0.01    0.780 (0.6381, 0.9221)   0.006 (0.0049, 0.0074)
```

#### 6.12 CADD 6%, Sparse Track

##### CEU

```
ML estimates: (whole genome)
standard error method: None
negative log-likelihood: 197839997860.2112
number of successful starts: 10000 (100.0% total)
pi_0 = 0.00102636
pi  = 0.000801495
mut. rate = 1.68e-08
Ne = 25,659 (if mu=1e-8), Ne = 12,829 (if mu=2e-8)
R2 = 61.8005% (in-sample)  60.3663% (out-sample)
W =
          cadd6
------  -------
1e-08     0
1e-07     0
1e-06     0
1e-05     0
0.0001    0.046
0.001     0.317
0.01      0.638
```

##### CHB

```
ML estimates: (whole genome)
standard error method: None
negative log-likelihood: 61824669694.580444
number of successful starts: 10000 (100.0% total)
pi_0 = 0.000970441
pi  = 0.000752601
mut. rate = 1.745e-08
Ne = 24,261 (if mu=1e-8), Ne = 12,130 (if mu=2e-8)
R2 = 60.4898% (in-sample)  58.67% (out-sample)
W =
          cadd6
------  -------
1e-08     0.003
1e-07     0
1e-06     0
1e-05     0
0.0001    0.024
0.001     0.403
0.01      0.57
```

#### YRI

```
ML estimates: (whole genome)
standard error method: None
negative log-likelihood: 250675581171.07547
number of successful starts: 10000 (100.0% total)
pi_0 = 0.00135936
pi  = 0.00106447
mut. rate = 1.657e-08
Ne = 33,983 (if mu=1e-8), Ne = 16,991 (if mu=2e-8)
R2 = 66.8084% (in-sample)  66.6575% (out-sample)
W =
          cadd6
------  -------
1e-08     0
1e-07     0
1e-06     0
1e-05     0
0.0001    0.09
0.001     0.109
0.01      0.801
```

### 6.13 CADD 8%, Full Track

#### CEU

```
ML estimates: (whole genome)
number jackknife samples: 241
standard error method: moving block jackknife
negative log-likelihood: 197849369068.89285
number of successful starts: 10000 (100.0% total)
pi_0 = 0.00102211 (0.00102, 0.00102)
pi  = 0.000801495
mut. rate = 1.208e-08  (8.74e-09, 1.54e-08)
Ne = 25,552 (if mu=1e-8), Ne = 12,776 (if mu=2e-8)
R2 = 60.8766% (in-sample)  59.2433% (out-sample)
W =
        cadd8                    other
------  -----------------------  -----------------------
1e-08   0.001 (-0.1544, 0.1557)  0.995 (0.9888, 1.0016)
1e-07   0.002 (-0.1762, 0.1804)  0.000 (-0.0061, 0.0061)
1e-06   0.000 (-0.0102, 0.0108)  0.000 (-0.0001, 0.0001)
1e-05   0.000 (-0.0002, 0.0003)  0.000 (-0.0000, 0.0000)
0.0001  0.068 (0.0475, 0.0880)   0.000 (-0.0000, 0.0000)
0.001   0.365 (0.2695, 0.4596)   0.000 (-0.0000, 0.0000)
0.01    0.565 (0.4218, 0.7074)   0.005 (0.0034, 0.0062)
```

#### CHB

```
ML estimates: (whole genome)
number jackknife samples: 214
standard error method: moving block jackknife
negative log-likelihood: 61827005013.645355
number of successful starts: 10000 (100.0% total)
pi_0 = 0.00097036 (0.00097, 0.00097)
pi  = 0.000752601
mut. rate = 1.204e-08  (9.22e-09, 1.49e-08)
Ne = 24,259 (if mu=1e-8), Ne = 12,129 (if mu=2e-8)
R2 = 59.6862% (in-sample)  57.7389% (out-sample)
W =
        cadd8                    other
------  -----------------------  -----------------------
1e-08   0.000 (-0.1398, 0.1399)  0.990 (0.9834, 0.9965)
1e-07   0.000 (-0.1330, 0.1331)  0.000 (-0.0061, 0.0061)
1e-06   0.000 (-0.0085, 0.0085)  0.000 (-0.0001, 0.0001)
1e-05   0.000 (-0.0002, 0.0002)  0.000 (-0.0000, 0.0000)
0.0001  0.040 (0.0316, 0.0492)   0.000 (-0.0000, 0.0000)
0.001   0.504 (0.3939, 0.6132)   0.000 (-0.0000, 0.0000)
0.01    0.456 (0.3568, 0.5552)   0.010 (0.0079, 0.0122)
```

#### YRI

```
ML estimates: (whole genome)
number jackknife samples: 263
standard error method: moving block jackknife
negative log-likelihood: 250687699969.473
number of successful starts: 10000 (100.0% total)
pi_0 = 0.00135876 (0.00136, 0.00136)
pi  = 0.00106447
mut. rate = 1.126e-08  (7.74e-09, 1.48e-08)
Ne = 33,968 (if mu=1e-8), Ne = 16,984 (if mu=2e-8)
R2 = 65.4089% (in-sample)  65.1251% (out-sample)
W =
        cadd8                    other
------  -----------------------  -----------------------
1e-08   0.013 (-0.2171, 0.2439)  0.989 (0.9817, 0.9956)
1e-07   0.000 (-0.1466, 0.1466)  0.000 (-0.0059, 0.0059)
1e-06   0.000 (-0.0058, 0.0058)  0.000 (-0.0001, 0.0001)
1e-05   0.000 (-0.0002, 0.0002)  0.000 (-0.0000, 0.0000)
0.0001  0.136 (0.0944, 0.1781)   0.000 (-0.0000, 0.0000)
0.001   0.108 (0.0673, 0.1492)   0.000 (-0.0000, 0.0000)
0.01    0.742 (0.5313, 0.9529)   0.011 (0.0079, 0.0148)
```

#### 6.14 CADD 8%, Sparse Track

##### CEU

```
ML estimates: (whole genome)
standard error method: None
negative log-likelihood: 197849464223.25174
number of successful starts: 10000 (100.0% total)
pi_0 = 0.00101859
pi  = 0.000801495
mut. rate = 1.255e-08
Ne = 25,464 (if mu=1e-8), Ne = 12,732 (if mu=2e-8)
R2 = 60.9207% (in-sample)  59.3772% (out-sample)
W =
          cadd8
------  -------
1e-08     0
1e-07     0
1e-06     0
1e-05     0
0.0001    0.063
0.001     0.354
0.01      0.583
```

##### CHB

```
ML estimates: (whole genome)
standard error method: None
negative log-likelihood: 61827132377.38223
number of successful starts: 10000 (100.0% total)
pi_0 = 0.000963444
pi  = 0.000752601
mut. rate = 1.309e-08
Ne = 24,086 (if mu=1e-8), Ne = 12,043 (if mu=2e-8)
R2 = 59.7755% (in-sample)  57.9086% (out-sample)
W =
          cadd8
------  -------
1e-08     0
1e-07     0
1e-06     0
1e-05     0
0.0001    0.032
0.001     0.469
0.01      0.499
```

#### YRI

```
ML estimates: (whole genome)
standard error method: None
negative log-likelihood: 250688310460.67032
number of successful starts: 10000 (100.0% total)
pi_0 = 0.00134845
pi  = 0.00106447
mut. rate = 1.223e-08
Ne = 33,711 (if mu=1e-8), Ne = 16,855 (if mu=2e-8)
R2 = 65.5761% (in-sample)  65.3002% (out-sample)
W =
          cadd8
------  -------
1e-08     0
1e-07     0
1e-06     0
1e-05     0
0.0001    0.12
0.001     0.106
0.01      0.774
```

### References

1000 Genomes Project Consortium et al. (2015). "A global reference for human genetic variation". en. In: *Nature* 526.7571, pp. 68–74.

Adrion, Jeffrey R et al. (2020). "A community-maintained standard library of population genetic models". en. In: *Elife* 9.

Agarwal, Ipsita, Zachary L Fuller, Simon R Myers, and Molly Przeworski (2023). "Relating pathogenic loss-of-function mutations in humans to their evolutionary fitness costs". en. In: *Elife* 12.

Barton, N H (1986). "The maintenance of polygenic variation through a balance between mutation and stabilizing selection". In: *Genet. Res.* 47.3, pp. 209–216.

— (2000). "Genetic hitchhiking". en. In: *Philos. Trans. R. Soc. Lond. B Biol. Sci.* 355.1403, pp. 1553–1562.

Buffalo, Vince and Graham Coop (2019). "The Linked Selection Signature of Rapid Adaptation in Temporal Genomic Data". en. In: *Genetics* 213.3, pp. 1007–1045.

Byrska-Bishop, Marta et al. (2022). "High-coverage whole-genome sequencing of the expanded 1000 Genomes Project cohort including 602 trios". en. In: *Cell* 185.18, 3426–3440.e19.

Charlesworth, B (1987). *The heritability of fitness.* Sexual selection: testing the alternatives.

Crow, James F (1958). "Some possibilities for measuring selection intensities in man". en. In: *Hum. Biol.* 30.1, pp. 1–13.

Cunningham, Fiona et al. (2022). "Ensembl 2022". en. In: *Nucleic Acids Res.* 50.D1, pp. D988–D995.

Durrett, Richard (2008). *Probability models for DNA sequence evolution.* Springer Science & Business Media.

Elyashiv, Eyal et al. (2016). "A Genomic Map of the Effects of Linked Selection in Drosophila". en. In: *PLoS Genet.* 12.8, e1006130.

García-Dorado, Aurora (2007). "Shortcut predictions for fitness properties at the mutation-selection-drift balance and for its buildup after size reduction under different management strategies". en. In: *Genetics* 176.2, pp. 983–997.

Gessler, D D (1995). "The constraints of finite size in asexual populations and the rate of the ratchet". en. In: *Genet. Res.* 66.3, pp. 241–253.

Good, Benjamin H and Michael M Desai (2013). "Fluctuations in fitness distributions and the effects of weak linked selection on sequence evolution". en. In: *Theor. Popul. Biol.* 85, pp. 86–102.

Green, Melville S (1954). "Markoff random processes and the statistical mechanics of time-dependent phenomena. II. Irreversible processes in fluids". In: *J. Chem. Phys.* 22.3, pp. 398–413.

Gutenkunst, Ryan N, Ryan D Hernandez, Scott H Williamson, and Carlos D Bustamante (2009). "Inferring the joint demographic history of multiple populations from multidimensional SNP frequency data". en. In: *PLoS Genet.* 5.10, e1000695.

Haigh, John (1978). "The accumulation of deleterious genes in a population—Muller's Ratchet". In: *Theor. Popul. Biol.* 14.2, pp. 251–267.

Haller, Benjamin C and Philipp W Messer (2019). "SLiM 3: Forward Genetic Simulations Beyond the Wright-Fisher Model". en. In: *Mol. Biol. Evol.* 36.3, pp. 632–637.

— (2023). "SLiM 4: Multispecies Eco-Evolutionary Modeling". en. In: *Am. Nat.* 201.5, E127–E139.

Higgs, Paul G and Glenn Woodcock (1995). "The accumulation of mutations in asexual populations and the structure of genealogical trees in the presence of selection". In: *J. Math. Biol.* 33.7, pp. 677–702.

Houle, D (1992). "Comparing evolvability and variability of quantitative traits". In: *Genetics.*

Hudson, R R and N L Kaplan (1995). "Deleterious background selection with recombination". en. In: *Genetics* 141.4, pp. 1605–1617.

Hudson, Richard R and Norman L Kaplan (1994). "Gene Trees with Background Selection". In: *Non-Neutral Evolution: Theories and Molecular Data.* Ed. by Brian Golding. Boston, MA: Springer US, pp. 140–153.

Illumina, Inc. (2020). *1000 Genomes Phase 3 Reanalysis with DRAGEN 3.5 and 3.7.* `https://registry.opendata.aws/ilmn-dragen-1kgp.`. Accessed: 2021-7-19.

International HapMap Consortium et al. (2007). "A second generation human haplotype map of over 3.1 million SNPs". en. In: *Nature* 449.7164, pp. 851–861.

Johnson, Steven G (2007). *The NLopt nonlinear-optimization package.* `https://github.com/stevengj/nlopt`.

Keightley, P D and W G Hill (1988). "Quantitative genetic variability maintained by mutation-stabilizing selection balance in finite populations". en. In: *Genet. Res.* 52.1, pp. 33–43.

Kelleher, Jerome, Kevin R Thornton, Jaime Ashander, and Peter L Ralph (2018). "Efficient pedigree recording for fast population genetics simulation". en. In: *PLoS Comput. Biol.* 14.11, e1006581.

Kimura, M (1957). "Some problems of stochastic processes in genetics". In: *Ann. Math. Stat.*

Kimura, Motoo (1969). "The Number of Heterozygous Nucleotide Sites Maintained in a Finite Population Due to Steady Flux of Mutations". In: *Genetics* 61.4, pp. 893–903.

Köster, Johannes and Sven Rahmann (2012). "Snakemake–a scalable bioinformatics workflow engine". en. In: *Bioinformatics* 28.19, pp. 2520–2522.

Kubo, Ryogo (1957). "Statistical-Mechanical Theory of Irreversible Processes. I. General Theory and Simple Applications to Magnetic and Conduction Problems". In: *J. Phys. Soc. Jpn.* 12.6, pp. 570–586.

McVicker, Graham, David Gordon, Colleen Davis, and Phil Green (2009). "Widespread genomic signatures of natural selection in hominid evolution". en. In: *PLoS Genet.* 5.5, e1000471.

Morgulis, Aleksandr, E Michael Gertz, Alejandro A Schäffer, and Richa Agarwala (2006). "A fast and symmetric DUST implementation to mask low-complexity DNA sequences". en. In: *J. Comput. Biol.* 13.5, pp. 1028–1040.

Murphy, David A, Eyal Elyashiv, Guy Amster, and Guy Sella (2022). "Broad-scale variation in human genetic diversity levels is predicted by purifying selection on coding and non-coding elements". In: *Elife* 11, e76065.

Nicolaisen, Lauren E and Michael M Desai (2013). "Distortions in Genealogies due to Purifying Selection and Recombination". en. In: *Genetics* 195.1, pp. 221–230.

Nordborg, Magnus, Brian Charlesworth, and Deborah Charlesworth (1996). "The effect of recombination on background selection*". In: *Genet. Res.* 67.02, pp. 159–174.

O'Fallon, Brendan D, Jon Seger, and Frederick R Adler (2010). "A continuous-state coalescent and the impact of weak selection on the structure of gene genealogies". en. In: *Mol. Biol. Evol.* 27.5, pp. 1162–1172.

Powell, M J D (2009). *The BOBYQA algorithm for bound constrained optimization without derivatives.* Tech. rep. Cambridge, UK: Department of Applied Mathematics and Theoretical Physics, Cambridge University.

Price, G R (1970). "Selection and covariance". In: *Nature.*

Robertson, Alan (1966). "A mathematical model of the culling process in dairy cattle". In: *Anim. Sci.* 8.1, pp. 95–108.

Santiago, E and A Caballero (1995). "Effective size of populations under selection". en. In: *Genetics* 139.2, pp. 1013–1030.

Santiago, E and A Caballero (1998). “Effective size and polymorphism of linked neutral loci in populations under directional selection”. en. In: *Genetics* 149.4, pp. 2105–2117.

Santiago, Enrique and Armando Caballero (2016). “Joint Prediction of the Effective Population Size and the Rate of Fixation of Deleterious Mutations”. en. In: *Genetics* 204.3, pp. 1267–1279.

Schrider, Daniel R and Andrew D Kern (2017). “Soft Sweeps Are the Dominant Mode of Adaptation in the Human Genome”. en. In: *Mol. Biol. Evol.* 34.8, pp. 1863–1877.

Siepel, Adam et al. (2005). “Evolutionarily conserved elements in vertebrate, insect, worm, and yeast genomes”. en. In: *Genome Res.* 15.8, pp. 1034–1050.

Smit, A F A, R Hubley, and P Green (2015). *RepeatMasker Open-4.0. 2013–2015.*

Tajima, Fumio (1983). “Evolutionary relationship of DNA sequences in finite populations”. In: *Genetics* 105.2, pp. 437–460.

Turelli, Michael and N H Barton (1990). “Dynamics of polygenic characters under selection”. In: *Theor. Popul. Biol.* 38.1, pp. 1–57.

Wakeley, John (2009). *Coalescent Theory: An Introduction.* Roberts and Company Publishers.

Walsh, Bruce and Michael Lynch (2018). *Evolution and Selection of Quantitative Traits.* en. Oxford University Press.

Wasserman, Larry (2004). *All of Statistics.* Springer New York.

Woolliams, J A, N R Wray, and R Thompson (1993). “Prediction of long-term contributions and inbreeding in populations undergoing mass selection”. In: *Genet. Res.* 62.3, pp. 231–242.

Wray, N R and R Thompson (1990). “Prediction of rates of inbreeding in selected populations”. en. In: *Genet. Res.* 55.1, pp. 41–54.

Wright, Sewall (1938). “Size of population and breeding structure in relation to evolution”. In: *Science* 87.2263, pp. 430–431.
